# Identification of synaptic PPT1 substrates highlight roles of depalmitoylation in disulfide bond formation and synaptic function

**DOI:** 10.1101/2020.05.02.074302

**Authors:** Erica L. Gorenberg, Sofia Massaro Tieze, Betül Yücel, Helen R. Zhao, Vicky Chou, Gregory S. Wirak, Susumu Tomita, TuKiet T. Lam, Sreeganga S. Chandra

## Abstract

Loss-of-function mutations in the depalmitoylating enzyme palmitoyl protein thioesterase 1 (PPT1) cause Neuronal Ceroid Lipofuscinosis type 1 (*CLN1*), a devastating neurodegenerative disease. Here, we provide a resource identifying PPT1 substrates. We utilized Acyl Resin-Assisted Capture and mass spectrometry to identify proteins with increased *in vivo* palmitoylation in PPT1 knockout mouse brains. We then validated putative substrates through direct depalmitoylation with recombinant PPT1. This stringent screen elucidated >100 novel PPT1 substrates at the synapse, including channels and transporters, G-protein-associated molecules, endo/exocytic components, synaptic adhesion molecules, and mitochondrial proteins. Cysteine depalmitoylation sites in transmembrane PPT1 substrates frequently participate in disulfide bonds in the mature protein. We confirmed that depalmitoylation regulates disulfide bond formation in a tertiary screen analyzing post-translational modifications. Collectively, the diverse PPT1 substrates highlight the role of PPT1 in mediating synapse functions, implicate molecular pathways in the etiology of *CLN1*, and advance our basic understanding of the purpose of depalmitoylation.

**Highlights:**

- ∼10% of palmitoylated proteins are palmitoyl protein thioesterase 1 (PPT1) substrates
- Unbiased proteomic approaches identify 9 distinct classes of PPT1 substrates, including synaptic adhesion molecules and endocytic proteins
- Protein degradation does not require depalmitoylation by PPT1
- Depalmitoylation mediates disulfide bond formation in transmembrane PPT1 substrates

## BACKGROUND

Palmitoylation is the post-translational addition of a C16:0 fatty acid chain to proteins, typically via thioester bond to cysteine residues. Palmitoylation is unique amongst lipid post-translational modifications (PTMs) in its reversibility—other lipid PTMs typically last a protein’s entire lifespan (Magee et al., 1987). Palmitoyl groups are added by a family of 23 palmitoyl acyl transferases (PATs or ZDHHCs; Lemonidis et al., 2015; Mitchell et al., 2006), and are removed by distinct depalmitoylating enzymes – palmitoyl protein thioesterases (PPTs), acyl protein thioesterases (APTs), and α/β-hydrolase domain-containing proteins (ABHDs; Lin and Conibear, 2015; Vartak et al., 2014; Vesa et al., 1995). The reversible nature of palmitate PTMs suggests that tight regulation of the palmitoylation/depalmitoylation cycle is necessary for proper protein function. Indeed, palmitoylation dynamics influence protein stability, function, membrane trafficking, and subcellular localization (Greaves and Chamberlain, 2007; Greaves et al., 2009; Hayashi et al., 2005; Lin et al., 2009).

Systematic and unbiased proteomic studies of the mammalian brain have identified over 600 palmitoylated proteins, a high percentage of which are localized to synapses (Collins et al., 2017; Kang et al., 2008; Segal-Salto et al., 2017). These large-scale screens suggest the importance of palmitoylation for synaptic function and neurotransmission. Recent studies have begun to identify the synaptic substrates of specific PATs (Fukata et al., 2004; Huang et al., 2004; Collura et al., 2020; Sanders et al., 2020). However, it is still unclear for most palmitoylated proteins which enzyme facilitates depalmitoylation in the brain and at the synapse.

Loss-of-function mutations in the depalmitoylating enzyme PPT1 lead to deficient depalmitoylation and synaptic function, and result in the neurodegenerative disease *CLN1* (Mitchison et al., 1998; Vesa et al., 1995). PPT1 therefore plays a critical role in synaptic function. However, the physiological substrates of PPT1 are almost entirely unknown, with the exceptions of the presynaptic chaperone CSPα, the G-protein Goα, and the mitochondrial F1 ATP synthase subunit O (Bagh et al., 2017; Duncan and Gilman, 1998; Lyly et al., 2007). Identification of the repertoire of PPT1 substrates is needed to understand how depalmitoylation deficits impact neuronal health.

In this study, we therefore undertook a two-step proteomic approach to identify PPT1 substrates and elucidate how depalmitoylation contributes to synaptic function and phenotypes of NCL. Our screen greatly expands the repertoire of known palmitoylated proteins in the brain. We detected PPT1 in all synaptic sub-compartments and identified ∼10% of palmitoylated synaptic proteins as PPT1 substrates. These proteins display increased palmitoylation independently of protein expression *in vivo*. The validated substrates fall into nine classes, which, strikingly, are related to phenotypes observed in PPT1 KO mice and *CLN1* patients, including seizures, decreased synapse density, mitochondrial dysfunction, synaptic vesicle endocytic deficits, impaired long-term potentiation (LTP), and retinal degeneration (Gupta et al., 2001; Kim et al., 2008; Sapir et al., 2019; Virmani et al., 2005; Wei et al., 2011; Friedrich et al., 2011). Notably, PPT1 depalmitoylation sites on validated substrates are frequently cysteine residues that participate in disulfide bonds, suggesting that a novel function of palmitoylation is to mediate these interactions. Our classification of PPT1 substrates provides a resource to enhance our understanding of depalmitoylation and its contributions to synaptic function, neuronal health, and the molecular basis of NCL.

## RESULTS

### WT and PPT1 KO brain palmitome expands the repertoire of known palmitoylated proteins in the brain and draws links between NCLs

Neuronal substrates of PPT1 are expected to exhibit increased palmitoylation in PPT1 KO mouse brains, but it is unclear whether these changes are independent of protein levels. Therefore, we designed an experimental workflow to analyze both the proteome and palmitome from the same WT and PPT1 KO brains by Label-Free Quantification Mass Spectrometry (LFQ-MS; **Figure 1A**). This scheme allowed for comparisons across both genotypes (WT vs PPT1 KO) and ‘-omes’ (proteome vs. palmitome) and utilized *in vivo* palmitoylation changes to identify PPT1 substrates. Synaptosomes and whole brain homogenates from WT and PPT1 KO mice were processed for Acyl Resin-Assisted Capture (Acyl RAC), as previously described (Forrester et al., 2010; Henderson et al., 2016). Briefly, disulfide bonds were reduced with tris(2-carboxyethyl)phosphine (TCEP) and resulting free thiols were blocked with N-ethylmaleimide (NEM; **Figure 1Bi**).

**Figure 1.**
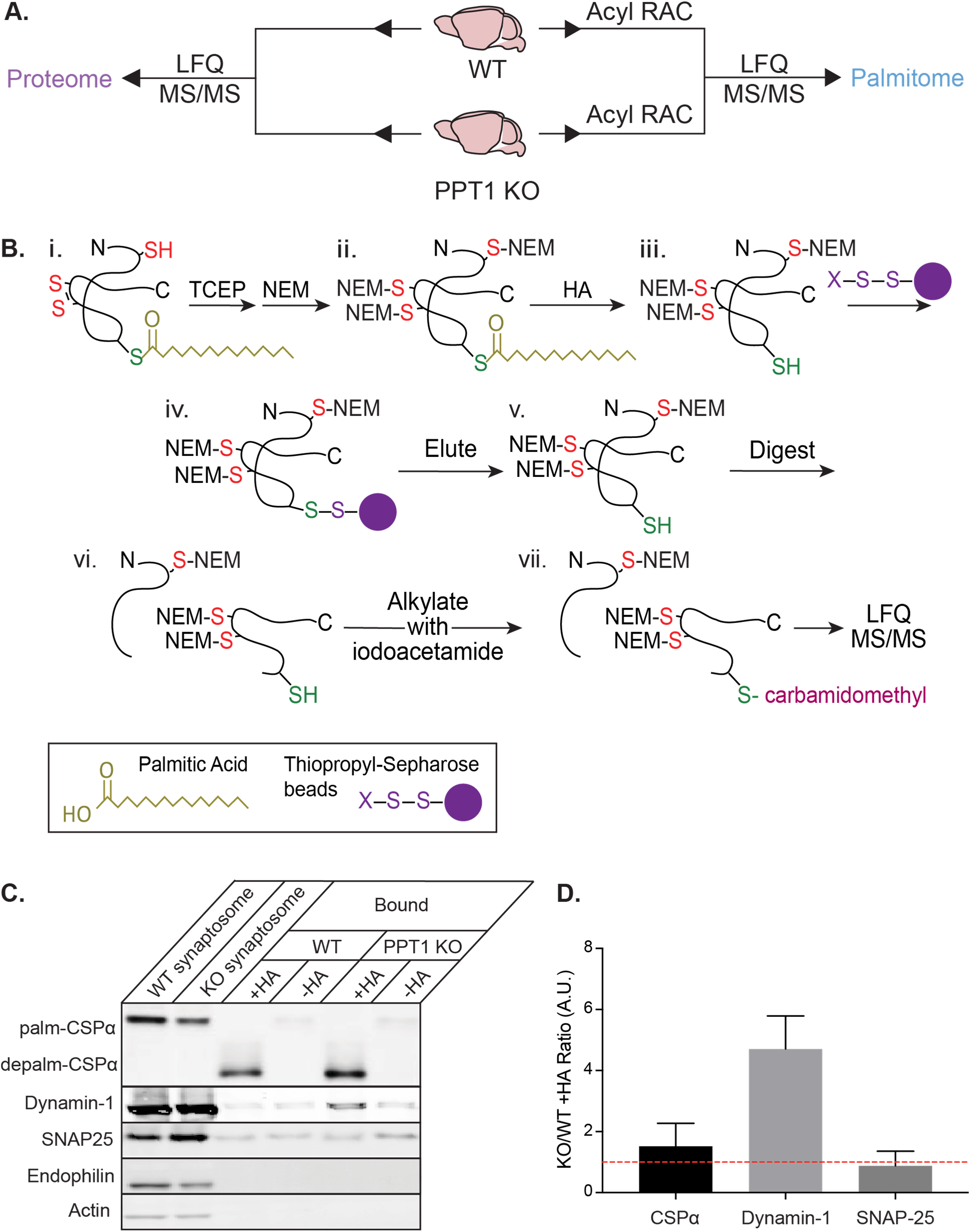
Generation of the proteome and palmitome from WT and PPT1 KO brains. A) Proteins were isolated from WT and PPT1 KO whole brains or synaptosomes for proteomic analysis by Label-Free Quantification Mass Spectrometry (LFQ MS/MS) to identify overall protein expression (proteome; left) or following Acyl Resin-Assisted Capture (Acyl RAC) to identify palmitoylated proteins (palmitome; right). B) Schematic of Acyl RAC. i) Disulfide bonds were reduced with tris(2-carboxyethyl)phosphine (TCEP) and residual thiol groups were blocked with N-ethylmaleimide (NEM). ii) Palmitate groups were hydrolyzed with hydroxylamine (HA), leaving behind an unblocked free thiol. iii) Free thiols were bound to thiopropyl sepharose beads, allowing for isolation of palmitoylated proteins. These proteins were then (iv) eluted from beads with DTT, (v) digested, and (vi) alkylated with iodoacetamide, followed by (vii) LFQ MS/MS analysis. C) Pull-down and immunoblotting of palmitoylated proteins. Addition of HA allowed CSPα, dynamin-1, and SNAP-25, all of which are palmitoylated proteins, to be pulled down on thiopropyl sepharose beads, but not endophilin or actin, which are not palmitoylated. Note that addition of HA removes palmitate and causes a molecular weight shift of palmitoylated CSPα to depalmitoylated CSPα. D) Quantification of western blot +HA band intensity (A.U), plotted as a ratio of KO/WT. CSPα is a known PPT1 substrate and shows elevated palmitoylation in KO/WT, while SNAP-25 does not (n = 3 experiments).

Palmitate groups were removed with hydroxylamine (HA; **Figure 1Bii**) and resulting unblocked free thiols were bound to thiopropyl sepharose beads (**Figure 1Biii**). Bound proteins were then eluted for Liquid Chromatography-Mass Spectrometry (LC-MS/MS; **Figure 1Biv**), selectively isolating previously palmitoylated proteins (**Figure 1C-D**). Proteome fractions were collected prior to depalmitoylation by HA (**Figure 1A, Bii**), while palmitome fractions were isolated following pulldown on and elution from thiopropyl sepharose beads (**Figure 1Biv**). Both the proteome and palmitome were analyzed by Label-Free Quantification-MS (LFQ-MS) following digestion and alkylation with iodoacetamide to modify previously-palmitoylated cysteines with a carbamidomethyl moiety (**Figure 1Bv-vii**).

The Acyl RAC workflow was conducted on WT and PPT1 KO mouse brains (n = 3, each in technical triplicate; age = 2 months) in a pair-wise fashion. We examined proteins common to all three experiments, cognizant of the fact that this stringency is likely to result in underestimation of the number of PPT1 substrates. The resulting MS analysis of the palmitome identified 1795 common palmitoylated proteins (**Figure S1A-B**), and analysis of the proteome identified 1873 common proteins (**Figure S1C-D**) between genotypes. Most of the proteins identified in the brain palmitome (69.8%; **Figure S1A**) were previously validated as palmitoylated or identified in palmitoylation datasets (SwissPalm; Blanc et al., 2015), which confirms our ability to successfully identify palmitoylated proteins using Acyl RAC. This analysis expands the number of previously known palmitoylated proteins in the brain (Collins et al., 2017; Kang et al., 2008; Segal-Salto et al., 2017).

Palmitoylated protein levels in the brain palmitome are highly correlated across genotypes (m = 0.9753 ± 0.0030; R^2^ = 0.9857; **Figure S1E**) and only 12 proteins display significantly increased expression (**Figure S1B**). As anticipated, PPT1 levels are significantly decreased in the PPT1 KO; detection of PPT1 may be explained by sequence similarity with PPT2 or translation of a truncated, catalytically inactive N-terminal PPT1 transcript (Gupta et al., 2001). Protein expression levels in the brain proteome were also highly correlated between WT and PPT1 KO brains (m = 0.9802 ± 0.0023; R^2^ = 0.9916; **Figure S1F**), in line with the lack of overt differences in neurological phenotypes at this age (2 months; Gupta et al., 2001).

We inspected the status of other known depalmitoylating enzymes – PPT2, APTs, and ABHDs – and found their levels unaltered, suggesting they do not display compensatory upregulation of protein expression due to loss of PPT1 (ABHD12, APT1, APT2; **Figure S1D**, green points). We find increased levels of ASAH1 (Acid Ceramidase), CATD (Cathepsin D), SCRB2 (Lysosome Membrane Protein 2), SAP (Prosaposin), and TPP1 (Tripeptidyl Peptidase 1) in the PPT1 KO palmitome (**Figure S1B**) and proteome (**Figure S1D**), while the remaining proteins with elevated palmitoylation—C1QC, PTTG, CATF, AP1B1, SRBS1, SRBS2, and RTN3—do not display concurrent elevation of protein expression. These five proteins were previously shown to be increased in PPT1 KO or *CLN1* models (Chandra et al., 2015; Sleat et al., 2017; Tyynela et al., 1993). Intriguingly, TPP1 and CATD loss-of-function mutations cause other forms of NCL (*CLN2* and *CLN10*, respectively) and isoforms of SAP accrue in *CLN1* brains (Tyynela et al., 1993), suggesting a common etiological pathway.

Overall, 72% of palmitoylated proteins (1290/1795) were also identified in the proteome sample (**Figure S2G**), allowing direct comparisons to distinguish increased palmitoylation from increased protein expression. For proteins with significantly increased expression in the palmitome (**Figure S2B**), we plotted palmitome versus proteome expression levels in PPT1 KO after normalization to WT (**Figure S2H**). We found that the ratios for the lysosomal proteins ASAH1, CATD, SCRB2, SAP, and TPP1 were unchanged or decreased, while those for AP1B1, SRBS1, SRBS2, and RTN3 show elevated palmitoylation relative to protein expression in PPT1 KO brains, indicating that the latter proteins can be considered putative PPT1 substrates (**Figure S2H**). In further support of this conclusion, all putative substrates but SRBS2 were previously shown to be palmitoylated (SwissPalm; Blanc et al., 2015).

### Putative PPT1 substrates are enriched at the synapse

Immunocytochemistry demonstrates that PPT1 is detected at synapses when expressed in neurons (**Figure S2A**), consistent with previous literature, while it is largely lysosomal in non-neuronal cell types (Ahtiainen et al., 2002; Kim et al., 2008; Lehtovirta et al., 2001). We confirmed the presence of endogenous PPT1 at the synapse by immunoblotting subcellular fractionation samples generated from WT and PPT1 KO mouse brains (age = 2 months). PPT1 is detected in all synaptic fractions in WT and is absent in PPT1 KO **(Figure S2B)**. As we identified very few palmitoylation changes between WT and PPT1 KO from the brain analyses, including changes to known substrates of PPT1—CSPα, Goα, and F1 ATPase subunit O (Bagh et al., 2017; Duncan and Gilman, 1998; Lyly et al., 2007)—we reasoned that sub-compartmentalization of PPT1 may have diluted the LFQ signal. Therefore, we proceeded to isolate synaptosomes (a biochemical synaptic preparation) from WT and PPT1 KO brains to enrich for PPT1 and its potential synaptic substrates. We then subjected synaptosomes to the same Acyl RAC workflow (**Figure 1B**). We identified high degrees of overlap between the 3 synaptosomal preparations for the palmitome (1379 common proteins; **Figure 2A-B**), and the proteome (1826 common proteins; **Figure 2C-D**) indicating excellent reproducibility,

**Figure 2.**
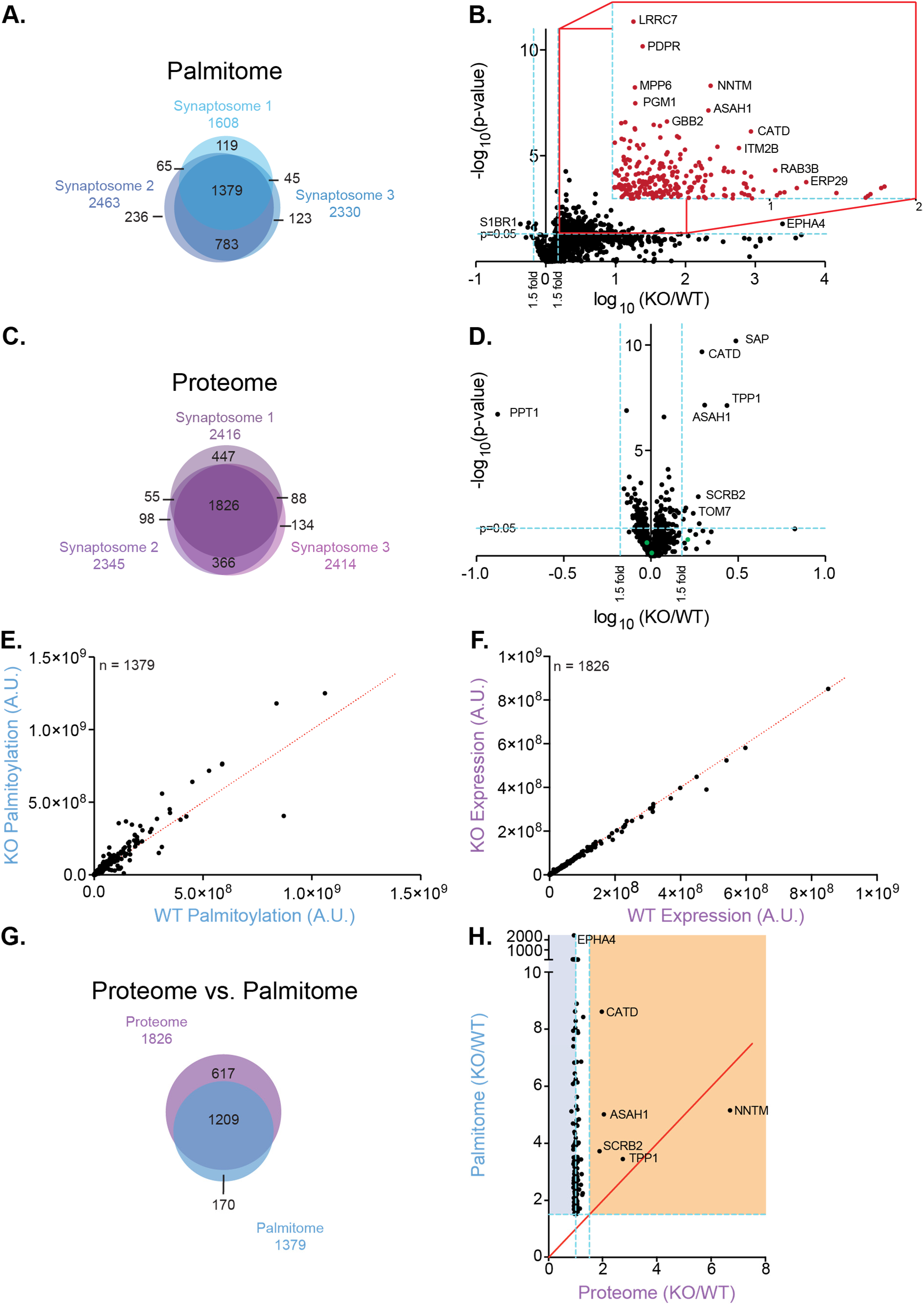
Putative PPT1 substrates are enriched at the synapse and display persistent palmitoylation. A) Venn diagram of 3 palmitome experiments exhibits 1379 common proteins. B) Volcano plot indicates 242 proteins are significantly differentially palmitoylated (4 decrease; 238 increase; 1.5-fold, p < 0.05; blue lines; **Table S2**). Inset shows significantly increased proteins. The p-value was calculated using a two-tailed t-test on 3 biological and 3 technical replicates. While the technical replicates do not meet the t-test condition for independence, we chose to proceed in this manner to generate more putative substrates that could later be validated with higher stringency. C) Venn diagram of 3 independent proteome experiments exhibits 1826 common proteins. D) Volcano plot of fold change between genotypes (KO/WT) for proteins identified in all 3 biological replicates. 11 proteins are significantly differentially expressed (1 decrease (PPT1); 10 increase; 1.5-fold, p < 0.05; blue lines). Other depalmitoylating enzymes (green points) do not display significant compensatory upregulation of protein expression. E) Palmitoylated protein expression is somewhat correlated between genotypes (R^2^ = 0.899; m = 1.134 ± 0.0197), with increased palmitoylation in PPT1 KO. F) Protein expression is almost perfectly correlated between genotypes (R^2^ = 0.9966; m = 0.969 ± 0.0016). Red lines indicate 1:1 WT to PPT1 KO protein expression ratio. G) Venn diagram of 1209 common proteins between whole brain proteome and palmitome (n=3 each). H) Protein expression levels compared to palmitoylation levels for significantly changed proteins in palmitome. Proteins in orange region are increased (1.5-fold, p < 0.05) in both proteome and palmitome. Proteins in blue region display decreased or unchanged protein expression and increased palmitoylation. Red line indicates equal expression and palmitoylation levels (x = y). While NNTM has a high fold change, it does not meet the p-value criterion for significantly increased protein expression (p = 0.0522) and is therefore excluded from this category.

When we compared synaptic proteomes across genotypes, we corroborated that protein expression between WT and PPT1 KO is highly correlated (m = 0.969 ± 0.0016; R^2^ = 0.9666; **Figure 2F**). Consistent with a synaptic localization and previous literature, we identified PPT1 in the synaptosomal proteome (**Table S1**; Ahtiainen et al., 2002; Kim et al., 2008; Lehtovirta et al., 2001). Notably, PPT1 is the only protein with significantly decreased levels in PPT1 KO synaptosomes, and only 10 proteins exhibited significantly increased expression (proteome KO/WT ratio >1.5; p <0.05; **Figure 2D**). The proteins with increased expression were also identified in the whole brain analysis (**Figure S1**), with the exceptions of the mitochondrial proteins TOM7, CX7A2, and NDUF4. As with the whole brain, we also identified other depalmitoylating enzymes in the synaptic proteome and found their levels to be unaltered (ABHD12, ABHDA, ABHGA, APT2; **Figure 2D**, green points).

Notably, there was a much lower correlation between WT and PPT1 KO palmitomes (m = 1.134 ± 0.0197; R^2^ = 0.8986; **Figure 2E**), with many (610; 44.3%) points above the y=x unity line indicating elevated palmitoylation in PPT1 KO synaptosomes. The remarkable increase in palmitoylation in PPT1 KO synaptosomes (**Figure 2B**) compared to whole brain (**Figure S1B**), supports PPT1-mediated depalmitoylation of diverse synaptic substrates, as a depalmitoylation deficiency is expected to lead to an accumulation of palmitoylated targets.

Indeed, the proteins of greatest interest as potential PPT1 substrates were those with significantly increased palmitoylation in PPT1 KO compared to WT (p <0.05, 1.5-fold change cut-off). We identified 227 such proteins (**Figure 2B**; **Table S2)** that were considered putative PPT1 substrates and include the previously known substrates, CSPα, F1 ATP synthase subunit O, and Goα (Bagh et al., 2017; Duncan and Gilman, 1998; Lyly et al., 2007). Most of the putative substrates were also identified in the whole brain proteome (77.1%; blue points; **Figure S1D**;) and palmitome (72.0%; blue points; **Figure S1B**), but their palmitoylation levels were not significantly altered, likely due to the synaptic compartmentalization of PPT1.

Approximately 66% of proteins present in the synaptic palmitome were also identified in the synaptic proteome (n = 1209; **Figure 2G**). To confirm that the increased levels of putative PPT1 substrates in the palmitome are not due to elevated protein expression, we plotted the KO/WT ratio to elucidate the relationship between palmitoylation and protein levels (**Figure 2H**). In line with our initial analyses (**Figure 2B** and **D**), we found only 4 proteins with increased expression and increased palmitoylation in PPT1 KO synaptosomes: ASAH1, CATD, SCRB2, and TPP1 (**Figure 3F**; orange region). For this group of proteins, we could not adequately discriminate increases in palmitoylation resulting from increased protein expression versus PPT1 deficiency. Hence, these proteins may exhibit depalmitoylation-dependent degradation or may not be PPT1 substrates. Therefore, these 4 proteins were removed from subsequent analyses, along with 19 additional proteins which were not identified in the proteome, as we could not confirm whether their increased palmitoylation was independent of protein expression level (**Table S2**).

**Figure 3.**
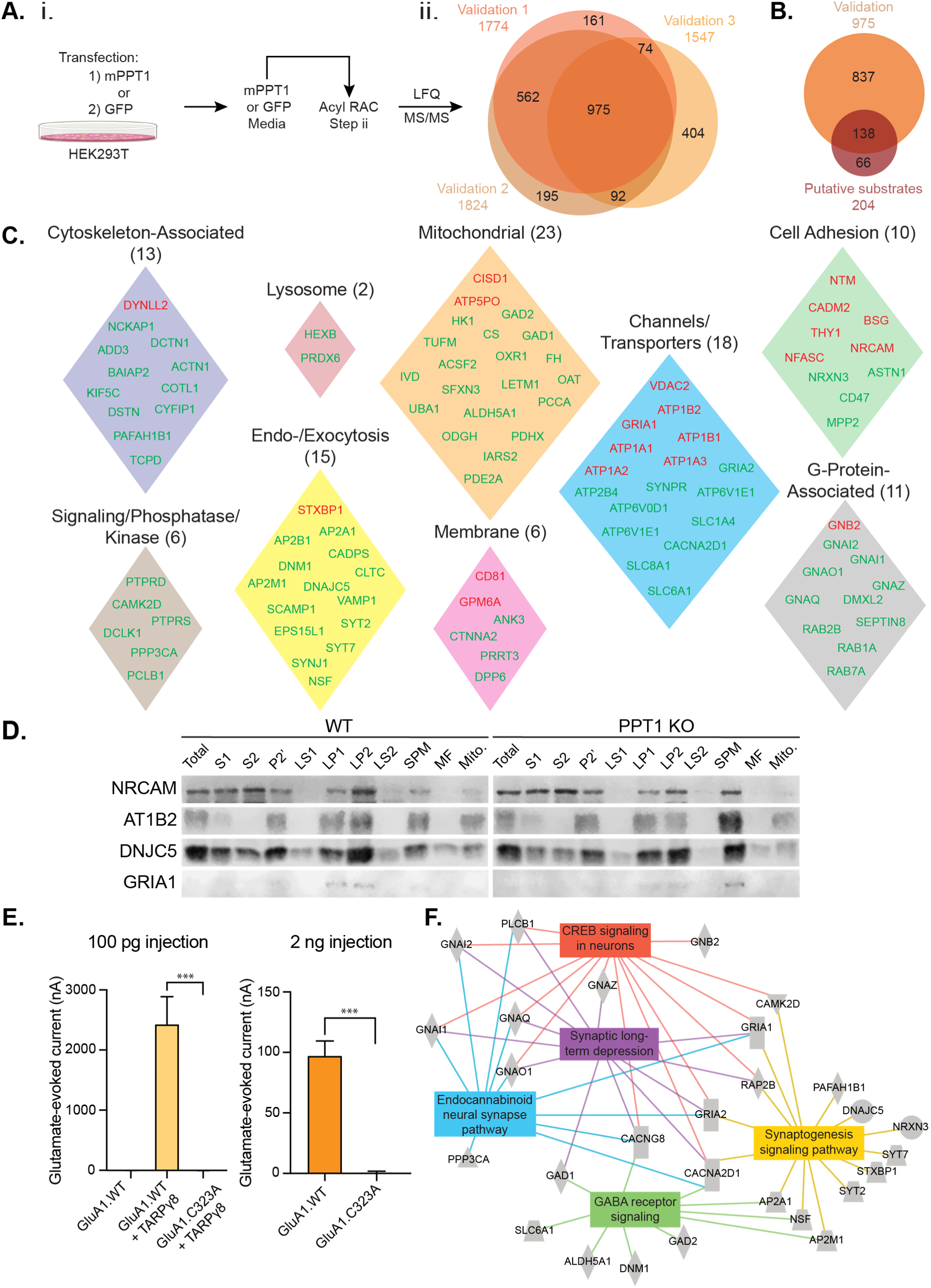
Direct depalmitoylation with PPT1 validates putative synaptic substrates. A) i. Schematic of PPT1-mediated depalmitoylation assay, in which solubilized synaptosomes are treated with recombinant mPPT1 or GFP generated from HEK293T cells during Acyl RAC (Figure 1Bii). ii. Venn diagram of PPT1-mediated depalmitoylation assay (n = 3 biological replicates, n = 3 technical replicates) shows 975 proteins common to replicates. B) Comparison of PPT1-mediated depalmitoylation proteins to putative substrates results in 138 validated PPT1 substrates. C) The 138 combined high- (red) and medium- (green) confidence validated proteins fall into curated UniProt functional and subcellular location groupings. D) Western blots of subcellular fractionation samples prepared from whole brains of WT and PPT1 KO mice. E) *Xenopus laevis o*ocytes were injected with wild type GluA1 (GluA1.WT) or C323A mutant GluA1 cRNA (100 pg) alone or with auxiliary factor TARPγ8 (100 pg). Glutamate (5 μm)-evoked current with cyclothiazide (50 μM) was measured by two-electrode voltage-clamp recording (n = 4; *** p < 0.001). GluA1.WT (2 ng) or GluA1.C323 (2 ng) were injected at higher concentrations to test spontaneous channel formation in the absence of auxiliary factors and glutamate (1mM)-evoked current with cyclothiazide (50 μm) was recorded (n = 6; *** p < 0.001). Data are presented as mean ± SEM. F) Ingenuity pathway analysis displays enrichment of high- and medium-confidence substrates for synaptic long-term depression, synaptogenesis signaling, GABA receptor signaling, endocannabinoid neural synapses, and CREB signaling in neurons.

The remaining putative substrates identified in both proteome and palmitome (n = 204; blue/white region, **Figure 2H**) showed elevated palmitoylation without a commensurate increase in protein levels. Significantly, we identified the putative substrate dynamin-1 in this subset, which showed increased Acyl RAC pulldown in the PPT1 KO synaptosomes by Western blotting (**Figure 1C-D**) but not SNAP-25, congruent with these results. The relative increase in palmitoylation in PPT1 KO ranged from 1.5-fold at the lower limit to 2400-fold (Ephrin A4; EphA4) at the upper limit, with known PPT1 substrates CSPα (DNJC5; 5.12-fold), F1 ATPase subunit O (ATPO; 3.39-fold), and the Goα subunit (GNAO; 4.81-fold) at 3-5-fold (**Figure 2H**).

Finally, we examined the remaining putative PPT1 substrates biochemically for evidence of palmitoylation. By analyzing individual peptides identified by LFQ-MS for each protein, we identified cysteine-containing peptides that were originally palmitoylated by their now-derivatized carbamidomethyl moiety. Out of the 204 putative substrates, we detected a specific palmitoylation site for 101 proteins (49.5%; **Table S3**).

### Direct depalmitoylation with PPT1 validates putative synaptic substrates

To validate the 204 putative PPT1 substrates identified in our initial Acyl RAC screen, we developed a modified secondary Acyl RAC screen in which recombinant mouse PPT1 expressed in HEK293T cells is used as the depalmitoylating reagent, rather than hydroxylamine. We previously established that recombinant PPT1 can depalmitoylate its substrate CSPα *in vitro* (Henderson et al., 2016). Therefore, mouse PPT1 was incubated with WT synaptosomes at Step ii of the Acyl RAC protocol (**Figure 1Bii, Figure 3Ai**).

MS analysis of PPT1-depalmitoylated samples identified 975 common proteins over 3 biological replicates (**Figure 3Aii)**. 138 of the 204 putative substrates identified in the primary screen were validated (**Figure 3B**). Significantly, investigation of the peptide data allowed us to identify the carbamidomethyl-modified peptide, and thus the specific palmitoylated cysteine residue, for a subset of these proteins, as described above. We used this information to define degrees of stringency for the categorization of protein hits from the validation screen, as follows: 1) High-confidence PPT1 substrates are those for which matching carbamidomethyl peptides were identified in both the initial Acyl RAC WT/PPT1 KO screen and the PPT1-mediated depalmitoylation validation screen (n = 26; **Table 1**). 2) Medium-confidence substrates are those identified in both the primary and validation screens with ≥2 unique peptides per protein and a confidence score >100 (n = 112; **Table 2**). 3) Residuals were proteins identified as putative substrates only in the secondary validation screen (n = 831; **Table S4**).

**Table 1.** High-confidence validated PPT1 substrates. X = validation of given dataset; blank cells are lack of identification in that dataset. SwissPalm score indicates percentage of palmitoylation experiments in which the protein has been identified.

| Protein Name<br>(Uniprot ID) | Gene<br>name | Description | Palmitome<br>Ratio | Palmitome<br>p-value | Proteome<br>Ratio | Proteome<br>p-value | CRAPome<br>(%) | Kang<br>et al. | Swiss Palm<br>(%) | CSS<br>Palm |
| --- | --- | --- | --- | --- | --- | --- | --- | --- | --- | --- |
| AT1A1_MOUSE | *Atp1a1* | Sodium/potassium-transporting ATPase subunit alpha-1 | 1.831 | 0.007 | 0.960 | 0.318 | 10.949 |  | 5.882 | X |
| AT1A2_MOUSE | *Atp1a2* | Sodium/potassium-transporting ATPase subunit alpha-2 | 1.744 | 0.003 | 1.068 | 0.498 | 22.871 |  | 5.882 |  |
| AT1A3_MOUSE | *Atp1a3* | Sodium/potassium-transporting ATPase subunit alpha-3 | 1.746 | 0.030 | 1.042 | 0.695 | 24.574 |  | 5.882 |  |
| AT1B1_MOUSE | *Atp1b1* | Sodium/potassium-transporting ATPase subunit beta-1 | 2.180 | 0.019 | 0.982 | 0.536 | 0.243 | X | 5.882 |  |
| AT1B2_MOUSE | *Atp1b2* | Sodium/potassium-transporting ATPase subunit beta-2 | 6.852 | 0.018 | 1.043 | 0.471 | 0.000 | X | 5.882 | X |
| ATPO_MOUSE | *Atp5po* | ATP synthase subunit O, mitochondrial | 6.138 | 0.044 | 1.060 | 0.277 | 0.000 | X | 5.882 |  |
| BASI_MOUSE | *Bsg* | Basigin | 2.695 | 0.016 | 0.927 | 0.101 | 9.732 | X | 5.882 | X |
| CADM2_MOUSE | *Cadm2* | Cell adhesion molecule 2 | 1.975 | 0.004 | 0.988 | 0.559 | 0.000 |  | 11.76 | X |
| CD81_MOUSE | *Cd81* | CD81 antigen | 2.554 | 0.042 | 1.035 | 0.940 | 0.243 |  | 29.41 |  |
| CISD1_MOUSE | *Cisd1* | CDGSH iron-sulfur domain-containing protein 1 | 1.920 | 0.050 | 0.992 | 0.635 | 1.703 |  | 35.29 |  |
| DYL2_MOUSE | *Dynll2* | Dynein light chain 2, cytoplasmic | 4.192 | 0.027 | 0.996 | 0.808 | 9.246 |  | 0 | X |
| GBB2_MOUSE | *Gnb2* | Guanine nucleotide-binding protein G(I)/G(S)/G(T) subunit beta-2 | 2.963 | 0.003 | 0.985 | 0.636 | 15.329 |  | 2.92 | X |
| GPM6A_MOUSE | *Gpm6a* | Neuronal membrane glycoprotein M6-a | 5.625 | 0.007 | 1.043 | 0.658 | 0.243 | X | 5.882 | X |
| GRIA1_MOUSE | *Gria1* | Glutamate receptor 1 | 1.866 | 0.046 | 0.992 | 0.635 | 0.000 | X | 5.882 | X |
| KCRB_MOUSE | *Ckb* | Creatine kinase B-type | 1.815 | 0.029 | 0.959 | 0.242 | 39.659 |  | 11.76 |  |
| LDHB_MOUSE | *Ldhb* | L-lactate dehydrogenase B chain | 1.928 | 0.050 | 1.006 | 0.939 | 39.903 | X | 11.76 |  |
| LGI1_MOUSE | *Lgi1* | Leucine-rich glioma-inactivated protein 1 | 2.047 | 0.013 | 1.014 | 0.883 | 0.000 | X | 11.76 | X |
| NDKA_MOUSE | *Nme1* | Nucleoside diphosphate kinase A | 5.185 | 0.020 | 1.058 | 0.169 | 29.927 |  | 29.41 |  |
| NDUS1_MOUSE | *Ndufs1* | NADH-ubiquinone oxidoreductase 75 kDa subunit, mitochondrial | 1.859 | 0.005 | 0.967 | 0.283 | 4.623 |  | 29.41 | X |
| NFASC_MOUSE | *Nfasc* | Neurofascin | 1.524 | 0.033 | 1.028 | 0.745 | 0.000 | X | 35.29 | X |
| NRCAM_MOUSE | *Nrcam* | Neuronal cell adhesion molecule | 1.714 | 0.011 | 1.148 | 0.314 | 0.000 |  | 35.29 | X |
| NTRI_MOUSE | *Ntm* | Neurotrimin | 4.060 | 0.033 | 0.988 | 0.574 | 0.000 | X | 47.06 | X |
| STXB1_MOUSE | *Stxbp1* | Syntaxin-binding protein 1 | 1.752 | 0.036 | 1.058 | 0.358 | 0.487 |  | 7.056 | X |
| THY1_MOUSE | *Thy1* | Thy-1 membrane glycoprotein | 3.357 | 0.022 | 1.069 | 0.541 | 0.487 |  | 17.65 |  |
| VDAC2_MOUSE | *Vdac2* | Voltage-dependent anion-selective channel protein 2 | 2.929 | 0.029 | 0.946 | 0.058 | 14.112 | X | 35.29 | X |
| VISL1_MOUSE | *Vsnl1* | Visinin-like protein 1 | 2.451 | 0.040 | 0.912 | 0.039 | 0.243 | X | 41.18 | X |

**Table 2.** Medium-confidence validated PPT1 substrates. X = validation of given dataset; blank cells are lack of identification in that dataset. SwissPalm score indicates percentage of palmitoylation experiments in which the protein has been identified.

| Protein Name<br>Uniprot ID | Gene<br>Name | Description | Palmitome<br>Ratio | Palmitome<br>p-value | Proteome<br>Ratio | Proteome<br>p-value | CRAPome<br>(%) | Kang<br>et al. | Swiss<br>Palm (%) | CSS<br>Palm |
| --- | --- | --- | --- | --- | --- | --- | --- | --- | --- | --- |
| 1433B_MOUSE | *Ywhab* | 14-3-3 protein beta/alpha | 6.1651 | 0.0345 | 0.9858 | 0.5572 |  |  | 0 | X |
| ACON_MOUSE | *Aco2* | Aconitate hydratase, mitochondrial | 2.376 | 0.027 | 1.026 | 0.810 | 6.569 |  | 0 | X |
| ACSF2_MOUSE | *Acsf2* | Medium-chain acyl-CoA ligase ACSF2, mitochondrial | 6.8591 | 0.0431 | 1.1975 | 0.0262 | 0.4866 |  | 0 | X |
| ACTN1_MOUSE | *Actn1* | Alpha-actinin-1 | 2.9415 | 0.0091 | 1.0425 | 0.5896 |  |  | 0 |  |
| ADDG_MOUSE | *Add3* | Gamma-adducin | 2.0263 | 0.0064 | 0.9829 | 0.5518 | 7.2993 |  | 0 |  |
| AIFM1_MOUSE | *Aifm1* | Apoptosis-inducing factor 1, mitochondrial | 2.8165 | 0.0430 | 0.9702 | 0.4086 |  |  | 0 | X |
| AL1L1_MOUSE | *Aldh1l1* | Cytosolic 10-formyltetrahydrofolate dehydrogenase | 3.1219 | 0.0122 | 1.0338 | 0.5529 | 0.9732 |  | 0 | X |
| ANK3_MOUSE | *Ank3* | Ankyrin-3 | 2.4417 | 0.0253 | 1.0168 | 0.8537 | 3.6496 |  | 0.487 |  |
| AP2A1_MOUSE | *Ap2a1* | AP-2 complex subunit alpha-1 | 1.6594 | 0.0299 | 0.9960 | 0.7766 | 5.8394 |  | 1.217 | X |
| AP2B1_MOUSE | *Ap2b1* | AP-2 complex subunit beta | 2.3904 | 0.0227 | 0.9999 | 0.8375 |  |  | 2.433 |  |
| AP2M1_MOUSE | *Ap2m1* | AP-2 complex subunit mu | 2.5018 | 0.0466 | 0.9800 | 0.5819 | 5.1095 | X | 2.433 | X |
| ASTN1_MOUSE | *Astn1* | Astrotactin-1 | 1.6647 | 0.0160 | 0.9877 | 0.7745 | 0.2433 | X | 5.353 | X |
| AT2B4_MOUSE | *Atp2b4* | Plasma membrane calcium-transporting ATPase 4 | 1.7388 | 0.0114 | 0.9954 | 0.7361 | 3.1630 |  | 5.882 |  |
| ATAD3_MOUSE | *Atad3* | ATPase family AAA domain-containing protein 3 | 1.7738 | 0.0372 | 0.9826 | 0.4550 |  |  | 5.882 | X |
| ATPG_MOUSE | *Atp5f1c* | ATP synthase subunit gamma, mitochondrial | 3.3911 | 0.0316 | 1.0012 | 0.7784 |  |  | 5.882 | X |
| BAIP2_MOUSE | *Baiap2* | Brain-specific angiogenesis inhibitor 1-associated protein 2 | 2.5954 | 0.0460 | 0.9172 | 0.0518 | 0.9732 |  | 5.882 |  |
| CA2D1_MOUSE | *Cacna2d1* | Voltage-dependent calcium channel subunit alpha-2/delta-1 | 3.1679 | 0.0182 | 0.9515 | 0.2687 | 0.7299 |  | 11.76 | X |
| CAPS1_MOUSE | *Cadps* | Calcium-dependent secretion activator 1 | 2.4798 | 0.0483 | 0.9837 | 0.5833 | 0.4866 |  | 17.65 | X |
| CBPE_MOUSE | *Cpe* | Carboxypeptidase E | 3.1240 | 0.0467 | 1.0509 | 0.2354 | 0.0000 |  | 17.65 | X |
| CCG8_MOUSE | *Cacng8* | Voltage-dependent calcium channel gamma-8 subunit | 3.1719 | 0.0204 | 0.9912 | 0.6364 | 0.0000 | X | 22.14 | X |
| CD47_MOUSE | *Cd47* | Leukocyte surface antigen CD47 | 2.9829 | 0.0084 | 0.9688 | 0.3699 | 0.0000 | X | 29.41 |  |
| CISY_MOUSE | *Cs* | Citrate synthase, mitochondrial | 2.9715 | 0.0258 | 1.0134 | 0.8591 |  |  | 47.06 |  |
| CLH1_MOUSE | *Cltc* | Clathrin heavy chain 1 | 2.8044 | 0.0232 | 0.9978 | 0.8026 |  |  | 52.94 |  |
| CLUS_MOUSE | *Clu* | Clusterin | 7.8734 | 0.0278 | 1.0804 | 0.3769 | 1.7032 |  | 58.82 | X |
| COTL1_MOUSE | *Cotl1* | Coactosin-like protein | 2.2434 | 0.0267 | 1.0294 | 0.0758 | 0.2433 | X | 0 | X |
| COX2_MOUSE | *Mtco2* | Cytochrome c oxidase subunit 2 | 1.6676 | 0.0472 | 1.0305 | 0.5074 |  |  | 0 |  |
| COX41_MOUSE | *Cox4i1* | Cytochrome c oxidase subunit 4 isoform 1, mitochondrial | 8.0469 | 0.0285 | 0.9785 | 0.4664 | 4.3796 | X | 0 |  |
| CRIP2_MOUSE | *Crip2* | Cysteine-rich protein 2 | 6.8277 | 0.0496 | 0.9438 | 0.1959 | 0.4866 |  | 0 | X |
| CTNA2_MOUSE | *Ctnna2* | Catenin alpha-2 | 1.8749 | 0.0463 | 1.0109 | 0.8808 | 3.1630 |  | 0 | X |
| CYFP1_MOUSE | *Cyfp1* | Cytoplasmic FMR1-interacting protein 1 | 11.9841 | 0.0440 | 1.0567 | 0.3692 |  |  | 0 | X |
| DCE1_MOUSE | *Gad1* | Glutamate decarboxylase 1 | 2.8429 | 0.0430 | 1.0195 | 0.8945 | 0.0000 |  | 0 |  |
| DCE2_MOUSE | *Gad2* | Glutamate decarboxylase 2 | 1.877 | 0.050 | 1.052 | 0.246 | 0.000 |  | 0 | X |
| DCLK1_MOUSE | *Dclk1* | Serine/threonine-protein kinase DCLK1 | 2.6248 | 0.0212 | 1.0263 | 0.7258 | 0.4866 |  | 0 | X |
| DCTN1_MOUSE | *Dctn1* | Dynactin subunit 1 | 1.5601 | 0.0173 | 0.9741 | 0.3428 |  |  | 0 | X |
| DEST_MOUSE | *Dstn* | Destrin | 4.3473 | 0.0220 | 0.9684 | 0.4662 |  |  | 0 | X |
| DMXL2_MOUSE | *Dmxl2* | DmX-like protein 2 | 1.6352 | 0.0331 | 0.9837 | 0.5503 | 0.2433 |  | 0 | X |
| DNJC5_MOUSE | *Dnajc5* | DnaJ homolog subfamily C member 5 | 5.1200 | 0.0109 | 0.8396 | 0.0177 | 0.0000 | X | 0 | X |
| DPP6_MOUSE | *Dpp6* | Dipeptidyl aminopeptidase-like protein 6 | 1.5302 | 0.0185 | 0.9976 | 0.7721 | 0.2433 |  | 0 |  |
| DYN1_MOUSE | *Dnm1* | Dynamin-1 | 2.3159 | 0.0428 | 0.9753 | 0.4249 | 4.6229 |  | 0.243 | X |
| EFTU_MOUSE | *Tufm* | Elongation factor Tu, mitochondrial | 1.5273 | 0.0058 | 0.9989 | 0.7913 |  |  | 0.487 | X |
| EP15R_MOUSE | *Eps15l1* | Epidermal growth factor receptor substrate 15-like 1 | 1.8051 | 0.0421 | 0.9319 | 0.1354 | 6.5693 |  | 0.487 |  |
| ERP29_MOUSE | *Erp29* | Endoplasmic reticulum resident protein 29 | 17.3706 | 0.0265 | 1.0100 | 0.9527 | 6.5693 |  | 0.73 | X |
| F10A1_MOUSE | *St13* | Hsc70-interacting protein | 2.0988 | 0.0335 | 0.9517 | 0.2199 |  |  | 0.973 | X |
| FA49B_MOUSE | *Fam49b* | Protein FAM49B | 2.0721 | 0.0411 | 0.9483 | 0.2902 | 0.9732 |  | 0.973 |  |
| FAS_MOUSE | *Fasn* | Fatty acid synthase | 2.0774 | 0.0122 | 1.0023 | 0.9295 |  |  | 1.217 | X |
| FUMH_MOUSE | *Fh* | Fumarate hydratase, mitochondrial | 2.4212 | 0.0422 | 1.0295 | 0.6925 | 5.8394 |  | 2.433 |  |
| GNAI1_MOUSE | *Gnai1* | Guanine nucleotide-binding protein G(i) subunit alpha-1 | 3.3180 | 0.0287 | 0.9355 | 0.2010 |  | X | 3.406 | X |
| GNAI2_MOUSE | *Gnai2* | Guanine nucleotide-binding protein G(i) subunit alpha-2 | 4.6764 | 0.0374 | 0.9933 | 0.6470 |  | X | 3.406 | X |
| GNAO_MOUSE | *Gnao1* | Guanine nucleotide-binding protein G(o) subunit alpha | 4.8055 | 0.0254 | 1.0334 | 0.7333 |  | X | 3.893 | X |
| GNAQ_MOUSE | *Gnaq* | Guanine nucleotide-binding protein G(q) subunit alpha | 3.9366 | 0.0329 | 1.0152 | 0.9241 | 2.9197 | X | 4.136 |  |
| GNAZ_MOUSE | *Gnaz* | Guanine nucleotide-binding protein G(z) subunit alpha | 2.8957 | 0.0265 | 0.9815 | 0.5618 | 1.4599 | X | 4.38 | X |
| GPDA_MOUSE | *Gpd1* | Glycerol-3-phosphate dehydrogenase | 2.0714 | 0.0477 | 0.9542 | 0.2820 | 0.0000 |  | 5.109 | X |
| GRIA2_MOUSE | *Gria2* | Glutamate receptor 2 (GluR-2) | 2.0033 | 0.0397 | 0.9980 | 0.8023 | 0.0000 | X | 5.882 |  |
| HEXB_MOUSE | *Hexb* | Beta-hexosaminidase subunit beta | 2.2503 | 0.0402 | 1.2536 | 0.0001 | 1.7032 |  | 5.882 |  |
| HXK1_MOUSE | *Hk1* | Hexokinase-1 | 1.864 | 0.032 | 1.041 | 0.483 | 9.002 |  | 8.516 | X |
| HS12A_MOUSE | *Hspa12a* | Heat shock 70 kDa protein 12A | 2.0260 | 0.0458 | 1.0011 | 0.8019 | 0.2433 |  | 5.882 | X |
| HSP74_MOUSE | *Hspa4* | Heat shock 70 kDa protein 4 | 2.0908 | 0.0249 | 0.9931 | 0.7273 |  |  | 7.299 | X |
| IMPA1_MOUSE | *Impa1* | Inositol monophosphatase 1 | 3.8661 | 0.0392 | 0.9405 | 0.0634 | 2.1898 | X | 8.516 | X |
| IVD_MOUSE | *Ivd* | Isovaleryl-CoA dehydrogenase, mitochondrial | 1.6243 | 0.0289 | 1.0186 | 0.5851 | 2.4331 |  | 11.76 | X |
| KCC2D_MOUSE | *Camk2d* | Calcium/calmodulin-dependent protein kinase type II subunit delta | 1.5646 | 0.0170 | 0.9913 | 0.7451 | 3.6496 |  | 11.76 | X |
| KIF5C_MOUSE | *Kif5c* | Kinesin heavy chain isoform 5C | 1.6691 | 0.0351 | 0.9917 | 0.6342 |  |  | 11.76 | X |
| LETM1_MOUSE | *Letm1* | Mitochondrial proton/calcium exchanger protein | 1.8560 | 0.0428 | 0.9614 | 0.2821 | 4.3796 |  | 11.76 | X |
| LIPA3_MOUSE | *Ppfia3* | Liprin-alpha-3 | 1.6859 | 0.0239 | 0.9300 | 0.1882 | 0.2433 |  | 11.76 | X |
| LIS1_MOUSE | *Pafah1b1* | Platelet-activating factor acetylhydrolase IB subunit alpha | 2.0802 | 0.0263 | 0.9655 | 0.3596 | 4.3796 |  | 16.55 |  |
| M2OM_MOUSE | *Slc25a11* | Mitochondrial 2-oxoglutarate/malate carrier | 3.1066 | 0.0359 | 1.0222 | 0.8429 |  |  | 17.65 |  |
| MPP2_MOUSE | *Mpp2* | MAGUK p55 subfamily member 2 | 2.3788 | 0.0249 | 1.1196 | 0.0430 | 0.0000 | X | 17.65 |  |
| NAC1_MOUSE | *Slc8a1* | Sodium/calcium exchanger 1 | 1.6078 | 0.0206 | 1.0694 | 0.2621 | 0.2433 | X | 23.53 | X |
| NCALD_MOUSE | *Ncald* | Neurocalcin-delta | 45.0986 | 0.0333 | 0.8888 | 0.0594 | 0.0000 |  | 23.53 | X |
| NCKP1_MOUSE | *Nckap1* | Nck-associated protein 1 | 1.7610 | 0.0369 | 1.0207 | 0.6890 | 8.5158 |  | 25.06 |  |
| NRX3A_MOUSE | *Nrxn3* | Neurexin-3 | 2.8629 | 0.0254 | 1.0057 | 0.9531 | 0.0000 |  | 41.18 | X |
| NSF_MOUSE | *Nsf* | Vesicle-fusing ATPase | 1.7968 | 0.0111 | 0.9932 | 0.6500 | 7.5426 |  | 41.18 | X |
| OAT_MOUSE | *Oat* | Ornithine aminotransferase, mitochondrial | 2.3834 | 0.0408 | 1.1231 | 0.2428 |  |  | 47.06 |  |
| ODO1_MOUSE | *Ogdh* | 2-oxoglutarate dehydrogenase, mitochondrial | 2.7338 | 0.0474 | 1.0213 | 0.8685 | 7.7859 |  | 47.06 | X |
| ODPX_MOUSE | *Pdhx* | Pyruvate dehydrogenase protein X component, mitochondrial | 1.5552 | 0.0191 | 0.9543 | 0.0361 | 2.9197 |  | 52.94 |  |
| OPA1_MOUSE | *Opa1* | Dynamin-like 120 kDa protein, mitochondrial | 1.9052 | 0.0425 | 0.9759 | 0.3227 | 3.1630 | X | 64.71 | X |
| OXR1_MOUSE | *Oxr1* | Oxidation resistance protein 1 | 1.6551 | 0.0027 | 0.9682 | 0.3713 | 0.0000 |  | 0 |  |
| PCCA_MOUSE | *Pcca* | Propionyl-CoA carboxylase alpha chain, mitochondrial | 1.5242 | 0.0322 | 0.9542 | 0.2035 |  |  | 0 | X |
| PDE2A_MOUSE | *Pde2a* | cGMP-dependent 3',5'-cyclic phosphodiesterase | 1.7097 | 0.0100 | 1.0481 | 0.3392 | 0.7299 | X | 0.243 | X |
| PGM1_MOUSE | *Pgm1* | Phosphoglucomutase-1 | 1.9775 | 0.0013 | 1.0370 | 0.6444 | 3.8929 |  | 2.433 |  |
| PLCB1_MOUSE | *Plcb1* | 1-phosphatidylinositol 4,5-bisphosphate phosphodiesterase beta-1 | 1.9949 | 0.0198 | 1.0061 | 0.7702 | 0.0000 |  | 5.882 | X |
| PP2BA_MOUSE | *Ppp3ca* | Serine/threonine-protein phosphatase 2B catalytic subunit alpha isoform | 1.7693 | 0.0242 | 0.9565 | 0.1996 | 1.7032 | X | 11.76 |  |
| PRDX6_MOUSE | *Prdx6* | Peroxioredoxin-6 | 1.6130 | 0.0299 | 1.0689 | 0.0748 |  | X | 17.65 |  |
| PRRT3_MOUSE | *Prrt3* | Proline-rich transmembrane protein 3 | 2.2877 | 0.0290 | 0.9837 | 0.5961 | 0.2433 |  | 17.65 | X |
| PTPRD_MOUSE | *Ptprd* | Receptor-type tyrosine-protein phosphatase delta | 2.8642 | 0.0307 | 1.0097 | 0.9828 | 0.0000 | X | 17.65 | X |
| PTPRS_MOUSE | *Ptprs* | Receptor-type tyrosine-protein phosphatase S | 2.2591 | 0.0261 | 0.9756 | 0.4610 | 0.0000 |  | 29.41 |  |
| PYGB_MOUSE | *Pygb* | Glycogen phosphorylase, brain form | 3.9945 | 0.0321 | 1.0839 | 0.0150 | 4.6229 |  | 35.29 |  |
| RAB1A_MOUSE | *Rab1a* | Ras-related protein Rab-1A | 3.9014 | 0.0486 | 0.9073 | 0.0628 |  |  | 37.96 | X |
| RAB7A_MOUSE | *Rab7a* | Ras-related protein Rab-7a | 1.8488 | 0.0354 | 1.0068 | 0.8316 |  | X | 41.18 | X |
| RAP2B_MOUSE | *Rap2b* | Ras-related protein Rap-2b | 8.6218 | 0.0214 | 0.9733 | 0.4554 | 1.7032 | X | 0 | X |
| S12A2_MOUSE | *Slc12a2* | Solute carrier family 12 member 2 | 2.5952 | 0.0396 | 0.9641 | 0.3628 | 1.2165 |  | 0 | X |
| SAHH2_MOUSE | *Ahcyl1* | S-adenosylhomocysteine hydrolase-like protein 1 | 2.7124 | 0.0136 | 1.0495 | 0.2339 | 3.8929 |  | 0 | X |
| SATT_MOUSE | *Slc1a4* | Neutral amino acid transporter A | 8.4285 | 0.0440 | 1.2718 | 0.0002 | 0.0000 | X | 0.243 | X |
| SC6A1_MOUSE | *Slc6a1* | Sodium- and chloride-dependent GABA transporter 1 | 3.4145 | 0.0466 | 1.0470 | 0.4440 | 0.0000 | X | 0.973 | X |
| SCAM1_MOUSE | *Scamp1* | Secretory carrier-associated membrane protein 1 | 3.2759 | 0.0336 | 0.9115 | 0.0531 | 0.7299 | X | 1.46 | X |
| SDHB_MOUSE | *Sdhb* | Succinate dehydrogenase [ubiquinone] iron-sulfur subunit, mitochondrial | 2.8442 | 0.0205 | 1.0124 | 0.8729 | 4.3796 |  | 2.676 | X |
| SEPT8_MOUSE | *Septin8* | Septin-8 | 1.7848 | 0.0199 | 1.0574 | 0.3595 |  | X | 5.109 |  |
| SFXN1_MOUSE | *Sfxn1* | Sideroflexin-1 | 7.9333 | 0.0425 | 1.0657 | 0.3687 | 6.5693 | X | 5.882 | X |
| SFXN3_MOUSE | *Sfxn3* | Sideroflexin-3 | 2.3725 | 0.0162 | 0.9884 | 0.6007 | 0.9732 | X | 5.882 | X |
| SSDH_MOUSE | *Aldh5a1* | Succinate-semialdehyde dehydrogenase, mitochondrial | 1.6663 | 0.0050 | 1.0534 | 0.4846 | 4.3796 |  | 5.882 | X |
| SYIM_MOUSE | *lars2* | Isoleucine--tRNA ligase, mitochondrial | 1.8534 | 0.0348 | 1.0386 | 0.5893 | 4.3796 |  | 8.516 | X |
| SYNJ1_MOUSE | *Synj1* | Synaptojanin-1 | 2.1789 | 0.0289 | 1.0211 | 0.8514 | 2.9197 |  | 9.489 | X |
| SYNPR_MOUSE | *Synpr* | Synaptoporin | 6.445 | 0.018 | 0.963 | 0.365 | 0.000 |  | 11.76 | X |
| SYT2_MOUSE | *Syt2* | Synaptotagmin-2 | 2.4842 | 0.0494 | 0.9862 | 0.7697 | 0.0000 | X | 11.76 | X |
| SYT7_MOUSE | *Syt7* | Synaptotagmin-7 | 2.7289 | 0.0129 | 1.0495 | 0.6112 | 0.0000 | X | 11.76 | X |
| TERA_MOUSE | *Vcp* | Transitional endoplasmic reticulum ATPase) | 2.1826 | 0.0257 | 1.0105 | 0.9593 | 0.4063 |  | 14.36 |  |
| UBA1_MOUSE | *Uba1* | Ubiquitin-like modifier-activating enzyme 1 | 2.3035 | 0.0188 | 0.9517 | 0.2816 | 0.2603 |  | 23.53 | X |
| UCRI_MOUSE | *Uqcrrf1* | Cytochrome b-c1 complex subunit Rieske, mitochondrial | 4.8396 | 0.0492 | 1.1149 | 0.0848 | 2.4331 | X | 28.71 |  |
| VA0D1_MOUSE | *Atp6v0d1* | V-type proton ATPase subunit d | 4.6384 | 0.0334 | 1.0246 | 0.7738 | 1.2165 |  | 29.41 | X |
| VAMP1_MOUSE | *Vamp1* | Vesicle-associated membrane protein 1 | 6.0833 | 0.0172 | 0.8937 | 0.0789 | 0.0000 | X | 29.41 |  |
| VATE1_MOUSE | *Atp6v1e1* | V-type proton ATPase subunit E | 2.8166 | 0.0390 | 0.9533 | 0.1839 | 3.8929 |  | 29.41 |  |
| VPS35_MOUSE | *Vps35* | Vacuolar protein sorting-associated protein 35 | 2.2835 | 0.0102 | 0.9934 | 0.6715 | 8.5158 |  | 47.06 |  |
| VTA1_MOUSE | *Vta1* | Vacuolar protein sorting-associated protein VTA1 homolog | 39.3736 | 0.0420 | 0.9281 | 0.0333 | 2.1898 |  | 0 |  |

To determine whether our validated substrates are corroborated by prior computational or experimental evidence of palmitoylation, we cross-referenced validated hits with CSS Palm (Ren et al., 2008), Swiss Palm (Blanc et al., 2015), and Kang et al., 2008. Notably, these comparisons identified evidence of palmitoylation for 100% (26/26) of high confidence substrates (**Table 1**) and 94% (105/112) of medium-confidence substrates (**Table 2**). Validated proteins also include all known PPT1 substrates—CSPα, Goα, and F1 subunit O of mitochondrial ATP synthase—and exclude substrates of other depalmitoylating enzymes, such as PSD-95, an abundant synaptic protein that is exclusively depalmitoylated by ABHD17A, 17B, and 17C (Yokoi et al., 2016). Putative APT1 substrates such as RAB9A, RAB6A, N-RAS, R-RAS, GAP-43, and SNAP-23 are also excluded (Duncan and Gilman, 1998; Kong et al., 2013; Liu et al., 2019). We also screened our results against a PPT1 interactome dataset (Sapir et al., 2019) and found 4 high-confidence substrates (DYL2, LDHB, STXB1, THY1) and 7 medium-confidence substrates (DYN1, KIF5C, SSDH, SYT2, TERA, UBA1, VA0D1) to be present. Together, these data comprehensively substantiate that validated PPT1 substrates are palmitoylated proteins and that PPT1 has substrate specificity.

### Classification of PPT1 substrates highlights diverse synaptic functions

To obtain a better understanding of the functional role of PPT1-dependent depalmitoylation, the 138 high- and medium-confidence PPT1 substrates were analyzed using UniProt Gene Ontology, resulting in 9 distinct classes (**Figure 3C**): 1) Cytoskeletal proteins (n = 13) 2) Lysosomal proteins (n = 2); 3) Mitochondrial proteins (n = 23); 4) Synaptic cell adhesion molecules (n = 10); 5) Channels and transporters (n = 18); 6) Kinases and phosphatases (n = 6); 7) Exo- and endocytic proteins (n = 15); 8) Membrane proteins (n = 6); 9) G-protein-associated molecules (n = 11); and 10) Other proteins (n = 34). 75% of high- and medium-confidence validated proteins (104/138) fall nicely into defined classes with functions at the synapse. Our results agree with prior evidence that palmitoylation regulates ion channels and transporters, G-protein-associated molecules, and mitochondrial proteins (Goddard and Watts, 2012; Montersino and Thomas, 2015; Shipston, 2011). Our findings that synaptic adhesion molecules and endocytic proteins undergo PPT1-mediated depalmitoylation are novel.

To gain insight into the role of PPT1-mediated depalmitoylation at the synapse, we searched for the sub-synaptic localization of high-confidence substrates described in prior studies. Most proteins were demonstrated to be localized to the synaptic plasma membrane (15/26), with the remaining proteins found in the synaptic cytosol (6/26) or at organelles such as mitochondria and lysosomes (3/26; **Table S5**; UniProt). None of these substrates appear to be localized exclusively to the post-synaptic cytosol. The existence of both synaptic plasma membrane and cytosol substrates is intriguing and in line with the subcellular localization of PPT1 (**Figure S2B**). Western blot analysis of subcellular fractions from WT and PPT1 KO brains demonstrates that select validated PPT1 substrates (NRCAM, AT1B2, DNJC5, GRIA1; **Figure 3D** and synaptic markers (**Figure S2B**) are localized to the expected synaptic subcellular fraction.

We next investigated peptides with carbamidomethyl-modified cysteines (known palmitoylation sites) detected for high-confidence PPT1 substrates to identify common features. Interestingly, we find that the carbamidomethyl peptides for most membrane-associated substrates are extracellular and face the synaptic cleft (**Table S6**). Strikingly, we find that for all 12 peptides identified as extracellular and carbamidomethyl-modified, the modified cysteines participate in disulfide bonds in the mature protein (**Table S6**). This suggests that cysteines could be shared for palmitoylation and disulfide bond formation, and perhaps that intracellular palmitoylation precedes disulfide bond formation, as proposed previously (Bechtel et al., 2020; Itoh et al., 2018; Yu et al., 2017). Many of the proteins containing these extracellular peptides fall into the broad classes of cell adhesion molecules (NFASC, NRCAM, CADM2/SynCAM2, NTRI, THY1, BASI), and channels/transporters (GRIA1, AT1B1, AT1B2; **Figure 3C**).

To investigate the importance of cysteines that are shared for palmitoylation and disulfide bond formation, we studied the AMPA receptor (AMPAR) subunit GluA1 (GRIA1). GluA1 was identified as a high-confidence PPT1 substrate with a palmitoylation site on C323. Based on a recent crystal structure, C323, which typically forms a disulfide bond to C75 (**Table S6**), is found in an extracellular loop (Herguedas et al., 2019). This site has not been previously identified as palmitoylated, though GluA1 has known intracellular and transmembrane palmitoylation sites (C811 and C585, respectively; Herguedas et al., 2019). Since aberrant palmitoylation of these sites has functional consequences (Itoh et al., 2018; Spinelli et al., 2009), we examined whether a GluA1 Cysteine-Alanine (C323A) point mutation alters AMPAR function. When expressed in *Xenopus laevis* ooctyes, both with auxiliary subunit TARPγ8 and independently at high concentration, GluA1 C323A exhibits abolished glutamate-evoked AMPAR currents (**Figure 3E**). Intriguingly, GluA2 and AMPAR accessory factors (AP2A1, AP2B1, AP2M1, NSF, KCC2D) were also identified as PPT1 substrates (**Figure 3C**).

We used Ingenuity Pathway Analysis (IPA) to determine functional pathways in which high- and medium-confidence PPT1 substrates participate in an unbiased manner. The top pathways identified were (1) GABA receptor signaling, (2) CREB signaling in neurons, (3) Synaptic long-term depression, (4) Endocannabinoid neuronal synapse pathway, and (5) Synaptogenesis signaling pathway (**Figure 3F**). These functional pathways are consistent with *CLN1* phenotypes and the 9 ontological classes we defined (**Figure 3C**).

### Structural motifs contribute to PPT1 substrate specificity

Since we find that PPT1 is moderately specific and confirmed that it can depalmitoylate 138 proteins *in vivo* (∼10% of the synaptic palmitome), we carried out two independent analyses to identify determinants of its substrate specificity. First, we sought to determine whether classes of PPT1 substrates with several high confidence hits have distinct recognition sites. We discovered that the synaptic adhesion PPT1 substrates are conserved and belong to the Immunoglobulin (Ig) domain-containing class of homotypic adhesion molecules (**Figure 4A**). The best characterized of these are CADM2 (SynCAM2) and NRCAM. Based on their structures, we identified that the carbamidomethyl peptides in most of these proteins map to one of two crucial cysteines in extracellular IgG domains (**Figure 4B-C**; Fogel et al., 2010). The palmitoylated cysteines are on exposed and accessible beta-strands and normally form intra-domain disulfide bonds to maintain the typical IgG domain fold (**Figure 4C**). The palmitoylated cysteines identified in the other synaptic adhesion molecules are known to mediate intra- or intermolecular interactions via disulfide bonds (Fogel et al., 2011). Decreased colocalization of CADM2 and SYPH in PPT1 KO neurons (**Figure 4D-E**) suggests that altered depalmitoylation of synaptic adhesion molecules impacts subsynaptic localization. Overall, our data suggest that PPT1 can recognize the IgG sequence or fold.

**Figure 4.**
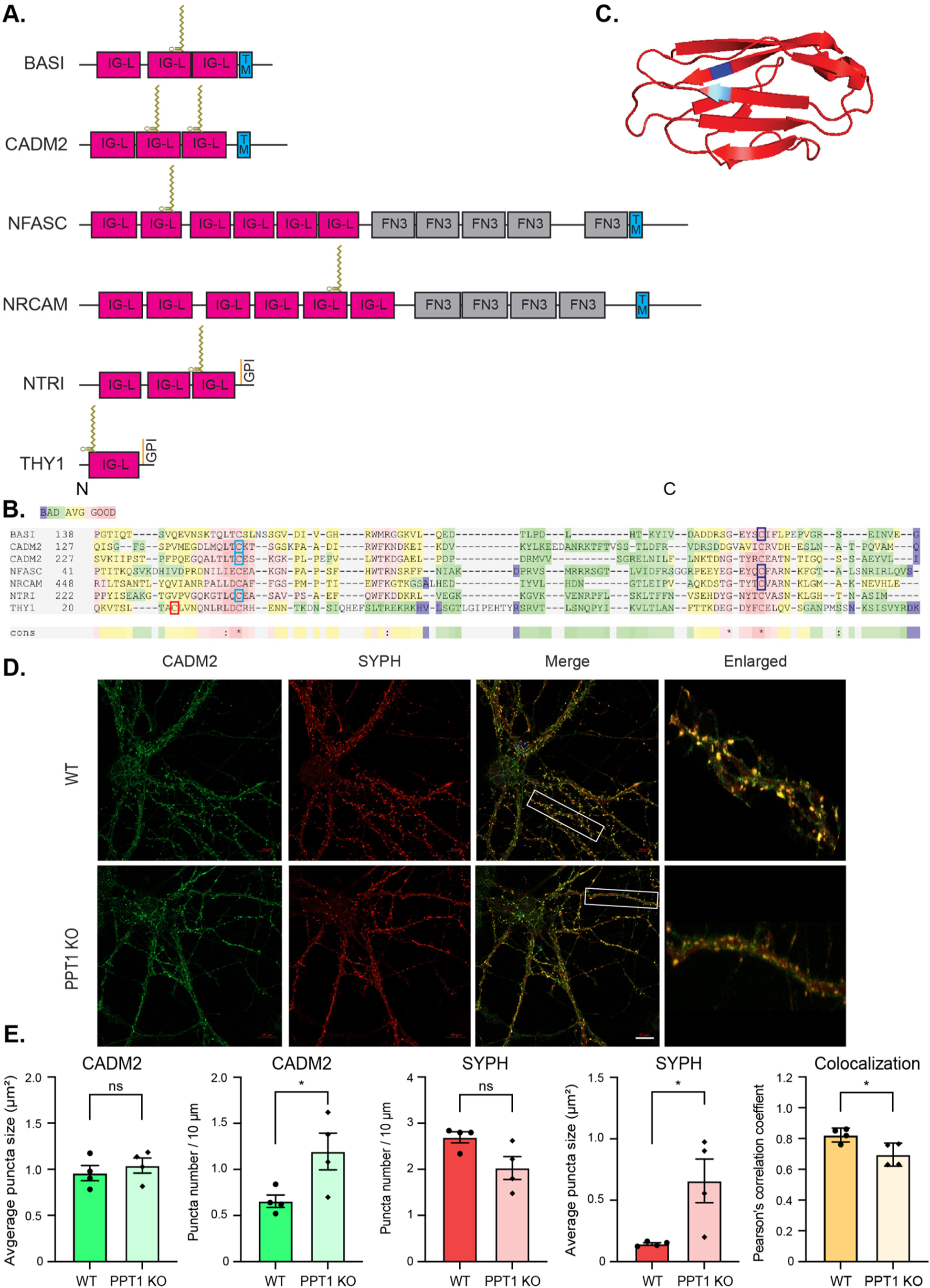
PPT1 depalmitoylates the IgG domain of synaptic adhesion molecules. A) Domain organization of IgG domain-containing synaptic adhesion molecules identified as high-confidence PPT1 substrates with palmitoylation sites represented as lipid chains. IG-L – Ig-like; FN3 – fibronectin 3; TM – transmembrane. B) IgG domain-containing synaptic adhesion molecules show high homology surrounding the palmitoylated cysteines. Red indicates high homology; green indicates low homology. The two conserved cysteines at the N- (light blue) and C- termini (dark blue) of this domain are both identified as carbamidomethylated in different experiments. C) Representative IgG domain structure (CADM2; PDB, 3M45) with position of palmitoylated cysteines highlighted in blue. D) Endogenous syncam2 (CADM2) and synaptophysin 1 (SYPH) colocalize on MAP2+ neurites in WT and PPT1 KO mouse primary neuronal cultures (scale bar 10 µm; 50 µm enlarged ROI). E) Quantifications of syncam2 and synaptophysin 1 puncta number (per 10 µm neurite segment), average puncta size (µm^2^), and colocalization (Pearson’s correlation coefficient). Data presented as mean of 5 ROIs per culture ± SD (n = 4 cultures; * p > 0.5).

Next, we tested whether motifs recognized by partner palmitoylating enzymes are also the primary determinant for PPT1 substrate recognition. Two DHHC/PATs, DHHC5 and DHHC17, localize to synapses (Li et al., 2010; Pandya et al., 2017). DHHC17 and its *Drosophila* homolog, HIP-14, are known to traffic many synaptic proteins in a palmitoylation-dependent manner, including the PPT1 substrate CSPα (Greaves et al., 2009). DHHC17 is the only PAT with a known substrate-recognition motif (Lemonidis et al., 2015). Its recognition sequence ([VIAP][VIT]xxQP) is usually adjacent to the palmitoylated cysteine. We screened the 138 validated PPT1 substrates for those containing the DHHC17 recognition motif and identified 9 cytosolic proteins, including CLH1 (Clathrin heavy chain), DYN1 (Dynamin 1), SYNJ1 (Synaptojanin 1), and DNJC5 (CSPα) (**Tables 1-2; Figure S3A**). IPA identified “Clathrin-mediated endocytosis signaling” as the top pathway involving this subset of PPT1 substrates (**Figure S3B**), consistent with the known function these proteins play in synaptic vesicle endocytosis (Saheki and De Camilli, 2012). Together, these data suggest that the DHHC17 recognition site is a determinant of substrate specificity for the endocytic class of PPT1 substrates.

### Tertiary post-translational modification screen detects coincidence of palmitoylation and disulfide bonds

To more directly query the relationship between disulfide bond formation and palmitoylation amongst validated PPT1 substrates, we performed a modified Acyl RAC screen of synaptosomes prepared from 3 WT and 3 PPT1 KO mouse brains (each in technical triplicate; age = 2 months; **Figure 5A**). First, free thiols were labelled with NEM (**Figure 5Ai**), then TCEP was used to reduce disulfide bonds (**Figure 5Aii**). The newly generated thiol groups were then labelled with d_5_-*N*-ethylmaleimide (dNEM), which has +5 molecular weight shift due to deuterium (McDonagh et al., 2015), allowing for the differentiation of cysteines that exist as free thiols versus those in disulfide bonds (**Figure 5Aii**). The remainder of the Acyl RAC workflow (**Figure 5Aiii–ix**) was then conducted as previously described (**Figure 1B**) to isolate the synaptic palmitome for LFQ MS/MS (n = 3551 proteins). Carbamidomethyl peptides were detected for 100% (26/26) of high-confidence substrates (**Table S5**), 59% (66/112) of medium-confidence substrates, and 0% of residual substrates in this tertiary screen, indicating excellent reproducibility and highlighting the efficacy of our stringent confidence thresholds.

**Figure 5.**
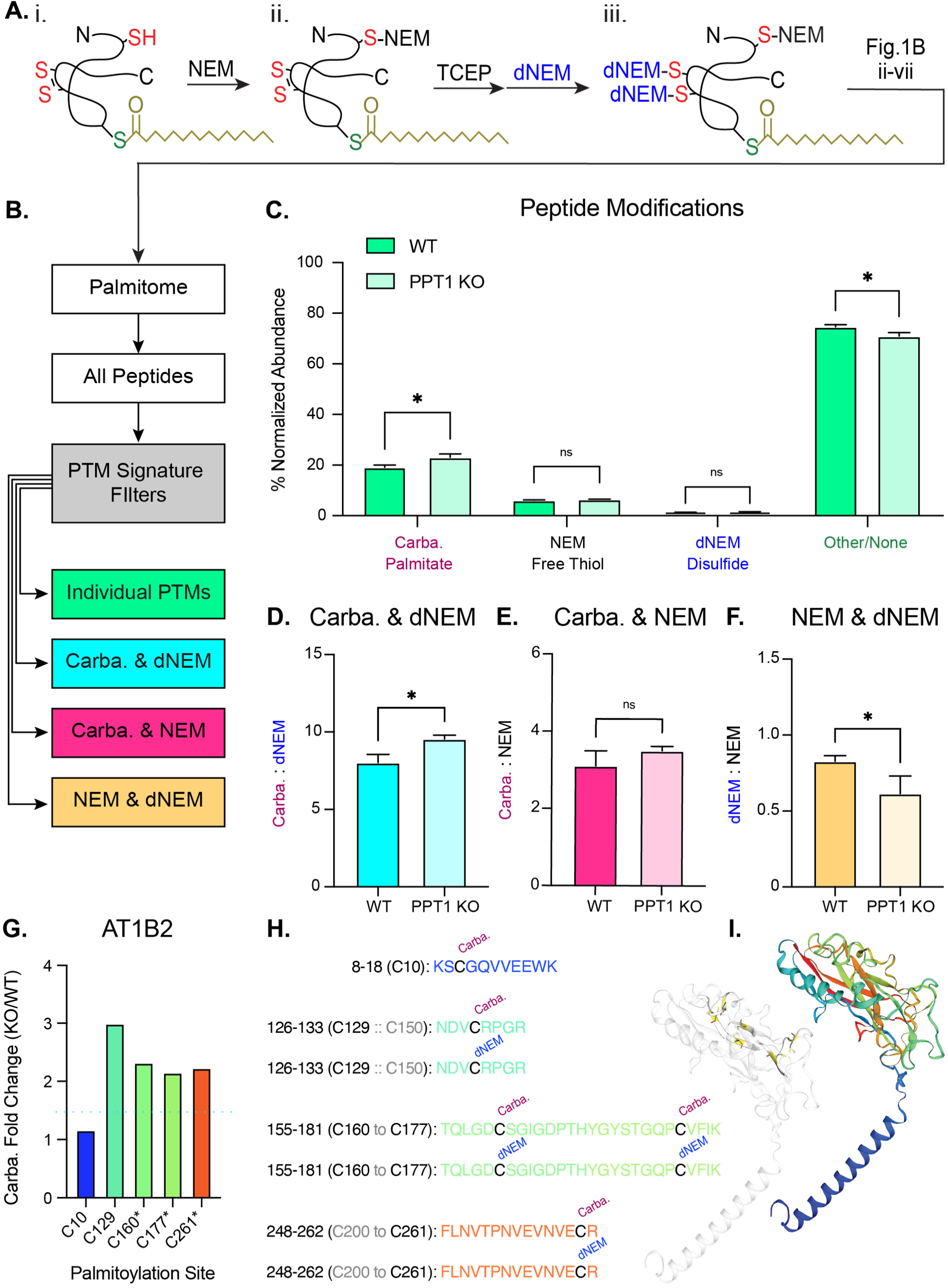
PPT1-mediated depalmitoylation facilitates disulfide bond formation. A) Schematic of modified Acyl RAC. i. Free thiols were blocked with NEM prior to reduction of disulfide bonds with TCEP. ii. Free thiols generated by TCEP treatment were modified with heavy-labeled NEM (dNEM). iii. Acyl RAC was then conducted as previously described (**Figure 1Bii-iv**). B) Pipeline for post-translational modification (PTM) analysis of Acyl RAC peptide data. Peptides were aggregated from three independent proteomic screens with 3 biological replicates. All peptides were filtered by individual PTMs or by coincidence signatures where two distinct PTMs were found to modify the same cysteine residue. C) Global distribution of modified peptides. Bars represent percent (%) average normalized abundance of the different peptide classes. The value of each replicate is calculated as the summed abundance of peptides with the given PTM normalized to the total abundance for all detected peptides. Cysteine residues with PTMs are interpreted as follows: carbamidomethyl (carba.) = palmitate, N-ethylmaleimide (NEM) = free thiol, d_5_-*N*-ethylmaleimide (dNEM) = disulfide bond, other or no moieties (Other/None). D-F) Ratios of PTM normalized abundance for peptides where two PTMs are found to modify the same cysteine residue. The value of each replicate is calculated as the summed abundance of peptides with the first PTM divided by the summed normalized abundance of peptides with the second PTM (mean ± SD; * p < 0.05, unpaired, two-tailed t-test). G) Peptide analysis of a high-confidence PPT1 substrate, Sodium/Potassium transporting ATPase subunit β1 (AT1B2). Normalized abundance fold change (PPT1 KO/WT) of carbamidomethyl AT1B2 peptides (1.5-fold change indicated by blue dashed line). Palmitoylated cysteine sites marked with an asterisk (*) were confirmed to be PPT1 substrates in the validation screen. There is a persistence of palmitates on cysteines that normally participate in disulfide bonds in PPT1 KO (C129, C160, C177, C261; **Table S6**), while C10, which is not predicted to form a disulfide linkage, appears unchanged. H) Detected AT1B2 peptides are indicated. For those sequences with cysteines predicted to form disulfide bonds, carba. and dNEM moieties were found to modify that cysteine residue. I) Ribbon structure of AT1B2 (SWISS-MODEL; P14231) with disulfide bonds highlighted. Bar graph and peptide sequences are color-coded according to location in the ribbon structure.

To examine PTMs, we aggregated peptide data from the three biological replicates to capture as many modified peptides per protein as possible (n = 16,381 total peptides). We then filtered peptides by individual PTMs of interest (**Figure 5B**): carbamidomethyl (carba.) indicates the presence of a palmitate group, NEM indicates a free thiol, dNEM indicates a disulfide bond, and Other/None describes unmodified peptides or those with a PTM not pertinent to this study (**Figure 5C**). Carbamidomethyl-modifications were the most abundant PTM of interest (WT = 19.2 ± 0.9%; PPT1 KO = 23.0 ± 1.3%). As expected, carbamidomethyl peptides were significantly more abundant in the PPT1 KO (p = 0.0147), reflecting the increase in palmitoylated substrates due to PPT1 deficiency. Other/None peptides exhibited a concomitant significant decrease in the PPT1 KO (WT = 74.6 ± 0.83%; PPT1 KO = 70.60 ± 1.44%; p = 0.0142). NEM peptides were also abundant but did not display significant changes between genotypes (WT = 6.2 ± 0.2%; PPT1 KO = 6.3 ± 0.1%; p = 0.4688). dNEM-modified peptides were rarest, constituting less than 1% of the total normalized peptide abundance (WT = 0.83 ± 0.03%; PPT1 KO = 0.97 ± 0.1%; p = 0.1302; **Figure 5C**).

We next examined instances where multiple PTMs are found to modify the same cysteine residue (**Figure 5D-F**) by comparing the lists of peptides curated by modification. We identified sequences present in multiple lists (**Figure 5B**), then examined the ratio of the coincident PTMs (**Figure D-F**). A significant increase in carba.:dNEM was observed in PPT1 KO (WT = 8.02 ± 0.53; PPT1 KO = 9.55 ± 0.25; p = 0.0108; **Figure 5D**), indicating the persistence of palmitoyl groups on cysteines that form disulfide bonds. We did not observe significant changes to carba.:NEM in PPT1 KO (WT = 3.10 ± 0.39; PPT1 KO = 3.50 ± 0.11; p = 0.1669; **Figure 5E**), suggesting that excess palmitoylation alone does not significantly affect the proportion of free thiols. dNEM:NEM decreased significantly in PPT1 KO (WT = 0.83 ± 0.04; PPT1 KO = 0.61 ± 0.12%, p = 0.0385; **Figure 5F**). Together, these data suggest that there is a persistence of palmitates in PPT1 KO on disulfide bonding cysteines. We next examined our curated peptide modification data with a focus on high confidence PPT1 substrates. Carbamidomethyl moieties were detected on 94% (17/18) of validated PPT1 depalmitoylation sites predicted to form a disulfide bond (sole exception is NRCAM C519; **Table S5**). This result again highlights the reproducibility of our method and corroborates our previous observation that palmitoylation and disulfide bond formation may have a functional relationship. dNEM moieties, validating the presence of a disulfide bond, were found on 33% (6/18) of these sites (AT1B1 C126, AT1B2 C160 & C177, AT1B2 C261, GPM6A C192, THY1 C28; **Table S5**). Since three of these sites were from a single protein, the Na^+^/K^+^ transporting ATPase subunit β2 (AT1B2), we chose to investigate the peptide data for this protein in greater detail (**Figure 5G-I**). Four unique AT1B2 peptides with five carbamidomethylated cysteines (C10, C129, C160 & C177, C261) were identified in the tertiary screen (**Figure 5G-H**). C10, which is not predicted to form a disulfide bond, was previously identified as a palmitoylation site (Collins et al., 2017). The other palmitoylation sites are novel to the best of our knowledge and are predicted to form disulfide bonds (**Table S5**). Our data bear this out, as each of these sites was found to be modified by dNEM. C160, C177 and C261 are high-confidence PPT1 substrate sites (indicated by an asterisk in **Figure 5G**), while C10 and C129 did not meet this criterion in the initial screens. We examined normalized abundance fold change (PPT1 KO/WT) of AT1B2 carbamidomethyl peptides and observed a persistence of palmitate groups on cysteines that normally participate in disulfide bonds in PPT1 KO (**Figure 5G**). This result corroborates our data that C160, C177, and C261 are PPT1 depalmitoylation sites, and suggests that C129 may be as well, while C10 is likely not. Examination of the crystal structure of AT1B2 reveals that C129, C160, C177, and C261 account for all disulfide linkages in this molecule which serve to stabilize β-sheets that form the channel pore (**Figure 5I;** SWISS-MODEL; Bienert et al., 2017).

## DISCUSSION

We systematically and quantitatively characterized palmitoylation in WT and PPT1 KO mice and identified and validated PPT1 substrates. The broad and quantitative nature of our approach also enabled us to illuminate novel functions of depalmitoylation at the synapse. Here, we highlight the most salient insights revealed by our analyses to encourage others to use this resource in their own investigations.

### Generation of brain and synaptic palmitomes

In this study, we present a comprehensive dataset of brain (n = 1795) and synaptic palmitomes (n = 1378). This is the largest mouse brain palmitome to date and is a valuable resource to the scientific community (PRIDE accession PXD017270).

### Stringent validation of PPT1 substrates

As a result of our two-step screen, we identified 138 high- to medium-confidence PPT1 substrates. We are convinced that these proteins are *bona fide* substrates due to the stringent nature of our selection process: 1) Only proteins that were independently identified in 3 biological samples with robust identification measures were included (>2 unique peptides); 2) *In vivo* differences (KO/WT >1.5; p <0.05) were utilized as initial criteria for selection (**Figures 2, S1**); 3) Putative substrates were independently validated by direct PPT1 enzymatic depalmitoylation (**Figure 3**); 4) The tertiary screen confirmed the majority of the high- to medium confidence hits and, importantly, none of the non-validated or residual hits (**Figure 5**). 5) Previously published PPT1 substrates were validated (**Tables 1, 2**); 6) Palmitoylated proteins known to be exclusively depalmitoylated by AHBD-17 or APT1 were not identified, indicating specificity; 7) The PPT1 substrates cluster into known functional groups (**Figure 3**); 8) Synaptic adhesion molecules and endocytic PPT1 substrates have common structural motifs that explain substrate specificity (**Figures 4**, **S3**).

Collectively, our data indicate that PPT1 is a moderately selective enzyme that depalmitoylates ∼10% of the synaptic palmitome, while other depalmitoylating enzymes (APTs, ABHDs) are likely to depalmitoylate the remaining palmitoylated proteins. As with any proteome-wide screen, a fraction of PPT1 substrates was likely not identified, possibly amongst residual hits (**Table S4**) and should be reexamined in a case-by-case manner. Furthermore, PPT1 and other depalmitoylating enzymes may have overlapping substrates, although our results suggest this is uncommon, as the protein expression of APTs and ABHDs does not compensate for loss of PPT1 (**Figures 2D, S1D**).

### Elucidation of the relationship between PPT1-mediated depalmitoylation and protein expression

PPT1 depalmitoylation activity has long been considered necessary for protein degradation (Koster and Yoshii, 2019; Lu et al., 1996), as CLN1 patient-derived cells exhibit accumulation of lipid modified proteins (Hofmann et al., 1997; Lu et al., 1996). Having successfully identified substrates of PPT1, we find that in PPT1 KO, most substrates show significantly increased palmitoylation independently from increased protein expression, and exhibit either unchanged or decreased protein levels at the synapse (**Figure 2H**). This analysis suggests that for most proteins, removal of the palmitate group by PPT1 is either not required for proteasomal degradation, or another depalmitoylating enzyme facilitates this process. The noteworthy exceptions are the group of proteins with both increased expression and palmitoylation (n = 4; ASAH1, CATD, SCRB2, TPP1; **Figure 2H**), which precluded us from identifying them as PPT1 substrates in this study. TPP1 and CATD accumulation were previously observed in a mouse brain lysosomal proteome of late-stage *CLN1*, while SCRB2 and TPP1 accumulate in a comparable model of *CLN3* (Sleat et al., 2019), suggesting that these changes are relevant to NCL pathogenesis. It is possible that these proteins exhibit depalmitoylation-dependent degradation, as previously suggested to explain lipidated protein accumulation in *CLN1* disease (Hofmann et al., 1997). The contributions of ASAH1, CATD, SCRB2, and TPP1 to *CLN1*, and the etiological links between NCLs drawn by these proteins, are intriguing areas of investigation that fall beyond the scope of this study.

### Palmitoylation-depalmitoylation cycles at the synapse

The PATs DHHC5 and DHHC17 have been described to function locally and continuously to palmitoylate synaptic substrates (Fukata et al., 2004; Sialana et al., 2016). We documented a likely DHHC17-PPT1 partnership for palmitoylation-depalmitoylation cycles at the synapse (**Figure S3**) by identifying 9 synaptic PPT1 substrates that contain the DHHC17 recognition motif ([VIAP][VIT]xxQP; Lemonidis et al., 2015). These include CLH1, DYN1, SYNJ1, and DNJC5 (**Table S2**), which have known functions in synaptic vesicle recycling (Ferguson et al., 2007; Hinshaw, 2000; Newton et al., 2006; Royle and Lagnado, 2010; Umbach et al., 1994; Verstreken et al., 2003; Zhang et al., 2012; Zinsmaier et al., 1994). Strikingly, previous studies have identified synaptic vesicle endocytosis deficits in PPT1 KO neurons (Virmani et al., 2005), including reduced pre-synaptic vesicle pool size, deficits in evoked pre-synaptic vesicle release (Kim et al., 2008; Virmani et al., 2005), and the persistence of VAMP2 and SNAP25 on the presynaptic membrane (Kim et al., 2008). We hypothesize that palmitoylation-depalmitoylation dynamics, regulated by DHHC17 and PPT1, respectively, allow for efficient membrane association and dissociation of endocytic proteins during synaptic vesicle endocytosis. However, the exact timing and localization of depalmitoylation and the triggers for this cycle remain to be clarified. Identification of motifs recognized by DHHC5 may provide additional insights into local regulation of other PPT1 substrates.

### Depalmitoylation facilitates disulfide bond formation and critical synaptic functions

Previous literature has established that palmitoylation of synaptic proteins is essential for their trafficking from the soma to the nerve terminal (Fukata and Fukata, 2010; Kanaani et al., 2004). The palmitoylation of synaptic proteins destined for the secretory pathway is accomplished by resident PATs in the lumen of the ER or Golgi (Bechtel et al., 2020), while for other proteins, palmitoylation occurs on the cytosolic surface of these organelles. Though the precise PATs serving these distinct roles are not known, palmitoylation facilitates sorting (secretory) and hitching (cytosolic) of synaptic proteins to the synaptic vesicle precursor membrane (Greaves and Chamberlain, 2007; Hayashi et al., 2005). Beyond keeping synaptically-targeted proteins tethered to vesicles, we hypothesize that palmitoylation keeps synaptic proteins inert during axonal trafficking by preventing the formation of disulfide bonds. Hence, depalmitoylation by PPT1 may relieve brakes on ectopic adhesion interactions or enzymatic activity enforced by palmitate-mediated blockage of disulfide bonding. Once at the synapse, cytosolic proteins (such as G-proteins, endocytic proteins, kinases, and phosphatases) are released from the synaptic vesicle precursor, while membrane proteins (synaptic adhesion molecules, channels, transporters) are inserted into the synaptic membrane and interact in both *cis* and *trans*. In the case of membrane proteins with extracellular palmitoylation sites, it is likely that depalmitoylated cysteines spontaneously form disulfide bonds in the oxidative extracellular environment, leading to functional maturation (Bechtel et al., 2020; Cijsouw et al., 2018). Our results suggest that local synaptic depalmitoylation by PPT1 modulates these functions, though they do not exclude PPT1-mediated depalmitoylation of substrates en route to the synapse. In PPT1 KO brains, we predict that palmitoylated synaptic proteins traffic correctly (as PATs are normal, and we observe few changes in the synaptic proteome; **Figure 2D**) but may function poorly at the synapse due to absent or compromised depalmitoylation (**Figure 2B**). In line with this hypothesis, we confirmed the synaptic localization of several validated PPT1 substrates in PPT1 KO by subcellular fractionation (**Figure 3D, Figure S2B**).

Our finding that PPT1 depalmitoylates IgG domains of cell adhesion molecules suggests an important role for PPT1 in regulating synaptogenesis (**Figure 4A-C**). This finding is congruent with the impact of depalmitoylation on “synaptogenesis signaling” in PPT1 KO, identified by IPA (**Figure 3F**), as well as with previous findings on the importance of NRCAM palmitoylation for neuronal morphogenesis (Ponimaskin et al., 2008). A *Drosophila* orthologue of NRCAM was also identified in a genetic modifier screen of fly PPT1 KO-induced degeneration (Buff et al., 2007). Indeed, the aberrant palmitoylation of IgG-class adhesion molecule substrates may explain post-synaptic deficits previously described in *CLN1* models, including immature dendritic spine morphology of cultured PPT1-null neurons (Koster and Yoshii, 2019; Sapir et al., 2019). Similarly, our finding that PPT1 depalmitoylates GluA1 and several AMPAR accessory factors is congruent with “synaptic long-term depression” identified by IPA (**Figure 3F**), long-term potentiation (LTP) impairments reported in PPT1-deficient mice (Sapir et al., 2019), and accumulating evidence that palmitoylation plays roles in modulating structural synaptic plasticity (Ji and Skup, 2021). Since hyperpalmitoylation of known GluA1 palmitoylation sites can alter seizure susceptibility (Itoh et al., 2017), our discovery of a novel palmitoylation site that is critical for AMPAR function (C323, **Figure 3E**) opens avenues for investigation of the molecular basis of epilepsy in *CLN1*.

We identified AT1B2, the β2 subunit of the Na^+^/K^+^ transporting ATPase heterodimer, as a high-confidence PPT1 substrate. AT1B2 contains three pairs of disulfide-bonded cysteines (validated by dNEM moieties) that exhibit aberrant palmitoylation in the PPT1 KO. These data strongly suggest that PPT1-mediated depalmitoylation facilitates the stabilization of β-sheet structures in this molecule through the regulation of disulfide bonds (**Figure 5G-I**). Other Na^+^/K^+^ ATPase isoforms, AT1A1, AT1A2, AT1A3, and AT1B1, were also identified as high-confidence PPT1 substrates, accounting for all Na^+^/K^+^ ATPase subunits known to be expressed in neurons (**Figure 3C**; Senner et al., 2003). Similar to AT1B2, the β1 subunit (AT1B1) displays persistent palmitoylation in the PPT1 KO on a disulfide-bonded cysteine (C126; **Table S5**). α subunits are not predicted to form disulfide bonds, but also exhibit excessive palmitoylation, as is expected of PPT1 substrates (**Table S5**). AT1B2 and AT1A3 are known form a complex that is essential for anchoring the protein retinoschisin (RS1) to plasma membranes. In the retina, this cell-adhesion interaction maintains retinal architecture and the photoreceptor-bipolar cell synaptic structure (Molday et al., 2007; Friedrich et al., 2010; Plössl et al., 2019). Mutations in RS1 lead to juvenile retinoschisis, a form of early-onset macular degeneration (Molday et al., 2007; Friedrich et al., 2010; Plössl et al., 2019). Since juvenile retinal degeneration is also a feature of NCLs, these two diseases can be conflated and misdiagnosed in the clinic (Wright et al., 2020). This is the first piece of molecular evidence, to our knowledge, that hints at an etiological overlap. Furthermore, basigin (BASI), which is essential for normal retinal development (Hori et al., 2000), was also found to be a PPT1 substrate and colocalizes with AT1B2 in the retina (Lobato-Álvarez et al., 2016). AT1B2 has also been shown to regulate neuron-astrocyte adhesion (Senner et al., 2003), although its role as a cell adhesion molecule in the brain is not fully characterized. Na^+^/K^+^ ATPase subunits are classified as Channels/Transporters (**Figure 3C**) but can also be considered cell adhesion molecules; high-confidence PPT1 substrates predominantly fall in this category, highlighting the importance of depalmitoylation in synaptic adhesion interactions.

## CONCLUSIONS

Identification of synaptic PPT1 substrates provides a rational basis for investigating fundamental mechanisms by which depalmitoylation regulates synaptic functions. As aberrant palmitoylation has been described in Huntington’s disease, Alzheimer’s Disease, and amyotrophic lateral sclerosis (ALS), these data may have implications for multiple neurodegenerative disorders. This resource also establishes a detailed landscape of perturbations of depalmitoylation in NCL, opening avenues for molecular dissection of disease mechanisms.

## Acknowledgments

This work was supported by NIH (R01 NS064963, R01 NS110354, R01 NS083846, R21 NS094971), and DOD (W81XWH-17-1-0564). The proteomic experiments were supported by the Yale/NIDA Neuroproteomic Center (NIH DA018343). We would like to thank Weiwei Wang and Jean Kanyo for mass spectrometry sample preparation and data collection, and John E. Lee, Angus Nairn, and Arthur Horwich for reading and editing the manuscript. The Q-Exactive Plus mass spectrometer was funded in part by NIH SIG from the Office of The Director, NIH (S10OD018034).

## STAR METHODS

### LEAD CONTACT AND MATERIALS AVAILABILITY

Sreeganga Chandra is the lead contact for this paper. Mass spectrometry data are available through the ProteomeXchange Consortium in the PRIDE partner repository with accession PXD017270.

### EXPERIMENTAL MODEL AND SUBJECT DETAILS

PPT1 KO (B6;129-Ppt1tm1Hof/J), and WT C57BL6/J mice were obtained from The Jackson Labs. Animal care and housing complied with the Guide for the Care and Use of Laboratory Animals (National Academies Press, 2011) and were provided by the Yale Animal Resource Center (YARC). Animals were maintained in a 12-hour light/dark cycle with *ad libitum* access to food and water. Mice were 2 months of age and of both sexes. Sex differences were not measured because sexes were equally mixed in synaptosome experiments. Female *Xenopus laevis* frogs were obtained from Nasco for use in oocyte preparation. All experimental protocols involving animals were approved by the Institutional Animal Care & Use Committee (IACUC) at Yale University.

HEK 293T cells were obtained from American Type Culture Collection (ATCC).

### METHOD DETAILS

#### Preparation of synaptosomes

Forebrains were freshly harvested from 2 mice (age = 2 months), suspended in ice cold Buffer A (320 mM sucrose, 10 mM HEPES, pH 7.4 with protease inhibitors), then homogenized in 12 up-down passes at 900 rpm in a glass Teflon homogenizer. Homogenate (total) was centrifuged at 800 g for 10 minutes, 4°C. The supernatant (S1) was then centrifuged at 9000g for 15 minutes, 4°C. The resulting pellet was resuspended in 3 mL ice cold Buffer A and centrifuged at 9000g for 15 minutes, 4°C to obtain the synaptosomal supernatant (S2) and washed synaptosomes (P2’) for Acyl RAC and PPT1 validation.

#### Synaptic fractionation

Following synaptosome preparation from WT and PPT1 KO mouse whole brains (n=8 per genotype; age = 2 months), synaptic fractions were prepared as described by Huttner et al., 1983. Briefly, synaptosomes were lysed by hypoosmotic shock and the lysis supernatant (LS1) and synaptosomal membrane pellet (LP1) were collected. LP1 was resuspended and LS1 was centrifuged at 100,000g, 4°C for 1 hour to obtain synaptic vesicles (LP2) and synaptic cytosol (LS2). LP1 was then subjected to sucrose density gradient centrifugation at 48,000 g, 4°C for 2.5 hours to obtain the myelin fraction (MF), synaptic plasma membrane (SPM), and mitochondrial fraction (Mito.).

#### Acyl Resin-Assisted Capture

Acyl RAC was performed essentially as described in Henderson et al., 2016. Briefly, homogenized whole brains or prepared synaptosomes were blocked in 10 mM *N*-ethylmaleimide (NEM), then subjected to chloroform-methanol precipitation (Chloroform:MeOH:water :: 1:4:3 volumes) to remove free NEM. Samples were resuspended in 2% SDS, 50 mM Tris, 5 mM EDTA, pH 7.0 with protease inhibitors and 10 mM tris(2-carboxyethyl)phosphine (TCEP) and resolublized for 20-40 minutes at 37°C. Samples were then incubated at 4°C overnight with 10 mM NEM. Two additional chloroform-methanol precipitations were performed. Protein was resuspended in 2% SDS, 50 mM Tris, 5 mM EDTA, pH 7.4 and incubated at 37°C until complete dissolution. Samples were diluted 1:10 with 150 mM NaCl, 50 mM Tris, 5 mM EDTA, pH 7.4 with 0.2% Triton X-100, 1 mM PMSF, and protease inhibitor cocktail following dissolution. Another chloroform methanol precipitation was performed, and the protein precipitate was dissolved in 4% SDS in Buffer A (100 mM HEPES, 1 mM EDTA, pH 7.5) at 37°C. Following dissolution, the protein sample was diluted to 2% SDS in Buffer A and split into + hydroxylamine (HA) and control samples. At this step, samples for total protein expression were collected for mass spectrometry. HA was added to the paired samples at a total concentration of 500 mM at neutral pH and incubated at 4°C overnight. Thiopropyl sepharose beads were prewashed 4 times in water, incubated in Buffer B (1% SDS, 100mM HEPES, 1mM EDTA, pH 7.5), then incubated with protein samples for 2 hours at room temperature. The beads were washed five times with Buffer B to remove non-specific contaminants, followed by Buffer B with 50 mM dithiothretol (DTT) to elute previously-palmitoylated protein from the beads. Beads were incubated for 20 minutes at room temperature before supernatant was obtained for mass spectrometric analysis.

#### Mass Spectrometry

Proteomics analyses were performed and analyzed at the Yale Mass Spectrometry (MS) & Proteomics Resource of the W.M. Keck Foundation Biotechnology Resource Laboratory with the support of the Yale/NIDA Neuroproteomics Center. Brain homogenates or synaptosomes were lysed with RIPA buffer containing protease and phosphatase inhibitor cocktail using ultra-sonication. Cellular debris were removed by centrifugation at 16,000g for 10 minutes at 4°C. 150 μL of the supernatant was transferred to a new tube and proteins were precipitated with Chloroform:MeOH:water (100:400:300 μL). The protein pellet was washed three times with cold methanol prior to air drying for 5 minutes. Dried protein pellets were resolubilized with 8 M urea containing 400 mM ammonium bicarbonate. Protein concentration was measured using Nanodrop (Thermo Fisher Scientific; Waltham, MA), and 100 μg of each sample was taken for additional downstream sample preparation. The cysteines within the 100 μg protein samples were reduced with DTT at 37°C for 30 minutes and alkylated with iodoacetamide at room temperature in the dark for 30 minutes. Reduced and alkylated proteins were then digested with Lys-C (1:25 enzyme:protein ratio) overnight, and subsequently with trypsin (1:25 enzyme:protein ratio) for 7 hours at 37°C. Digestion was quenched with 20% trifluoroacetic acid, and samples were desalted using C18 reverse phase macrospin columns (The Nest Group Inc., Southborough, MA). Then, eluted peptides were dried using SpeedVac. Total peptide concentration was determined by nanodrop after reconstitution with 0.1% formic in water. Dilutions were made to ensure that equal total amounts (0.25 μg) were loaded on the column for Liquid Chromatography MS/MS analyses. Retention Calibration Mix (Thermo Fisher Scientific; Waltham, MA) was added equally to all the peptide solutions to be analyzed within the set of comparative samples for system quality control and downstream normalization. LFQ of protein samples was performed on a mass spectrometer (Thermo Scientific Orbitrap Fusion or Thermo Scientific Q-Exactive Plus) connected to a UPLC system (Waters nanoACQUITY) equipped with a Waters Symmetry® C18 180 μm × 20 mm trap column and a 1.7-μm, 75 μm × 250 mm nano ACQUITY UPLC column (35°C). Additional details on UPLC and mass spectrometer conditions can be found in Charkoftaki et al., 2019. The LC-MS/MS data was processed using Progenesis QI Proteomics software (Nonlinear Dynamics, version 4.0), and protein identification was carried out using the Mascot search algorithm (Matrix Science, Boston MA).

#### Pathway analysis

Pathway analyses were conducted with three distinct software platforms – Ingenuity Pathway Analysis (IPA) available from Qiagen, Gene Ontology Resource, and STRING analysis.

#### PPT1 validation

HEK293T cells were plated at 2 X 10^6^ cells per plate on 10 cm^2^ dishes one hour prior to transfection with LCV-mPPT1 or LCV-FUGW using GenePorter3000. After 48 hours of incubation, growth media was removed from the cells and filtered through a 0.22 μm filter. Media was then concentrated using a 10,000 NMWL centrifugal filter at 2900g at 4°C for a total of 40 minutes. Concentrated media was then brought to 5 mM MgCl_2_ and 10 mM Tris HCl, pH 7.2 and protease inhibitors were added. The Acyl RAC procedure was performed as above, except synaptosomes were resuspended at pH 5.5 and divided into 3 parts prior to HA incubation. PPT1 media, GFP media, or HA was then added to synaptosomes during the HA step of Acyl RAC to depalmitoylate proteins. The samples were then subjected to LFQ mass spectrometry as above.

#### Generation of custom PPT1 antibody

Polyclonal antibodies were raised in rabbit against a PPT1 peptide antigen (STLYTEDRLGLKKMDKAGK), and serum was affinity purified (Thermo Fisher Life Custom Antibodies, Life Sciences Solutions).

#### Western blot analyses

SDS-PAGE and western blots were performed using standard procedures. Images and densitometry values (where applicable) were collected using a LI-COR Odyssey imaging system. Western blot signal quantifications are represented as mean ± SEM.

#### Immunocytochemistry (ICC) & confocal microscopy

Primary hippocampal neurons from WT and PPT1 KO mice (P0) were cultured on coverslips, as previously described (Vargas et al., 2014). For lentiviral transductions, lentivirus particles were added to the media at DIV4. At DIV 14, neurons were fixed with 4% buffered paraformaldehyde (PFA) with 4% sucrose, washed in 1X PBS, and blocked in 3% goat serum at room temperature. Neurons were incubated in primary antibodies overnight at 4°C, then in Alexa-conjugated secondaries at 4°C. Coverslips were prepared in antifade mounting medium with DAPI (H-1000 Vectashield) and sealed. Fluorescent images were collected with a Zeiss LSM 800 laser scanning confocal microscope with a 63X oil immersion objective.

#### Plasmid construction and lentiviral preparation

GluA1 Cysteine 323 to Alanine (GluA1.C323A) was generated using QuikChange site-directed mutagenesis (Agilent). GluA1.WT, GluA1.C323A, and TARPγ8 were subcloned into pGEMHE (Addgene). Lentiviruses for hPPT1 were prepared by transfecting HEK293T cells with LCV-hPPT1 and viral constructs. Virus was collected from the media, concentrated, and stored at -80 degrees until use.

#### *Xenopus laevis* oocyte electrophysiology

Oocytes were surgically extracted from anesthetized *Xenopus laevis* frogs and defolliculated with collagenase, as previously described (Salm et al., 2020). Two-electrode voltage-clamp (TEVC) recordings were then performed as previously described (Kim et al., 2010). Briefly, cRNAs were transcribed *in vitro* with a T7 mMessage mMachine (Ambion). Manually selected *X. laevis* oocytes were injected with cRNA of GluA1.WT or GluA1.C323A (100 pg), either with TARPγ8 (100 pg), or alone at a higher concentration (2 ng). Two-electrode voltage-clamp recordings (E_h_ = -70 mV) were taken 2 days after injection at room temperature. Glutamate (1 mM) was bath applied in recording solution (90 mM NaCl, 1.0 mM KCl, 1.5 mM CaCl_2_, 10 mM HEPES, pH 7.4) with cyclothiazide (50 µM) to block desensitization of AMPA receptors. Data are represented as mean ± SEM.

### QUANTIFICATION AND STATISTICAL ANALYSIS

MS protein data were normalized by comparing the abundance of the spike in Pierce Retention Time Calibration mixture among all the samples as well as the sum of squares of spectral count for each replicate. Relative protein-level fold changes were calculated from the sum of all unique, normalized peptide ion abundances for each protein on each run. Only proteins identified by at least two unique peptides were considered. Mass spectrometry data p-values were calculated using a two-tailed t-test (n=3 biological, 3 technical replicates; n=9 total per genotype per experiment). While this does not meet the t-test condition for independence, we proceeded to maximize our hits. We did not use FDR correction for multiple comparisons in our screen to maximize our hits. In addition, we used 1.5-fold up- or down-regulation as cut-offs for the mass spectrometric data. Peptide-level data are represented as percentages or ratios of summed normalized peptide ion abundances, as described in figure legends. Peptide quantifications are represented as mean ± SD.

ICC image analysis was executed in FIJI and performed blinded to genotype. Files were converted to 8-bit images and 50 µM dendritic ROIs were selected for analysis. The “analyze particles” function was used to obtain puncta number and average puncta size for puncta larger than 20-pixel units (mean gray value scale: CADM2, 100-255; SYPH, 75-255). To assess colocalization, Pearson’s correlation coefficient was calculated using the “Coloc 2” function. Data are represented as mean ± SD.

Throughout, p-values were calculated using two-tailed t-tests, and p-values <0.05 were considered significant. All protein and gene names are listed using UniProt nomenclature. Figures were generated in GraphPad Prism 7.0, Adobe Illustrator, and Ingenuity Pathway Analysis.

### KEY RESOURCES TABLE

| REAGENT or RESOURCE | SOURCE | IDENTIFIER |
| --- | --- | --- |
| **Antibodies** |  |  |
| Rabbit polyclonal anti-CSPα (used at 1:5000) | Chemicon | Cat# AB1576; RRID: AB_90794 |
| Rabbit polyclonal anti-endophilin 1 (used at 1:1000) | Synaptic Systems | Cat# 159 002; RRID: AB_887757 |
| Mouse monoclonal anti-SNAP-25 (used at 1:5000) | Covance | Cat# SMI-81; RRID: AB_2315336 |
| Mouse monoclonal anti-β-Actin (used at 1:1000) | GeneTex | Cat# GTX629630; RRID: AB_2728646 |
| Mouse monoclonal anti-dynamin (used at 1:2000) | BD Biosciences | Cat# 610246; RRID: AB_397641 |
| Rabbit polyclonal anti-NrCAM (used at 1:10,000) | Abcam | Cat# ab24344; RRID: AB_448024 |
| Rabbit polyclonal anti-ATP1B2 (used at 1:200) | Thermo Fisher Scientific | Cat# PA5-11852; AB_2857261 |
| Rabbit polyclonal anti-GluA1 (used at 1:1000) | Millipore | Cat# AB1504; RRID: AB_2113602 |
| Rat anti-Syncam2 (used at 1:2000 for ICC) | Gift from the laboratory of Dr. Thomas Biederer | N/A – Custom Antibody |
| Chicken polyclonal anti-MAP2 (used at 1:500 for ICC) | Millipore | Cat# AB5543; RRID: AB_571049 |
| Mouse monoclonal anti-myelin basic protein (MBP) (used at 1:750) | Abcam | Cat# ab62631; RRID: AB_956157 |
| Rabbit polyclonal anti-synaptogyrin 3 (used at 1:1000) | Synaptic Systems | Cat# 103 3030; RRID: AB_2619753 |
| Mouse monoclonal anti-synaptophysin 1 (used at 1:10,000 for WB, 1:500 for ICC) | Synaptic Systems | Cat# 101 011; RRID: AB_887822 |
| Rabbit monoclonal anti-α-Synuclein (used at 1:1000) | Cell Signaling Technology | Cat# 4179; RRID: AB_1904156 |
| Rabbit polyclonal anti-PPT1 (used at 1:200 for ICC) | GeneTex | Cat #: GTX110677;<br>RRID: AB_1951370 |
| Rabbit polyclonal anti-PPT1 (used at 1:50 for WB) | This paper;<br>Thermo Fisher<br>Scientific Life<br>Sciences<br>Solutions | Custom Antibody;<br>Project VH2380 |
| IRDye 680RD goat anti-mouse IgG secondary | LI-COR<br>Biosciences | Cat# 926-68070;<br>RRID: AB_10956588 |
| IRDye 680RD goat anti-rabbit IgG secondary | LI-COR<br>Biosciences | Cat# 926-68071;<br>RRID: AB_10956166 |
| IRDye 800CW goat anti-rabbit IgG secondary | LI-COR<br>Biosciences | Cat# 926-32211;<br>RRID: AB_621843 |
| IRDye 800CW donkey anti-mouse IgG secondary | LI-COR<br>Biosciences | Cat# 926-32212;<br>RRID: AB_621847 |
| Goat anti-rat IgG (H+L) cross-adsorbed, Alexa Fluor 488 | Invitrogen | Cat# A11006; RRID:<br>AB_141373 |
| Goat polyclonal anti-chicken IgY (H+L) cross-adsorbed, Alexa Fluor 633 secondary | Invitrogen | Cat# A21103; RRID:<br>AB_2535756 |
| **Chemicals, Peptides, and Recombinant Proteins** |  |  |
| tris(2-carboxyethyl)phosphine (TCEP) | Thermo Fisher<br>Scientific | Cat# 20490 |
| Hydroxylamine (HA) | Millipore Sigma | Cat# 467804 |
| *N*-Ethylmaleimide (NEM) | Thermo Fisher<br>Scientific | Cat# 23030 |
| Thiopropyl sepharose 4B | Millipore Sigma | Cat# T8512 |
| *d*<sub>5</sub>-*N*-ethylmaleimide (dNEM) | Cambridge<br>Isotope<br>Laboratories | Cat# DLM-6711-10 |
| QuikChange II XL Site-Directed Mutagenesis Kit | Agilent | Cat# 200521 |
| **Deposited Data** |  |  |
| Analyzed mass spectrometry data | This paper | Tables |
| Raw data | ProteomeXchang<br>e Consortium | PXD017270 |
| **Experimental Models: Cell Lines** |  |  |
| HEK 293T cells | American Type<br>Culture Collection | Cat# CRL-3216 |
| **Experimental Models: Organisms/Strains** |  |  |
| *Mus musculus* - B6;129-Ppt1tm1Hof/J | The Jackson<br>Labs | JAX: 004313 |
| *Xenopus laevis* | Nasco | LM00531 |
| **Recombinant DNA** |  |  |
| LCV-hPPT1 | This paper | N/A |
| LCV-FUGW | This paper | N/A |
| pGEMHE-GluA1.WT | This paper | N/A |
| pGEMHE-GluA1.C323A | This paper | N/A |
| pGEMHE-TARPG8 | This paper | N/A |

| **Software and Algorithms** |  |  |
| --- | --- | --- |
| CSS Palm | Ren et al., 2008 | http://csspalm.biocuckoo.org/ |
| Swiss Palm | Blanc et al., 2015 | https://swisspalm.org/ |
| Ingenuity Pathway Analysis | Qiagen | https://www.qiagenbioinformatics.com/products/ingenuity-pathway-analysis/ |
| GraphPad Prism 7 | GraphPad | https://www.graphpad.com/scientific-software/prism/ |
| Progenesis QI Proteomics software V4.0 | Nonlinear Dynamics | http://www.nonlinear.com/progenesis/qi-for-proteomics/download/ |
| Mascot search algorithm | Matrix Science | http://www.matrixscience.com/search_form_select.html |
| WebLogo | Crooks et al., 2004 | https://weblogo.berkeley.edu/ |
| SWISS-MODEL | Bienert et al., 2017 | https://swissmodel.expasy.org |

