## Supplemental Figures & Tables for "Identification of synaptic PPT1 substrates highlight roles of depalmitoylation in disulfide bond formation and synaptic function"

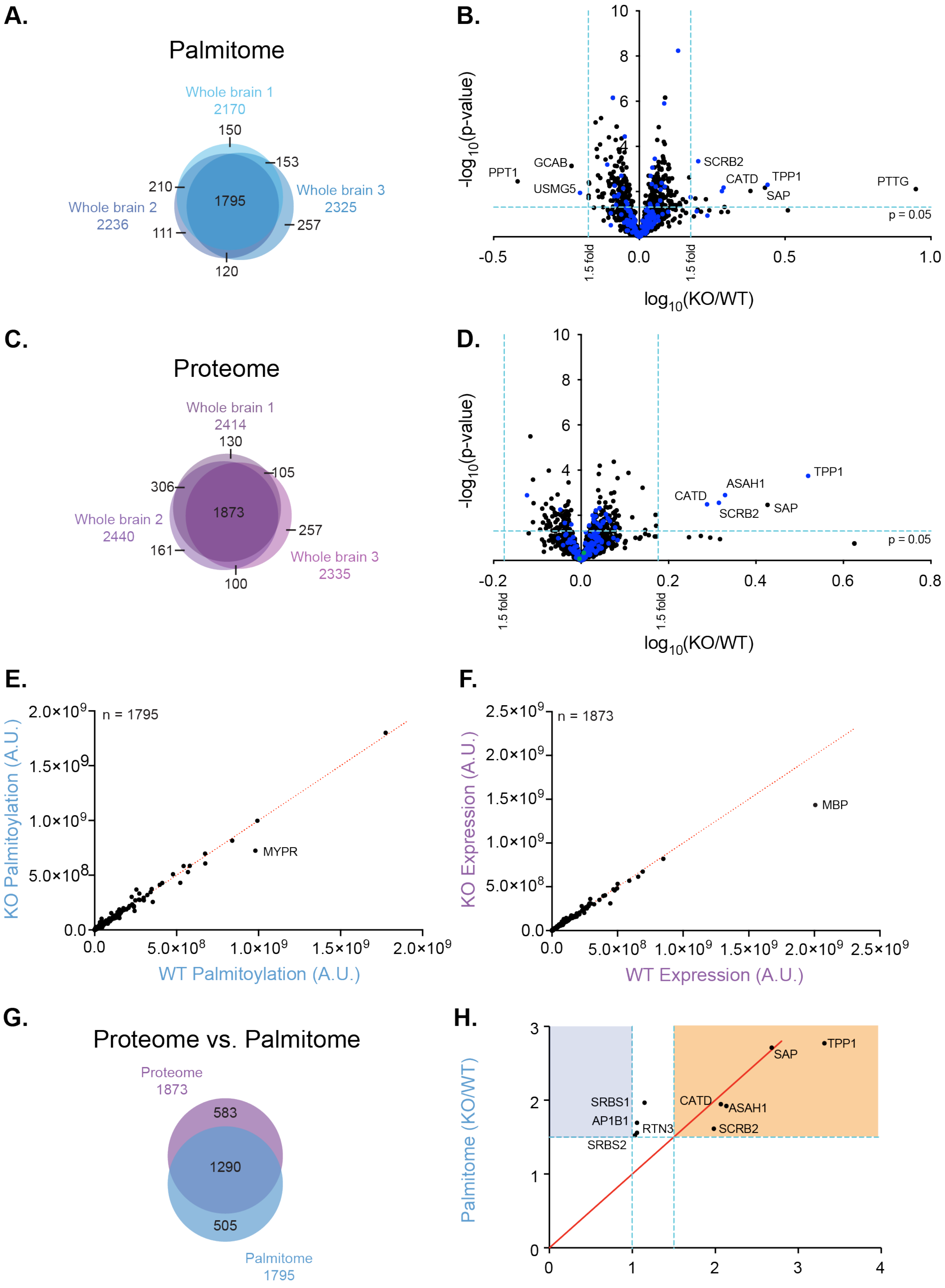

A.
Palmitome
Whole brain 1
2170
Whole brain 2
2236
Whole brain 3
2325
150
153
210
1795
257
111
120
B.
-log10(p-value)
log10(KO/WT)
PPT1
GCAB
USMG5
SCRB2
CATD
TPP1
SAP
PTTG
p = 0.05
1.5 fold
C.
Proteome
Whole brain 1
2414
Whole brain 2
2440
Whole brain 3
2335
130
105
306
1873
257
161
100
D.
-log10(p-value)
log10(KO/WT)
TPP1
ASAH1
CATD
SCRB2
SAP
p = 0.05
1.5 fold
E.
n = 1795
MYPR
KO Palmitoylation (A.U.)
WT Palmitoylation (A.U.)
F.
n = 1873
MBP
KO Expression (A.U.)
WT Expression (A.U.)
G.
Proteome vs. Palmitome
Proteome
1873
583
1290
505
Palmitome
1795
H.
Palmitome (KO/WT)
Proteome (KO/WT)
SAP
TPP1
SRBS1
AP1B1
RTN3
SRBS2
CATD
ASAH1
SCRB2

**Supplemental Figure 1. Generation of WT and PPT1 KO palmitome and proteome expands the repertoire of known palmitoylated proteins in the brain**. A) Venn diagram of 3 independent palmitome experiments exhibits 1795 common proteins. B) Volcano plot of fold change between genotypes (KO/WT) for common proteins with putative synaptic PPT1 substrates in blue (**Figure 2B, Table S2**). 15 palmitoylated proteins are significantly differentially expressed in whole brain (3 decreased, including PPT1; 12 increased; 1.5-fold, $p < 0.05$; blue lines). C) Venn diagram of 3 independent proteome experiments exhibits 1873 common proteins. D) Volcano plot of fold change between genotypes (KO/WT) for common proteins with putative synaptic PPT1 substrates in blue (**Figure 2B, Table S2**). 5 proteins are significantly upregulated (1.5-fold, $p < 0.05$; blue lines). Other depalmitoylating enzymes (green points) do not display compensatory upregulation of protein expression. E) Palmitoylated protein expression was highly correlated between genotypes for palmitome hits ($m = 0.9753 \pm 0.0030$; $R^2 = 0.9857$). F) Protein expression was highly correlated between genotypes for proteome hits ($m = 0.8675 \pm 0.0037$; $R^2 = 0.7711$). When the MBP outlier was removed, this correlation was improved ($m = 0.9802 \pm 0.0023$; $R^2 = 0.9916$). Red lines indicate 1:1 WT to PPT1 KO protein expression ratio. G) Venn diagram of 1290 proteins common between whole brain proteome and palmitome (n = 3 each). H) Protein expression levels compared to palmitoylation levels for significantly changed proteins in palmitome. Proteins in orange region are significantly increased (1.5-fold; $p < 0.05$) in both the proteome and palmitome. Proteins in blue region display decreased or unchanged protein expression and increased palmitoylation. Red line indicates equal expression and palmitoylation levels ($x = y$).

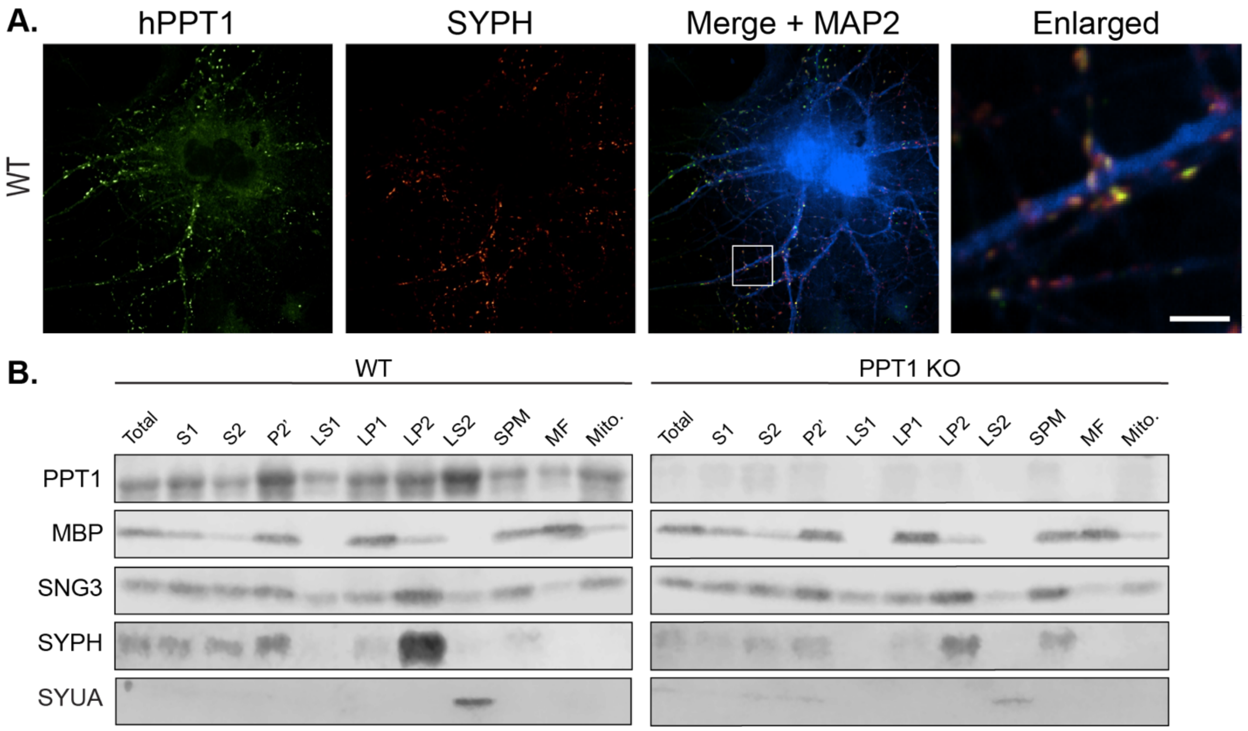


**Supplemental Figure 2**. **PPT1 is expressed in multiple synaptic membrane and cytosolic sub-compartments in neurons.** A) hPPT1 (green) colocalizes with synaptophysin 1 (SYPH; red) in MAP2+ neurites in WT mouse primary neuronal culture (10 μm scale bar). Lentiviral transduction of human PPT1 was performed due to lack of commercially available antibodies that can detect mouse PPT1 by immunocytochemistry (including the custom antibody used for western blotting). B) Endogenous PPT1 is present in all synaptic fractions of a subcellular fractionation of whole brains (8 WT and 8 PPT1 KO mice; age = 2 months). Markers of synaptic sub-compartments are appropriately localized: myelin basic protein (MBP) in myelin fraction (MF); synaptogyrin 3 (SNG3) and synaptophysin 1 (SYPH) in synaptic vesicle enriched fraction (LP2) and synaptic plasma membrane (SPM); α-synuclein (SYUA) in synaptic cytosolic fraction (LS2).

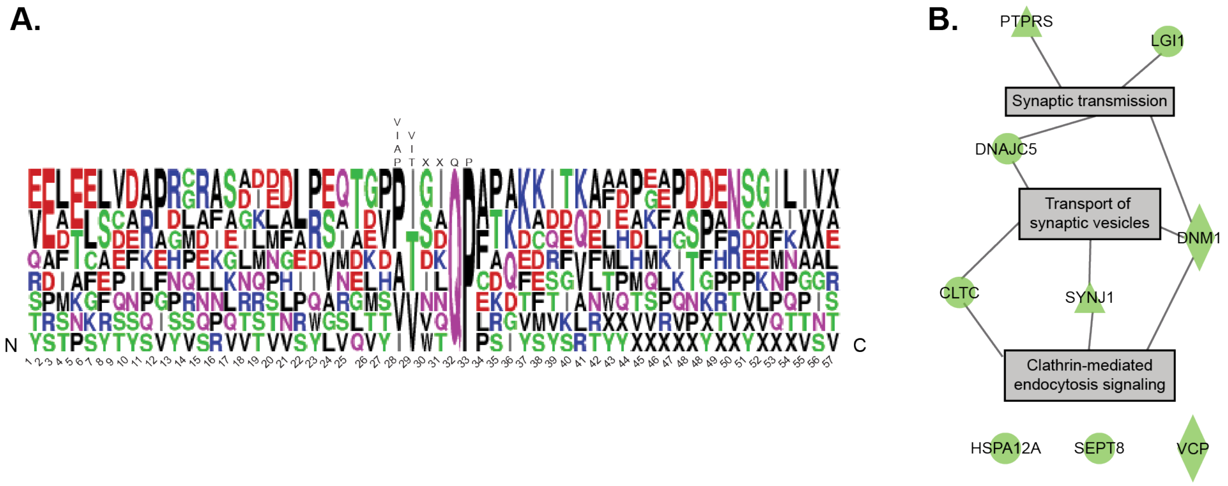


**Supplemental Figure 3**. **Endocytic PPT1 substrates contain the DHHC17 recognition motif.** A) Alignment of DHHC17 recognition motif for 9 endocytic PPT1 substrates with 26 amino acids on either side. The motif is noted above the logo plot. There are no other motifs surrounding the DHHC17 motif. B) Ingenuity pathway analysis of DHHC17 motif-containing proteins identifies clathrin-mediated endocytosis signaling as the top enriched pathway. These pathways account for 6 of the 9 motif-containing proteins.

**Table S1. WT and PPT1 KO synaptic protein expression.** Average KO/WT protein expression ratios for data shown in **Figure 2D**. P-values were calculated using two-tailed t-tests. Blue denotes decreases (<1.5 fold), red denotes increases (>1.5 fold), while black denotes unchanged expression.

| Uniprot ID | Average Ratio (KO/WT) | P value |
|---|---|---|
| PPT1_MOUSE | 0.1324065 | 1.9306E-07 |
| ASAH1_MOUSE | 2.03207633 | 7.1104E-08 |
| CATD_MOUSE | 1.9598571 | 2.0852E-10 |
| CX7A2_MOUSE | 1.89377338 | 0.02926753 |
| GFAP_MOUSE | 1.58759041 | 0.00525781 |
| NDUF4_MOUSE | 1.55884846 | 0.0073044 |
| NU4M_MOUSE | 1.53679812 | 0.01790171 |
| SAP_MOUSE | 3.07387932 | 6.389E-11 |
| SCRB2_MOUSE | 1.87158589 | 0.00153495 |
| TOM7_MOUSE | 1.74901867 | 0.00948743 |
| TPP1_MOUSE | 2.73139373 | 7.4374E-08 |
| 1433B_MOUSE | 0.98583501 | 0.5571686 |
| 1433E_MOUSE | 0.98173666 | 0.47927337 |
| 1433F_MOUSE | 0.92138057 | 0.09023641 |
| 1433G_MOUSE | 0.94088453 | 0.13610869 |
| 1433T_MOUSE | 0.9064798 | 0.05521134 |
| 1433Z_MOUSE | 0.91939427 | 0.03163892 |
| 2A5B_MOUSE | 0.91767961 | 0.17958149 |
| 2A5E_MOUSE | 0.93588692 | 0.20594846 |
| 2A5G_MOUSE | 0.92516276 | 0.27527372 |
| 2AAA_MOUSE | 0.95194269 | 0.21285607 |
| 2ABA_MOUSE | 0.95367966 | 0.22598709 |
| 3HIDH_MOUSE | 1.04327957 | 0.79398386 |
| 4F2_MOUSE | 1.02898308 | 0.66595994 |
| 68MP_MOUSE | 1.2916347 | 0.1393345 |
| 6PGD_MOUSE | 0.98481313 | 0.75742688 |
| 6PGL_MOUSE | 0.99064213 | 0.75351609 |
| A4_MOUSE | 0.91521678 | 0.11759947 |
| AACS_MOUSE | 0.90033075 | 0.14738068 |
| AAK1_MOUSE | 0.98735689 | 0.64757685 |
| AATC_MOUSE | 0.99724389 | 0.88741276 |
| AATM_MOUSE | 1.01539692 | 0.81680797 |
| ABCB7_MOUSE | 1.03437721 | 0.72455275 |
| ABCB8_MOUSE | 0.92454716 | 0.04636432 |

| | | |
|---|---|---|
| ABCD3_MOUSE | 1.05480501 | 0.51271416 |
| ABD12_MOUSE | 1.01144992 | 0.72044301 |
| ABHDA_MOUSE | 1.11382811 | 0.31998147 |
| ABHGA_MOUSE | 0.94834703 | 0.23558947 |
| ABI1_MOUSE | 1.00358768 | 0.84630625 |
| ABI2_MOUSE | 0.89302221 | 0.16879112 |
| ABLM1_MOUSE | 0.99187713 | 0.62553276 |
| ABLM2_MOUSE | 0.98401907 | 0.70335008 |
| ABR_MOUSE | 0.98729854 | 0.6186061 |
| ACACA_MOUSE | 0.94863088 | 0.29824119 |
| ACAD8_MOUSE | 0.95668948 | 0.37781067 |
| ACAD9_MOUSE | 0.9803481 | 0.49322535 |
| ACADL_MOUSE | 1.11932528 | 0.07927052 |
| ACADM_MOUSE | 1.04129721 | 0.52577499 |
| ACADS_MOUSE | 1.0463713 | 0.82805935 |
| ACADV_MOUSE | 1.03872925 | 0.38873496 |
| ACBD6_MOUSE | 0.89777266 | 0.10916865 |
| ACBG1_MOUSE | 0.9736729 | 0.50146583 |
| ACBP_MOUSE | 0.94736514 | 0.34919227 |
| ACDSB_MOUSE | 1.02078461 | 0.76794946 |
| ACES_MOUSE | 0.96364846 | 0.54106011 |
| ACLY_MOUSE | 0.98858338 | 0.64280925 |
| ACO11_MOUSE | 0.97019004 | 0.46162536 |
| ACO13_MOUSE | 1.07276753 | 0.30443162 |
| ACOC_MOUSE | 1.02463082 | 0.62056124 |
| ACON_MOUSE | 1.02583766 | 0.81032263 |
| ACOT9_MOUSE | 1.05205746 | 0.23715939 |
| ACOX1_MOUSE | 1.35749642 | 0.01158117 |
| ACPM_MOUSE | 1.15626307 | 0.05123628 |
| ACSF2_MOUSE | 1.19753129 | 0.02615009 |
| ACSF3_MOUSE | 1.02599796 | 0.83642539 |
| ACSL1_MOUSE | 1.03678754 | 0.49813489 |
| ACSL3_MOUSE | 1.00424691 | 0.83509572 |
| ACSL6_MOUSE | 1.04949839 | 0.22151683 |
| ACTB_MOUSE | 0.92146447 | 0.09707842 |
| ACTC_MOUSE | 1.08163565 | 0.54517998 |
| ACTG_MOUSE | 0.94785731 | 0.3123261 |
| ACTN1_MOUSE | 1.04246345 | 0.58964527 |
| ACTN2_MOUSE | 1.00436901 | 0.9617644 |

| | | |
|---|---|---|
| ACTN4_MOUSE | 0.89503777 | 0.0658385 |
| ACTY_MOUSE | 1.01662349 | 0.64068865 |
| ACTZ_MOUSE | 1.00274656 | 0.90438894 |
| ACY2_MOUSE | 0.75121584 | 0.00017773 |
| ACYP1_MOUSE | 0.95202532 | 0.200298 |
| ADA11_MOUSE | 1.01879609 | 0.85015647 |
| ADA22_MOUSE | 1.04358024 | 0.76282963 |
| ADA23_MOUSE | 0.99141222 | 0.6482934 |
| ADCK1_MOUSE | 0.99950778 | 0.98478884 |
| ADCY5_MOUSE | 0.93989396 | 0.1990365 |
| ADCY9_MOUSE | 0.97061256 | 0.48192215 |
| ADDA_MOUSE | 1.00947852 | 0.86514912 |
| ADDB_MOUSE | 1.04320106 | 0.21928638 |
| ADDG_MOUSE | 0.98292048 | 0.55179324 |
| ADHX_MOUSE | 0.95361516 | 0.0875614 |
| ADK_MOUSE | 1.02157436 | 0.87405279 |
| ADPRH_MOUSE | 1.03361479 | 0.68142929 |
| ADT1_MOUSE | 1.00478358 | 0.92765781 |
| ADT2_MOUSE | 0.99952531 | 0.75706883 |
| AFAD_MOUSE | 1.0307234 | 0.78500273 |
| AFG32_MOUSE | 0.99103875 | 0.6276594 |
| AGAP2_MOUSE | 0.98011748 | 0.47876885 |
| AGAP3_MOUSE | 0.99019542 | 0.68336469 |
| AGFG1_MOUSE | 1.00131728 | 0.92463358 |
| AGK_MOUSE | 1.08320684 | 0.62013365 |
| AGM1_MOUSE | 0.88484876 | 0.0086762 |
| AGRL1_MOUSE | 1.01473453 | 0.78894441 |
| AGRL3_MOUSE | 0.94233571 | 0.25288306 |
| AHSA1_MOUSE | 1.03581732 | 0.67341969 |
| AIFM1_MOUSE | 0.97016079 | 0.40862561 |
| AINX_MOUSE | 1.15678731 | 0.12331887 |
| AK1A1_MOUSE | 0.98262512 | 0.52720552 |
| AKA12_MOUSE | 0.88688352 | 0.23433627 |
| AKA7A_MOUSE | 1.01473012 | 0.82872316 |
| AKAP5_MOUSE | 1.01304215 | 0.95406776 |
| AKCL2_MOUSE | 1.01667196 | 0.94332204 |
| AL1A1_MOUSE | 1.01979462 | 0.79180123 |
| AL1B1_MOUSE | 0.99870187 | 0.77295367 |
| AL1L1_MOUSE | 1.033779 | 0.55294913 |

| | | |
|---|---|---|
| AL3A2_MOUSE | 0.96422921 | 0.43583518 |
| AL4A1_MOUSE | 1.07133763 | 0.26487501 |
| AL7A1_MOUSE | 1.11255825 | 0.11033686 |
| AL9A1_MOUSE | 1.04619003 | 0.33937178 |
| ALBU_MOUSE | 0.73056189 | 0.00300301 |
| ALDH2_MOUSE | 1.01015857 | 0.8397427 |
| ALDOA_MOUSE | 1.01221097 | 0.9584375 |
| ALDOC_MOUSE | 1.05019299 | 0.51227259 |
| ALDR_MOUSE | 0.97163715 | 0.36689111 |
| ALG2_MOUSE | 0.94537116 | 0.35234149 |
| ALR_MOUSE | 0.9387795 | 0.46413448 |
| AMD_MOUSE | 1.09569923 | 0.22764743 |
| AMPB_MOUSE | 0.9187945 | 0.13921191 |
| AMPD2_MOUSE | 1.04600762 | 0.52170757 |
| AMPH_MOUSE | 0.8869798 | 0.07170537 |
| AMPL_MOUSE | 1.08568661 | 0.23477562 |
| AMRP_MOUSE | 0.95903601 | 0.41822429 |
| AN32A_MOUSE | 0.86474585 | 0.03681287 |
| ANK1_MOUSE | 1.04075792 | 0.81146394 |
| ANK2_MOUSE | 1.00200971 | 0.84572697 |
| ANK3_MOUSE | 1.01682377 | 0.85371332 |
| ANM5_MOUSE | 0.90454203 | 0.07330723 |
| ANS1B_MOUSE | 0.9868218 | 0.55807941 |
| ANXA3_MOUSE | 0.96377447 | 0.57577386 |
| ANXA5_MOUSE | 1.06104112 | 0.0196424 |
| ANXA6_MOUSE | 1.02629433 | 0.33126602 |
| ANXA7_MOUSE | 0.90795603 | 0.21982937 |
| AOFA_MOUSE | 1.03214509 | 0.27253897 |
| AOFB_MOUSE | 1.13364749 | 0.10433588 |
| AP180_MOUSE | 1.01811375 | 0.86287515 |
| AP1B1_MOUSE | 1.0048851 | 0.85289081 |
| AP1G1_MOUSE | 1.02365273 | 0.58922965 |
| AP1M1_MOUSE | 1.00142115 | 0.91196347 |
| AP2A1_MOUSE | 0.9960234 | 0.77657554 |
| AP2A2_MOUSE | 1.00618472 | 0.9535714 |
| AP2B1_MOUSE | 0.99987012 | 0.83752095 |
| AP2M1_MOUSE | 0.98002538 | 0.58189395 |
| AP2S1_MOUSE | 0.95923402 | 0.36236442 |
| AP3B2_MOUSE | 0.97649273 | 0.38641444 |

| | | |
|---|---|---|
| AP3D1_MOUSE | 1.03504899 | 0.44215605 |
| AP3M2_MOUSE | 0.92518115 | 0.25674898 |
| APEH_MOUSE | 1.02170056 | 0.86370785 |
| APMAP_MOUSE | 1.15227943 | 0.07341667 |
| APOE_MOUSE | 1.0812543 | 0.56552118 |
| AQP4_MOUSE | 1.28825636 | 0.00808742 |
| ARBK1_MOUSE | 1.0158497 | 0.9584319 |
| ARC1A_MOUSE | 1.0531272 | 0.6923467 |
| ARF1_MOUSE | 1.0347915 | 0.83563863 |
| ARF5_MOUSE | 1.06912599 | 0.44875724 |
| ARF6_MOUSE | 1.14332595 | 0.18694691 |
| ARFG1_MOUSE | 0.96255979 | 0.33486025 |
| ARHG2_MOUSE | 0.99281239 | 0.67501943 |
| ARHG7_MOUSE | 1.27181402 | 0.418026 |
| ARK72_MOUSE | 1.08551521 | 0.16493593 |
| ARL3_MOUSE | 1.04119279 | 0.60250279 |
| ARL6_MOUSE | 0.95776521 | 0.33337084 |
| ARL8A_MOUSE | 1.37307239 | 0.20876189 |
| ARL8B_MOUSE | 1.18856673 | 2.5796E-07 |
| ARLY_MOUSE | 0.99671467 | 0.86622609 |
| ARM10_MOUSE | 0.97750237 | 0.47536565 |
| ARMC1_MOUSE | 1.10962526 | 0.34182471 |
| ARP10_MOUSE | 0.98782761 | 0.70727122 |
| ARP2_MOUSE | 1.00865463 | 0.8602501 |
| ARP3_MOUSE | 1.02234308 | 0.85924876 |
| ARP3B_MOUSE | 0.97602424 | 0.38563952 |
| ARP5L_MOUSE | 0.97062534 | 0.32796031 |
| ARPC2_MOUSE | 1.05881648 | 0.64536015 |
| ARPC3_MOUSE | 0.98864959 | 0.60010269 |
| ARPC4_MOUSE | 1.00251771 | 0.97396826 |
| ARPC5_MOUSE | 1.00454984 | 0.88192802 |
| ARRB1_MOUSE | 1.17781089 | 0.09450326 |
| ASAP1_MOUSE | 1.02446025 | 0.44293523 |
| ASGL1_MOUSE | 0.98519099 | 0.50916938 |
| ASNA_MOUSE | 1.04192616 | 0.38204748 |
| ASSY_MOUSE | 1.11586184 | 0.02223865 |
| ASTN1_MOUSE | 0.9876944 | 0.77449576 |
| AT1A1_MOUSE | 0.9595546 | 0.31795481 |
| AT1A2_MOUSE | 1.06757864 | 0.4977135 |

| | | |
|---|---|---|
| AT1A3_MOUSE | 1.04236915 | 0.69483968 |
| AT1A4_MOUSE | 1.04424697 | 0.91097065 |
| AT1B1_MOUSE | 0.98238956 | 0.53580402 |
| AT1B2_MOUSE | 1.04328394 | 0.47105468 |
| AT1B3_MOUSE | 0.99751792 | 0.84914292 |
| AT2A2_MOUSE | 1.0596817 | 0.16985843 |
| AT2B1_MOUSE | 1.0270506 | 0.86560869 |
| AT2B2_MOUSE | 1.01730849 | 0.99801045 |
| AT2B4_MOUSE | 0.99535077 | 0.73609152 |
| AT5F1_MOUSE | 1.00319234 | 0.87444768 |
| AT8A1_MOUSE | 0.96619097 | 0.34216116 |
| ATAD1_MOUSE | 1.04746733 | 0.45438604 |
| ATAD3_MOUSE | 0.98255806 | 0.45504613 |
| ATCAY_MOUSE | 1.05933576 | 0.39736066 |
| ATIF1_MOUSE | 1.05066462 | 0.6879365 |
| ATLA1_MOUSE | 0.97194914 | 0.4438328 |
| ATOX1_MOUSE | 0.93376963 | 0.15060377 |
| ATP5E_MOUSE | 1.00809842 | 0.99806239 |
| ATP5H_MOUSE | 0.92005157 | 0.09611631 |
| ATP5I_MOUSE | 0.97613267 | 0.42718863 |
| ATP5L_MOUSE | 1.09959222 | 0.18266955 |
| ATP5S_MOUSE | 1.03155476 | 0.8118487 |
| ATPA_MOUSE | 0.99393389 | 0.72710112 |
| ATPB_MOUSE | 1.0172077 | 0.86278579 |
| ATPD_MOUSE | 0.99637597 | 0.93358862 |
| ATPG_MOUSE | 1.00122588 | 0.77843614 |
| ATPK_MOUSE | 0.98789021 | 0.60452132 |
| ATPO_MOUSE | 1.05952768 | 0.27717713 |
| ATX10_MOUSE | 0.98139324 | 0.57779581 |
| AUHM_MOUSE | 0.97972141 | 0.62492621 |
| AUXI_MOUSE | 0.98502072 | 0.58811037 |
| AVL9_MOUSE | 0.93631189 | 0.12373438 |
| B2L13_MOUSE | 1.06604817 | 0.93140479 |
| BACH_MOUSE | 0.99629742 | 0.82209133 |
| BAG5_MOUSE | 0.89824369 | 0.17397646 |
| BAG6_MOUSE | 0.92829455 | 0.32757858 |
| BAIP2_MOUSE | 0.91716911 | 0.05178492 |
| BAP31_MOUSE | 1.09602585 | 0.48593776 |
| BASI_MOUSE | 0.92651477 | 0.10129023 |

| | | |
|---|---|---|
| BASP1_MOUSE | 0.8743612 | 0.19000108 |
| BCAS1_MOUSE | 0.90566058 | 0.05902413 |
| BCAS3_MOUSE | 0.96906907 | 0.34555007 |
| BCAT1_MOUSE | 1.07346554 | 0.19801492 |
| BCS1_MOUSE | 1.13379284 | 0.20483242 |
| BDH_MOUSE | 0.96620927 | 0.33887951 |
| BGAL_MOUSE | 0.93461801 | 0.2449892 |
| BIG3_MOUSE | 0.89871907 | 0.08650094 |
| BIN1_MOUSE | 0.90107359 | 0.04818179 |
| BLMH_MOUSE | 0.92873827 | 0.10889336 |
| BOLA1_MOUSE | 0.94794213 | 0.50879243 |
| BORG4_MOUSE | 0.93035347 | 0.18178535 |
| BPHL_MOUSE | 1.04955425 | 0.60709743 |
| BPNT1_MOUSE | 0.99460767 | 0.69089273 |
| BRNP1_MOUSE | 1.29594192 | 0.08909956 |
| BRSK2_MOUSE | 1.0463491 | 0.855304 |
| BSN_MOUSE | 1.00353359 | 0.83568539 |
| BTBDH_MOUSE | 1.212659 | 0.46128881 |
| C170B_MOUSE | 1.15183827 | 0.12726058 |
| C1QBP_MOUSE | 1.0023987 | 0.92550211 |
| C1TC_MOUSE | 1.00011439 | 0.72533191 |
| C1TM_MOUSE | 0.98036808 | 0.48750339 |
| C2C2L_MOUSE | 1.0179609 | 0.8488582 |
| C560_MOUSE | 1.25260395 | 0.03019851 |
| CA198_MOUSE | 1.15048926 | 0.74403633 |
| CA2D1_MOUSE | 0.95154417 | 0.26865256 |
| CA2D2_MOUSE | 1.02906302 | 0.94937902 |
| CA2D3_MOUSE | 1.01430167 | 0.95635947 |
| CAB39_MOUSE | 0.9523452 | 0.10933287 |
| CAC1A_MOUSE | 0.98578988 | 0.63522475 |
| CAC1B_MOUSE | 1.04865125 | 0.32909221 |
| CAC1E_MOUSE | 0.92656924 | 0.07838562 |
| CACB4_MOUSE | 0.96890641 | 0.45065068 |
| CACP_MOUSE | 1.04455971 | 0.36116959 |
| CAD10_MOUSE | 0.95799348 | 0.45151004 |
| CAD13_MOUSE | 1.08840294 | 0.29656473 |
| CADH2_MOUSE | 0.92883115 | 0.14238845 |
| CADM1_MOUSE | 0.99888376 | 0.92261315 |
| CADM2_MOUSE | 0.98822666 | 0.55928359 |

| | | |
|---|---|---|
| CADM3_MOUSE | 0.96229727 | 0.23162547 |
| CADM4_MOUSE | 0.99921341 | 0.74677347 |
| CAH2_MOUSE | 0.9190263 | 0.15752963 |
| CAH4_MOUSE | 0.94563712 | 0.33906131 |
| CALB1_MOUSE | 0.94978381 | 0.12675795 |
| CALB2_MOUSE | 0.84185513 | 0.00569191 |
| CALR_MOUSE | 1.1544788 | 0.0004207 |
| CALU_MOUSE | 0.91843886 | 0.28027337 |
| CALX_MOUSE | 0.9622982 | 0.36377171 |
| CAMKV_MOUSE | 1.00538188 | 0.90275622 |
| CAN2_MOUSE | 1.2112102 | 0.67318094 |
| CAN5_MOUSE | 0.96056614 | 0.36313268 |
| CANB1_MOUSE | 0.86871042 | 0.01601081 |
| CAND1_MOUSE | 0.95910474 | 0.16247384 |
| CAP1_MOUSE | 0.89514036 | 0.06809104 |
| CAP2_MOUSE | 0.98840445 | 0.7293977 |
| CAPS1_MOUSE | 0.98369221 | 0.58325628 |
| CAPS2_MOUSE | 0.98320306 | 0.67393579 |
| CAPZB_MOUSE | 1.00533049 | 0.73747021 |
| CATA_MOUSE | 1.04312494 | 0.64012675 |
| CATB_MOUSE | 0.93026251 | 0.25469791 |
| CAZA2_MOUSE | 0.97450427 | 0.24086604 |
| CBPE_MOUSE | 1.05094231 | 0.23541524 |
| CBR1_MOUSE | 0.97982834 | 0.58748925 |
| CBR3_MOUSE | 0.93294688 | 0.28053853 |
| CBR4_MOUSE | 1.0835083 | 0.09933198 |
| CC127_MOUSE | 1.5830901 | 0.44604952 |
| CC177_MOUSE | 0.90653937 | 0.17033253 |
| CC50A_MOUSE | 1.01095507 | 0.83571496 |
| CCD22_MOUSE | 0.82714551 | 0.03377642 |
| CCD51_MOUSE | 1.14015917 | 0.36030134 |
| CCD58_MOUSE | 0.84892381 | 0.01618713 |
| CCG8_MOUSE | 0.99124283 | 0.63643284 |
| CCHL_MOUSE | 1.01307138 | 0.96567431 |
| CCNY_MOUSE | 0.92286009 | 0.13024897 |
| CD166_MOUSE | 1.05170851 | 0.60211882 |
| CD47_MOUSE | 0.96876125 | 0.36990625 |
| CD81_MOUSE | 1.03479067 | 0.93953981 |
| CDC37_MOUSE | 1.00944186 | 0.87190281 |

| | | |
|---|---|---|
| CDC42_MOUSE | 1.16481882 | 0.07457434 |
| CDK14_MOUSE | 1.03842199 | 0.66498252 |
| CDK5_MOUSE | 0.94135374 | 0.18021069 |
| CDS2_MOUSE | 1.06361652 | 0.66715341 |
| CDV3_MOUSE | 0.84562167 | 0.00777649 |
| CE170_MOUSE | 0.95359754 | 0.19655359 |
| CH10_MOUSE | 1.01405085 | 0.7927479 |
| CH60_MOUSE | 1.05026896 | 0.44901962 |
| CHM4B_MOUSE | 0.88615465 | 0.07725902 |
| CHP1_MOUSE | 0.97361276 | 0.59807413 |
| CHRD1_MOUSE | 1.11581416 | 0.44098761 |
| CI172_MOUSE | 0.86262242 | 0.07564175 |
| CISD1_MOUSE | 0.99149775 | 0.63514151 |
| CISY_MOUSE | 1.01336742 | 0.85909404 |
| CK054_MOUSE | 0.88981071 | 0.03886698 |
| CKAP5_MOUSE | 0.93671912 | 0.00571075 |
| CLAP1_MOUSE | 1.01745178 | 0.89076905 |
| CLAP2_MOUSE | 1.02172874 | 0.88773472 |
| CLCA_MOUSE | 0.84338981 | 0.013501 |
| CLCB_MOUSE | 0.82982504 | 0.00669776 |
| CLCN3_MOUSE | 1.29392742 | 0.22838283 |
| CLD11_MOUSE | 0.81911986 | 0.01197204 |
| CLH1_MOUSE | 0.99784733 | 0.80263571 |
| CLIC4_MOUSE | 0.93548445 | 0.17646921 |
| CLIP2_MOUSE | 0.94767997 | 0.33256559 |
| CLPB_MOUSE | 1.00440141 | 0.887557 |
| CLPP_MOUSE | 1.07192347 | 0.51639365 |
| CLUS_MOUSE | 1.08039679 | 0.37687686 |
| CLYBL_MOUSE | 1.07281643 | 0.04874069 |
| CMC1_MOUSE | 1.03860837 | 0.59507996 |
| CMPK2_MOUSE | 1.08749447 | 0.32077969 |
| CMTD1_MOUSE | 1.34726272 | 0.07243413 |
| CN166_MOUSE | 0.91843793 | 0.19818151 |
| CN37_MOUSE | 0.92125707 | 0.39967553 |
| CNDP2_MOUSE | 0.9815315 | 0.54505329 |
| CNKR2_MOUSE | 0.920793 | 0.01017472 |
| CNRP1_MOUSE | 1.03108139 | 0.83873952 |
| CNTFR_MOUSE | 1.37733168 | 0.01384741 |
| CNTN1_MOUSE | 0.98686495 | 0.60633784 |

| | | |
|---|---|---|
| CNTN2_MOUSE | 1.03331635 | 0.70007104 |
| CNTP1_MOUSE | 0.97170666 | 0.46900266 |
| CNTP2_MOUSE | 1.01686306 | 0.89940382 |
| COA3_MOUSE | 0.98716024 | 0.54723272 |
| COASY_MOUSE | 1.15434533 | 0.03977472 |
| COF1_MOUSE | 0.91900253 | 0.03557167 |
| COF2_MOUSE | 0.94253162 | 0.09981511 |
| COQ3_MOUSE | 0.90916542 | 0.05388269 |
| COQ7_MOUSE | 1.04636963 | 0.48710164 |
| COQ9_MOUSE | 0.96062365 | 0.15531676 |
| COR1A_MOUSE | 0.9190303 | 0.16189404 |
| COR1B_MOUSE | 1.04250741 | 0.81872739 |
| COR1C_MOUSE | 0.99253369 | 0.70772359 |
| COR2B_MOUSE | 1.03678945 | 0.42309674 |
| COTL1_MOUSE | 1.02943345 | 0.07583358 |
| COX1_MOUSE | 1.17569611 | 0.04054381 |
| COX2_MOUSE | 1.0305403 | 0.50737917 |
| COX41_MOUSE | 0.97846337 | 0.46640039 |
| COX5A_MOUSE | 0.84849241 | 0.00315668 |
| COX5B_MOUSE | 0.94679574 | 0.15242048 |
| COX6C_MOUSE | 1.08279404 | 0.37027397 |
| CP46A_MOUSE | 1.04310061 | 0.53548127 |
| CPLX1_MOUSE | 0.81098979 | 0.00115705 |
| CPLX2_MOUSE | 0.91795691 | 0.08469109 |
| CPNE4_MOUSE | 0.99782195 | 0.84431327 |
| CPNE5_MOUSE | 0.98856654 | 0.58965193 |
| CPNE6_MOUSE | 1.02713074 | 0.89167052 |
| CPNS1_MOUSE | 0.98284553 | 0.57231254 |
| CPT1A_MOUSE | 1.06855799 | 0.19259065 |
| CPT2_MOUSE | 1.11046303 | 0.07564793 |
| CRIP2_MOUSE | 0.94383489 | 0.19591961 |
| CRK_MOUSE | 0.93457824 | 0.25668922 |
| CRKL_MOUSE | 0.94071745 | 0.25477558 |
| CRYAB_MOUSE | 0.97876345 | 0.63069604 |
| CRYM_MOUSE | 1.05918965 | 0.51321635 |
| CSDE1_MOUSE | 0.99040543 | 0.57757061 |
| CSK21_MOUSE | 0.99106267 | 0.71192639 |
| CSK22_MOUSE | 1.08951809 | 0.45089363 |
| CSK2B_MOUSE | 0.95229064 | 0.4786563 |

| | | |
|---|---|---|
| CSKI1_MOUSE | 0.98333622 | 0.58862239 |
| CSKP_MOUSE | 1.07619916 | 0.00986741 |
| CSN1_MOUSE | 1.05542673 | 0.16096471 |
| CSN2_MOUSE | 0.95422923 | 0.23502523 |
| CSN3_MOUSE | 0.98074366 | 0.56895981 |
| CSN4_MOUSE | 0.88975204 | 0.06950214 |
| CSN5_MOUSE | 0.98075794 | 0.52192456 |
| CSN6_MOUSE | 0.93798092 | 0.0997927 |
| CSN7A_MOUSE | 1.13906556 | 0.18974143 |
| CSN8_MOUSE | 0.92506182 | 0.14692901 |
| CSPG2_MOUSE | 0.99899671 | 0.78582444 |
| CSPG5_MOUSE | 0.99269714 | 0.83121483 |
| CSRP1_MOUSE | 1.01129448 | 0.97490544 |
| CTBP1_MOUSE | 1.00547639 | 0.95769061 |
| CTL1_MOUSE | 1.03967704 | 0.82031314 |
| CTL2_MOUSE | 1.14361577 | 0.00165209 |
| CTNA2_MOUSE | 1.01094537 | 0.88083354 |
| CTNB1_MOUSE | 1.05338736 | 0.5272842 |
| CTND1_MOUSE | 1.02841053 | 0.51018581 |
| CTND2_MOUSE | 1.02757612 | 0.98433427 |
| CTRO_MOUSE | 1.03887563 | 0.39144788 |
| CTTB2_MOUSE | 1.02500665 | 0.88456958 |
| CUL1_MOUSE | 1.11502018 | 0.33760585 |
| CUL2_MOUSE | 1.09130302 | 0.05861195 |
| CUL3_MOUSE | 0.96472128 | 0.30651475 |
| CUL5_MOUSE | 0.98430693 | 0.57982141 |
| CX6B1_MOUSE | 0.9192662 | 0.22444627 |
| CXA1_MOUSE | 1.2141345 | 0.00626056 |
| CY1_MOUSE | 1.01025959 | 0.99007648 |
| CYB5_MOUSE | 0.96179705 | 0.34467179 |
| CYB5B_MOUSE | 1.15539448 | 0.4025798 |
| CYBP_MOUSE | 1.07062605 | 0.33654221 |
| CYC_MOUSE | 0.89801836 | 0.28862416 |
| CYFP1_MOUSE | 1.05671175 | 0.369241 |
| CYFP2_MOUSE | 1.01266061 | 0.9889635 |
| CYTB_MOUSE | 1.17074555 | 0.08209781 |
| CYTC_MOUSE | 1.04549651 | 0.70773885 |
| CYTSB_MOUSE | 0.9534108 | 0.32053881 |
| D39U1_MOUSE | 1.07272007 | 0.63741409 |

| | | |
|---|---|---|
| DAAM1_MOUSE | 0.99471311 | 0.59917324 |
| DBNL_MOUSE | 0.90593185 | 0.07662524 |
| DC1I1_MOUSE | 1.06843691 | 0.42622884 |
| DC1I2_MOUSE | 0.98030454 | 0.60169112 |
| DC1L1_MOUSE | 0.95942033 | 0.15816691 |
| DC1L2_MOUSE | 0.94577638 | 0.18708307 |
| DCE1_MOUSE | 1.01945279 | 0.89450169 |
| DCE2_MOUSE | 1.05164401 | 0.24559098 |
| DCLK1_MOUSE | 1.02631587 | 0.72575584 |
| DCNL1_MOUSE | 0.93018092 | 0.45316203 |
| DCTN1_MOUSE | 0.97410227 | 0.3428325 |
| DCTN2_MOUSE | 0.8723489 | 0.04879818 |
| DCTN3_MOUSE | 0.96010336 | 0.32108891 |
| DCTN4_MOUSE | 0.9120714 | 0.02269415 |
| DDAH1_MOUSE | 1.00657171 | 0.79241971 |
| DDAH2_MOUSE | 0.89077874 | 0.01788754 |
| DDB1_MOUSE | 1.03718516 | 0.31974592 |
| DDC_MOUSE | 0.94106925 | 0.25370814 |
| DDX1_MOUSE | 1.02867098 | 0.73475988 |
| DDX3L_MOUSE | 0.96044118 | 0.35035191 |
| DDX6_MOUSE | 0.84713051 | 0.02090482 |
| DECR_MOUSE | 1.12356258 | 0.05415661 |
| DEMA_MOUSE | 0.99371961 | 0.82245602 |
| DEST_MOUSE | 0.96836695 | 0.46617128 |
| DGKB_MOUSE | 1.03468591 | 0.71279534 |
| DHB11_MOUSE | 1.26549881 | 0.43127931 |
| DHB12_MOUSE | 1.10137074 | 0.13365641 |
| DHB4_MOUSE | 1.04702299 | 0.34573986 |
| DHB8_MOUSE | 1.00397809 | 0.88354124 |
| DHE3_MOUSE | 1.08900467 | 0.18567775 |
| DHPR_MOUSE | 1.02703101 | 0.97977501 |
| DHRS1_MOUSE | 1.26654408 | 0.00214101 |
| DHRS4_MOUSE | 1.12242795 | 0.02608375 |
| DHSO_MOUSE | 0.91720589 | 0.18594359 |
| DIC_MOUSE | 1.04545093 | 0.49344677 |
| DIP2A_MOUSE | 0.99926077 | 0.68799539 |
| DIP2B_MOUSE | 1.00294245 | 0.8818966 |
| DIRA2_MOUSE | 0.98643459 | 0.70449186 |
| DJC11_MOUSE | 1.03453362 | 0.61718016 |

| | | |
|---|---|---|
| DLDH_MOUSE | 0.99575126 | 0.78047943 |
| DLG1_MOUSE | 0.9963699 | 0.68242344 |
| DLG2_MOUSE | 0.95438399 | 0.34617645 |
| DLG3_MOUSE | 0.99918288 | 0.6537367 |
| DLG4_MOUSE | 0.97712365 | 0.51314291 |
| DLGP1_MOUSE | 0.87991575 | 0.01500004 |
| DLGP2_MOUSE | 0.96138788 | 0.21621455 |
| DLGP3_MOUSE | 0.96268039 | 0.23103275 |
| DLGP4_MOUSE | 0.94903038 | 0.07619281 |
| DLRB1_MOUSE | 0.94692853 | 0.34518253 |
| DMXL2_MOUSE | 0.9836664 | 0.55030164 |
| DNJA1_MOUSE | 1.0849886 | 0.23511034 |
| DNJA2_MOUSE | 1.01726057 | 0.99256824 |
| DNJA3_MOUSE | 0.98547798 | 0.58687236 |
| DNJC5_MOUSE | 0.8396206 | 0.01771918 |
| DNM1L_MOUSE | 0.99609404 | 0.75402022 |
| DNPEP_MOUSE | 0.98793225 | 0.68215435 |
| DOCK3_MOUSE | 1.82859663 | 0.46020798 |
| DOPD_MOUSE | 0.99637672 | 0.89844903 |
| DP13A_MOUSE | 0.97109727 | 0.40096222 |
| DPP10_MOUSE | 1.00518735 | 0.92117178 |
| DPP3_MOUSE | 0.82733183 | 0.01200881 |
| DPP6_MOUSE | 0.99761789 | 0.77206678 |
| DPYL1_MOUSE | 1.05114335 | 0.66023886 |
| DPYL2_MOUSE | 0.96468052 | 0.45238635 |
| DPYL3_MOUSE | 0.92718025 | 0.09345604 |
| DPYL4_MOUSE | 0.93956619 | 0.16580306 |
| DPYL5_MOUSE | 1.02932723 | 0.43527973 |
| DREB_MOUSE | 0.98338056 | 0.6081713 |
| DRG2_MOUSE | 1.05655813 | 0.7342031 |
| DRS7B_MOUSE | 0.91558019 | 0.05046646 |
| DTNA_MOUSE | 1.10998409 | 0.67179201 |
| DUS3_MOUSE | 0.94153834 | 0.2262102 |
| DYHC1_MOUSE | 0.99996041 | 0.84369198 |
| DYL2_MOUSE | 0.99610748 | 0.80791181 |
| DYN1_MOUSE | 0.97528094 | 0.4249306 |
| DYN2_MOUSE | 0.88939552 | 0.04723384 |
| DYN3_MOUSE | 1.0115484 | 0.9810353 |
| DYST_MOUSE | 1.00624913 | 0.65550488 |

| | | |
|---|---|---|
| E41L1_MOUSE | 0.99050655 | 0.63297553 |
| E41L2_MOUSE | 1.04101442 | 0.40409144 |
| E41L3_MOUSE | 1.04682247 | 0.27021034 |
| EAA1_MOUSE | 1.13709342 | 0.05145232 |
| EAA2_MOUSE | 1.11686724 | 0.16111993 |
| ECH1_MOUSE | 1.07770154 | 0.27514606 |
| ECHA_MOUSE | 1.0736688 | 0.11715792 |
| ECHB_MOUSE | 1.05260365 | 0.22163638 |
| ECHM_MOUSE | 0.99901761 | 0.78677796 |
| ECI1_MOUSE | 1.11886048 | 0.08396099 |
| ECI2_MOUSE | 1.0349254 | 0.93652359 |
| EEA1_MOUSE | 0.98202061 | 0.74602379 |
| EF1A1_MOUSE | 1.04052376 | 0.65534008 |
| EF1A2_MOUSE | 1.24083043 | 0.14898333 |
| EF1B_MOUSE | 0.92366787 | 0.15514706 |
| EF1D_MOUSE | 0.95404727 | 0.32038185 |
| EF1G_MOUSE | 1.01892642 | 0.997511 |
| EF2_MOUSE | 1.01448902 | 0.9026785 |
| EFGM_MOUSE | 0.93074539 | 0.14217418 |
| EFHD2_MOUSE | 0.95021554 | 0.16893521 |
| EFR3B_MOUSE | 1.03110436 | 0.28523835 |
| EFTS_MOUSE | 1.0346267 | 0.40961755 |
| EFTU_MOUSE | 0.99893827 | 0.79130208 |
| EHD1_MOUSE | 0.94490567 | 0.07504018 |
| EHD3_MOUSE | 0.97369621 | 0.35187298 |
| EHD4_MOUSE | 1.00516853 | 0.99944337 |
| EI3JA_MOUSE | 0.8001277 | 0.05996764 |
| EIF3A_MOUSE | 0.99040482 | 0.60801567 |
| ELMO2_MOUSE | 0.89811497 | 0.01129495 |
| ELOB_MOUSE | 1.14898073 | 0.17528143 |
| ELOC_MOUSE | 1.26510449 | 0.22061637 |
| EMC1_MOUSE | 1.13960032 | 0.12573863 |
| ENAH_MOUSE | 0.93990372 | 0.47147706 |
| ENDD1_MOUSE | 0.9772786 | 0.47484263 |
| ENOA_MOUSE | 1.00683064 | 0.84549813 |
| ENOG_MOUSE | 0.91257124 | 0.13439704 |
| ENPL_MOUSE | 1.0451462 | 0.32731152 |
| ENPP6_MOUSE | 0.77429732 | 0.00174224 |
| ENSA_MOUSE | 0.93155012 | 0.23966422 |

| | | |
|---|---|---|
| ENTP2_MOUSE | 1.23727555 | 0.10957402 |
| EP15R_MOUSE | 0.93191318 | 0.13536493 |
| EPHA4_MOUSE | 0.93087299 | 0.18969246 |
| EPMIP_MOUSE | 1.34136797 | 0.03202506 |
| EPN1_MOUSE | 0.89128225 | 0.02558887 |
| ERC2_MOUSE | 0.91653563 | 0.05824564 |
| ERF1_MOUSE | 0.89501746 | 0.01896492 |
| ERLN2_MOUSE | 1.09780132 | 0.04843022 |
| ERMIN_MOUSE | 0.70136735 | 0.0006461 |
| ERP29_MOUSE | 1.0100245 | 0.95273278 |
| ES1_MOUSE | 1.0409925 | 0.44203393 |
| ESTD_MOUSE | 1.03124716 | 0.62795788 |
| ETFA_MOUSE | 1.0460668 | 0.43475353 |
| ETFB_MOUSE | 1.03113387 | 0.38415183 |
| ETFD_MOUSE | 1.04376991 | 0.43807702 |
| ETHE1_MOUSE | 0.97350823 | 0.41798135 |
| EXC6B_MOUSE | 0.9326987 | 0.25123265 |
| EXOC1_MOUSE | 0.95258354 | 0.30967313 |
| EXOC2_MOUSE | 1.07266349 | 0.08495018 |
| EXOC3_MOUSE | 1.03025299 | 0.59908017 |
| EXOC4_MOUSE | 1.05356086 | 0.40041786 |
| EXOC5_MOUSE | 0.90683323 | 0.17298168 |
| EXOC7_MOUSE | 1.00110641 | 0.84174034 |
| EXOC8_MOUSE | 0.95971674 | 0.5372182 |
| EXOG_MOUSE | 1.10823534 | 0.2551025 |
| EZRI_MOUSE | 1.13008439 | 0.00765815 |
| F10A1_MOUSE | 0.95167764 | 0.21988638 |
| F1142_MOUSE | 0.92751538 | 0.19115024 |
| F126B_MOUSE | 1.02347194 | 0.68135928 |
| F136A_MOUSE | 1.13011404 | 0.25662989 |
| F162A_MOUSE | 1.11419842 | 0.37186432 |
| F1712_MOUSE | 0.89734581 | 0.1516368 |
| F213A_MOUSE | 1.04187096 | 0.36204647 |
| FA49B_MOUSE | 0.94825525 | 0.29022506 |
| FA81A_MOUSE | 0.89641254 | 0.13158071 |
| FAAA_MOUSE | 1.07930457 | 0.32457786 |
| FAAH1_MOUSE | 1.18543021 | 0.42186774 |
| FABP5_MOUSE | 1.06887186 | 0.32985143 |
| FABP7_MOUSE | 0.89541554 | 0.01497807 |

| | | |
|---|---|---|
| FABPH_MOUSE | 0.97068313 | 0.38553989 |
| FAD1_MOUSE | 0.95526062 | 0.55876374 |
| FAF2_MOUSE | 1.11270418 | 0.4015437 |
| FAHD1_MOUSE | 1.04220832 | 0.75349186 |
| FAHD2_MOUSE | 1.00052738 | 0.82713292 |
| FAK1_MOUSE | 0.90546026 | 0.03915314 |
| FAK2_MOUSE | 0.86129274 | 0.01378658 |
| FARP1_MOUSE | 1.08695481 | 0.37927058 |
| FAS_MOUSE | 1.00228816 | 0.9294778 |
| FBP1L_MOUSE | 1.25682396 | 0.39202978 |
| FBX2_MOUSE | 0.89586835 | 0.05059159 |
| FBX41_MOUSE | 1.084194 | 0.08931452 |
| FERM2_MOUSE | 1.01558709 | 0.7528284 |
| FGF1_MOUSE | 1.00311236 | 0.96833547 |
| FIS1_MOUSE | 1.05631133 | 0.96275341 |
| FKB1A_MOUSE | 0.99037756 | 0.64307739 |
| FKBP2_MOUSE | 1.09562595 | 0.07091616 |
| FKBP4_MOUSE | 1.06555138 | 0.442951 |
| FKBP8_MOUSE | 1.0166283 | 0.83882213 |
| FLOT1_MOUSE | 1.0500364 | 0.2668803 |
| FLOT2_MOUSE | 1.03616288 | 0.15662698 |
| FNTA_MOUSE | 1.25514043 | 0.10376756 |
| FPPS_MOUSE | 0.97889044 | 0.52279687 |
| FRIH_MOUSE | 0.98876314 | 0.65045371 |
| FRS1L_MOUSE | 0.94665898 | 0.29738819 |
| FSCN1_MOUSE | 1.00096874 | 0.85630928 |
| FSD1_MOUSE | 1.08443745 | 0.32867667 |
| FUMH_MOUSE | 1.02945542 | 0.69250377 |
| FUND2_MOUSE | 0.98019326 | 0.63108954 |
| FXL16_MOUSE | 0.95774399 | 0.36142269 |
| FXYD6_MOUSE | 1.00550852 | 0.56875657 |
| FYN_MOUSE | 1.13212404 | 0.02046221 |
| G3P_MOUSE | 1.00495118 | 0.91216678 |
| G6PD1_MOUSE | 0.99192471 | 0.8355243 |
| G6PI_MOUSE | 1.01287995 | 0.98114521 |
| GABR1_MOUSE | 1.00921034 | 0.87709605 |
| GABR2_MOUSE | 1.19471926 | 0.09213734 |
| GABT_MOUSE | 1.14023909 | 0.0939069 |
| GAK_MOUSE | 1.01875869 | 0.91231741 |

| | | |
|---|---|---|
| GANAB_MOUSE | 1.03330722 | 0.34374353 |
| GAS7_MOUSE | 0.94527382 | 0.27295367 |
| GBB1_MOUSE | 0.98883087 | 0.63395604 |
| GBB2_MOUSE | 0.98445213 | 0.63578959 |
| GBG12_MOUSE | 1.21638441 | 0.13046898 |
| GBRA1_MOUSE | 1.03930183 | 0.51960803 |
| GBRA3_MOUSE | 1.05650769 | 0.54664987 |
| GBRB2_MOUSE | 1.00909209 | 0.65021137 |
| GBRG2_MOUSE | 1.03554937 | 0.60848859 |
| GCDH_MOUSE | 1.04903765 | 0.15152767 |
| GCSP_MOUSE | 1.00781441 | 0.90413185 |
| GCYB1_MOUSE | 0.9935508 | 0.76844967 |
| GD1L1_MOUSE | 1.01278793 | 0.96060401 |
| GDAP1_MOUSE | 1.05457175 | 0.52691484 |
| GDE1_MOUSE | 1.06358373 | 0.86682562 |
| GDIA_MOUSE | 0.95526691 | 0.35910827 |
| GDIB_MOUSE | 0.97499466 | 0.50050969 |
| GDIR1_MOUSE | 0.9310952 | 0.10893807 |
| GDIR2_MOUSE | 1.0145506 | 0.98386481 |
| GDPD1_MOUSE | 0.99640758 | 0.6984245 |
| GELS_MOUSE | 0.89959665 | 0.03404801 |
| GEPH_MOUSE | 0.96420145 | 0.12394965 |
| GGT7_MOUSE | 0.96157393 | 0.14200352 |
| GHC1_MOUSE | 1.02600894 | 0.69470951 |
| GHC2_MOUSE | 1.13098581 | 0.0148315 |
| GHITM_MOUSE | 1.01704093 | 0.82645665 |
| GIT1_MOUSE | 0.97039765 | 0.47000531 |
| GLNA_MOUSE | 1.02215468 | 0.90575509 |
| GLO2_MOUSE | 1.04258018 | 0.74734337 |
| GLOD4_MOUSE | 1.00728895 | 0.91548343 |
| GLPK_MOUSE | 0.99256418 | 0.62122889 |
| GLRX3_MOUSE | 1.01231348 | 0.87357842 |
| GLRX5_MOUSE | 1.00267135 | 0.69663536 |
| GLSK_MOUSE | 0.9905067 | 0.6601556 |
| GLTP_MOUSE | 0.80062921 | 0.00065311 |
| GLU2B_MOUSE | 0.95082672 | 0.58245884 |
| GMFB_MOUSE | 1.01674218 | 0.47509811 |
| GNA11_MOUSE | 0.96719908 | 0.35009611 |
| GNA13_MOUSE | 1.02471894 | 0.7575927 |

| | | |
|---|---|---|
| GNAI1_MOUSE | 0.93551604 | 0.20103305 |
| GNAI2_MOUSE | 0.99326702 | 0.64699244 |
| GNAL_MOUSE | 0.91293456 | 0.23308404 |
| GNAO_MOUSE | 1.0334088 | 0.73332862 |
| GNAQ_MOUSE | 1.01520292 | 0.9240941 |
| GNAS1_MOUSE | 0.99259823 | 0.67681486 |
| GNAZ_MOUSE | 0.98146205 | 0.56178768 |
| GNL1_MOUSE | 1.01604335 | 0.9174514 |
| GP158_MOUSE | 0.97225362 | 0.33973596 |
| GPD1L_MOUSE | 0.98262733 | 0.4886019 |
| GPDA_MOUSE | 0.95424183 | 0.2820429 |
| GPDM_MOUSE | 1.02325067 | 0.59866467 |
| GPM6A_MOUSE | 1.04278309 | 0.65782388 |
| GPM6B_MOUSE | 0.93013583 | 0.20792359 |
| GPX1_MOUSE | 1.14533197 | 0.42531989 |
| GPX41_MOUSE | 1.00030779 | 0.87783759 |
| GRAP1_MOUSE | 0.90511427 | 0.2555977 |
| GRB2_MOUSE | 0.96697427 | 0.36363443 |
| GRHPR_MOUSE | 1.00215138 | 0.89001022 |
| GRIA1_MOUSE | 0.9917401 | 0.63537587 |
| GRIA2_MOUSE | 0.99801825 | 0.80235 |
| GRIA3_MOUSE | 1.03254364 | 0.47479562 |
| GRIA4_MOUSE | 0.91835384 | 0.15364045 |
| GRIK2_MOUSE | 1.01228953 | 0.99953405 |
| GRIN1_MOUSE | 0.96095874 | 0.40598644 |
| GRM2_MOUSE | 0.98199966 | 0.53965784 |
| GRM3_MOUSE | 1.01481414 | 0.95494592 |
| GRM5_MOUSE | 0.98004857 | 0.73559814 |
| GRP75_MOUSE | 0.98695 | 0.57954 |
| GRP78_MOUSE | 0.99697679 | 0.75812564 |
| GRPE1_MOUSE | 0.96205474 | 0.41323019 |
| GSH1_MOUSE | 0.96178101 | 0.43616771 |
| GSHB_MOUSE | 0.85033172 | 0.02268042 |
| GSHR_MOUSE | 1.02567347 | 0.7619371 |
| GSK3A_MOUSE | 0.96525689 | 0.35794847 |
| GSK3B_MOUSE | 0.98757273 | 0.52690272 |
| GSLG1_MOUSE | 0.95830048 | 0.73752093 |
| GSTA4_MOUSE | 1.02662765 | 0.75157663 |
| GSTK1_MOUSE | 1.16557025 | 0.01233682 |

| | | |
|---|---|---|
| GSTM1_MOUSE | 1.15518006 | 0.08652935 |
| GSTM5_MOUSE | 0.95707989 | 0.08519237 |
| GSTM7_MOUSE | 1.03962189 | 0.97102673 |
| GSTO1_MOUSE | 0.97255265 | 0.40249991 |
| GSTP1_MOUSE | 1.01889342 | 0.97019997 |
| GTR1_MOUSE | 1.03829364 | 0.70525455 |
| GTR3_MOUSE | 0.97449832 | 0.4580693 |
| GUAA_MOUSE | 0.98635927 | 0.54324771 |
| GUAD_MOUSE | 0.9358393 | 0.0666416 |
| H2B1B_MOUSE | 1.80862575 | 0.09714643 |
| H4_MOUSE | 1.41146582 | 0.21945612 |
| HACD3_MOUSE | 1.09368568 | 0.30131823 |
| HBA_MOUSE | 0.9966499 | 0.74706481 |
| HBB1_MOUSE | 0.96325363 | 0.57335766 |
| HCD2_MOUSE | 1.0063288 | 0.93645434 |
| HCDH_MOUSE | 1.12175114 | 0.31525542 |
| HCN1_MOUSE | 0.94301013 | 0.48815647 |
| HCN2_MOUSE | 1.15046562 | 0.18074168 |
| HD_MOUSE | 0.91220661 | 0.12669536 |
| HDHD2_MOUSE | 1.16675473 | 0.14740017 |
| HDHD3_MOUSE | 1.19753055 | 0.08072622 |
| HEBP1_MOUSE | 0.82502717 | 0.01120527 |
| HECAM_MOUSE | 1.00378752 | 0.6768445 |
| HEM2_MOUSE | 1.04526424 | 0.97466326 |
| HEM6_MOUSE | 0.87976637 | 0.01536622 |
| HEMH_MOUSE | 1.06645626 | 0.30082923 |
| HEXB_MOUSE | 1.25357201 | 7.8042E-05 |
| HGS_MOUSE | 0.94769661 | 0.40013508 |
| HIBCH_MOUSE | 1.0059509 | 0.90455214 |
| HINT1_MOUSE | 1.02699273 | 0.93559054 |
| HINT2_MOUSE | 1.13807736 | 0.17038312 |
| HIP1R_MOUSE | 1.07038403 | 0.03480123 |
| HMGCL_MOUSE | 1.0969999 | 0.18029747 |
| HMOX2_MOUSE | 1.01384104 | 0.98304365 |
| HNRPD_MOUSE | 1.00846097 | 0.84607034 |
| HNRPK_MOUSE | 0.91002648 | 0.0314375 |
| HNRPQ_MOUSE | 0.97713969 | 0.35580287 |
| HOME1_MOUSE | 0.92744494 | 0.02917598 |
| HOME2_MOUSE | 0.96944594 | 0.46724973 |

| | | |
|---|---|---|
| HPCA_MOUSE | 0.88978881 | 0.13213266 |
| HPCL1_MOUSE | 0.81808759 | 0.01486627 |
| HPCL4_MOUSE | 1.00947008 | 0.9821272 |
| HPLN1_MOUSE | 1.01779418 | 0.74266599 |
| HPRT_MOUSE | 0.98731662 | 0.64991697 |
| HS105_MOUSE | 1.01309323 | 0.77386254 |
| HS12A_MOUSE | 1.00111196 | 0.80187899 |
| HS74L_MOUSE | 0.97532925 | 0.52169026 |
| HS90A_MOUSE | 1.0514396 | 0.77030603 |
| HS90B_MOUSE | 1.05436386 | 0.70537067 |
| HSDL1_MOUSE | 1.16066317 | 0.21571423 |
| HSDL2_MOUSE | 1.19055672 | 0.2311899 |
| HSP72_MOUSE | 0.89959424 | 0.02410683 |
| HSP74_MOUSE | 0.99306137 | 0.72726329 |
| HSP7C_MOUSE | 0.9805482 | 0.4997829 |
| HUWE1_MOUSE | 1.35876657 | 0.13870994 |
| HXK1_MOUSE | 1.04056069 | 0.48317218 |
| HYEP_MOUSE | 1.35368608 | 0.26282332 |
| HYES_MOUSE | 1.1278738 | 0.11275112 |
| HYOU1_MOUSE | 1.04149257 | 0.33220653 |
| ICAM5_MOUSE | 1.15414603 | 0.18747463 |
| IDH3A_MOUSE | 1.03306397 | 0.79348352 |
| IDHC_MOUSE | 0.98353491 | 0.532206 |
| IDHG1_MOUSE | 1.05339529 | 0.57543681 |
| IDHP_MOUSE | 1.08284323 | 0.10659142 |
| IDI1_MOUSE | 0.95106736 | 0.36793017 |
| IF2A_MOUSE | 0.97077302 | 0.43973674 |
| IF2M_MOUSE | 0.95094877 | 0.24604319 |
| IF4A1_MOUSE | 0.97879608 | 0.71371297 |
| IF4A2_MOUSE | 0.99653524 | 0.58493436 |
| IF4B_MOUSE | 0.84114672 | 0.08381776 |
| IF4G2_MOUSE | 0.90824229 | 0.1655952 |
| IF4G3_MOUSE | 2.13082595 | 0.2253589 |
| IF4H_MOUSE | 0.906618 | 0.01605254 |
| IF5A1_MOUSE | 1.07810511 | 0.46918629 |
| IGS21_MOUSE | 1.02254918 | 0.77797533 |
| IGSF8_MOUSE | 0.90445383 | 0.09063181 |
| ILDR2_MOUSE | 1.16725867 | 0.0204878 |
| ILEUA_MOUSE | 0.90670093 | 0.15485945 |

| | | |
|---|---|---|
| IMA4_MOUSE | 1.07562575 | 0.34381595 |
| IMB1_MOUSE | 0.96736532 | 0.22525699 |
| IMPA1_MOUSE | 0.94045773 | 0.06342261 |
| INP4A_MOUSE | 1.19387803 | 0.15036698 |
| INPP_MOUSE | 0.93495993 | 0.24798857 |
| IP3KA_MOUSE | 1.03272533 | 0.29738274 |
| IPO5_MOUSE | 1.03547547 | 0.50682953 |
| IPO7_MOUSE | 0.96886146 | 0.46091063 |
| IPO9_MOUSE | 1.12071018 | 0.62653297 |
| IPYR_MOUSE | 0.90182479 | 0.11972807 |
| IPYR2_MOUSE | 1.03919807 | 0.70320786 |
| IQEC1_MOUSE | 1.01964223 | 0.84060832 |
| IQEC2_MOUSE | 0.97027953 | 0.23230766 |
| IQEC3_MOUSE | 0.96746136 | 0.3646891 |
| ISC2A_MOUSE | 1.45600583 | 0.07998865 |
| ISCA1_MOUSE | 0.96274718 | 0.38011298 |
| ISCA2_MOUSE | 0.97379008 | 0.39297602 |
| ISCU_MOUSE | 1.00502571 | 0.92568684 |
| ITAV_MOUSE | 1.07288518 | 0.49259874 |
| ITPA_MOUSE | 1.01960454 | 0.99858423 |
| ITPR1_MOUSE | 0.94623038 | 0.40546749 |
| ITSN1_MOUSE | 0.90674873 | 0.13140552 |
| IVD_MOUSE | 1.01863486 | 0.58514458 |
| JAM3_MOUSE | 0.82781692 | 0.00443631 |
| JIP3_MOUSE | 0.98153519 | 0.47927761 |
| K0513_MOUSE | 0.92586537 | 0.05359384 |
| K1107_MOUSE | 0.98621988 | 0.41971006 |
| K1468_MOUSE | 0.94151881 | 0.21647293 |
| KAD1_MOUSE | 0.90544477 | 0.05105984 |
| KAD3_MOUSE | 1.07099764 | 0.07619417 |
| KAD4_MOUSE | 1.09068021 | 0.29152758 |
| KAD5_MOUSE | 0.88686762 | 0.21673854 |
| KALRN_MOUSE | 0.97906843 | 0.48625297 |
| KAP0_MOUSE | 1.00790209 | 0.86173492 |
| KAP2_MOUSE | 0.9940229 | 0.69684 |
| KAP3_MOUSE | 1.05555251 | 0.30122207 |
| KAPCA_MOUSE | 0.98782347 | 0.676332 |
| KAPCB_MOUSE | 1.13611473 | 0.30706864 |
| KAT3_MOUSE | 1.06803927 | 0.62888836 |

| | | |
|---|---|---|
| KBTBB_MOUSE | 1.01297014 | 0.91865502 |
| KCAB2_MOUSE | 1.14419735 | 0.01641592 |
| KCC1D_MOUSE | 0.92286057 | 0.20861129 |
| KCC2A_MOUSE | 0.94621948 | 0.2878846 |
| KCC2B_MOUSE | 0.93683967 | 0.23063677 |
| KCC2D_MOUSE | 0.99126464 | 0.7451315 |
| KCC2G_MOUSE | 0.88936051 | 0.06989424 |
| KCC4_MOUSE | 0.86225744 | 0.00240262 |
| KCD12_MOUSE | 0.89011137 | 0.09686777 |
| KCD16_MOUSE | 1.1265359 | 0.19257008 |
| KCJ10_MOUSE | 1.18909646 | 0.01845462 |
| KCMA1_MOUSE | 1.07136577 | 0.09498372 |
| KCNA1_MOUSE | 0.95670716 | 0.3306516 |
| KCNA6_MOUSE | 1.00342447 | 0.82905461 |
| KCNC3_MOUSE | 0.99013502 | 0.76865145 |
| KCND2_MOUSE | 0.96983282 | 0.59478545 |
| KCRB_MOUSE | 0.95922771 | 0.24147638 |
| KCRU_MOUSE | 0.98397 | 0.59662836 |
| KCY_MOUSE | 0.96140981 | 0.20306035 |
| KI21A_MOUSE | 1.01049604 | 0.94805992 |
| KIF1A_MOUSE | 0.94575693 | 0.14319105 |
| KIF2A_MOUSE | 0.99218939 | 0.70145481 |
| KIF5C_MOUSE | 0.99174493 | 0.63416541 |
| KINH_MOUSE | 0.98484243 | 0.59842804 |
| KKCC2_MOUSE | 0.89425397 | 0.05757814 |
| KLC1_MOUSE | 0.922739 | 0.14459159 |
| KLC2_MOUSE | 1.03582973 | 0.76964758 |
| KPCA_MOUSE | 1.05799711 | 0.63840216 |
| KPCB_MOUSE | 0.99422957 | 0.68465005 |
| KPCE_MOUSE | 0.92950432 | 0.07828037 |
| KPCG_MOUSE | 0.94592617 | 0.25412851 |
| KPRB_MOUSE | 0.97140031 | 0.56822374 |
| KPYM_MOUSE | 0.9950205 | 0.74245051 |
| KTN1_MOUSE | 0.93140812 | 0.49453738 |
| L1CAM_MOUSE | 0.95412136 | 0.14742163 |
| L2GL1_MOUSE | 0.93869431 | 0.10888802 |
| L2HDH_MOUSE | 1.02902983 | 0.77627952 |
| LACTB_MOUSE | 0.99047184 | 0.75012811 |
| LAMP1_MOUSE | 1.40152611 | 0.00716468 |

| | | |
|---|---|---|
| LAMP2_MOUSE | 1.26534597 | 0.03109832 |
| LANC1_MOUSE | 1.00586194 | 0.85129628 |
| LANC2_MOUSE | 0.96555558 | 0.37886485 |
| LASP1_MOUSE | 1.07232237 | 0.45671315 |
| LAT1_MOUSE | 1.09288664 | 0.32592752 |
| LDHA_MOUSE | 1.0037108 | 0.8738417 |
| LDHB_MOUSE | 1.00644869 | 0.93920366 |
| LEGL_MOUSE | 1.02286825 | 0.58196224 |
| LETM1_MOUSE | 0.96141347 | 0.28213711 |
| LGI1_MOUSE | 1.01443788 | 0.88293982 |
| LGI3_MOUSE | 0.98087268 | 0.64846702 |
| LGUL_MOUSE | 0.88465545 | 0.02776196 |
| LIGO1_MOUSE | 0.92535483 | 0.0662017 |
| LIN7A_MOUSE | 1.04419579 | 0.60689541 |
| LIN7B_MOUSE | 1.01568644 | 0.97559783 |
| LIN7C_MOUSE | 0.95853856 | 0.35366281 |
| LIPA2_MOUSE | 1.02838839 | 0.9503486 |
| LIPA3_MOUSE | 0.9300342 | 0.18815451 |
| LIS1_MOUSE | 0.9655425 | 0.35960492 |
| LKHA4_MOUSE | 1.03751804 | 0.73717574 |
| LNEBL_MOUSE | 0.98516024 | 0.66806 |
| LNP_MOUSE | 0.91369094 | 0.1144901 |
| LONM_MOUSE | 1.00933699 | 0.97320616 |
| LPPRC_MOUSE | 1.06369256 | 0.12686758 |
| LRC47_MOUSE | 0.93331977 | 0.21386983 |
| LRC4B_MOUSE | 0.94974307 | 0.25142476 |
| LRC57_MOUSE | 1.00165862 | 0.76285802 |
| LRC59_MOUSE | 0.92082264 | 0.27640336 |
| LRC8A_MOUSE | 1.04081756 | 0.74252674 |
| LRP1_MOUSE | 1.05107247 | 0.26354775 |
| LRRC7_MOUSE | 0.94852172 | 0.21733151 |
| LRRT4_MOUSE | 0.93034258 | 0.12192532 |
| LSAMP_MOUSE | 1.0151648 | 0.95933566 |
| LXN_MOUSE | 0.98544172 | 0.61593876 |
| LY6H_MOUSE | 0.73551181 | 0.0012169 |
| LYAG_MOUSE | 0.96022741 | 0.4989338 |
| LYPA2_MOUSE | 1.62648895 | 0.16854832 |
| M2OM_MOUSE | 1.02219577 | 0.84290835 |
| MA2C1_MOUSE | 1.06035861 | 0.96505014 |

| | | |
|---|---|---|
| MA7D2_MOUSE | 0.80489912 | 0.00460827 |
| MAAI_MOUSE | 1.15831227 | 0.08578306 |
| MACF1_MOUSE | 0.94708023 | 0.32818398 |
| MADD_MOUSE | 0.98573014 | 0.70255617 |
| MAG_MOUSE | 0.86701761 | 0.03794073 |
| MAGI2_MOUSE | 0.98109207 | 0.53730691 |
| MAOM_MOUSE | 1.01600988 | 0.86977351 |
| MAON_MOUSE | 1.00489047 | 0.84958449 |
| MAOX_MOUSE | 1.06690901 | 0.12073121 |
| MAP1A_MOUSE | 0.92195386 | 0.00325784 |
| MAP1B_MOUSE | 0.9432258 | 0.12478056 |
| MAP1S_MOUSE | 0.95264332 | 0.17528012 |
| MAP4_MOUSE | 0.89066556 | 0.01688194 |
| MAP6_MOUSE | 0.92475347 | 0.09264398 |
| MARC2_MOUSE | 1.11923826 | 0.03377912 |
| MARCS_MOUSE | 0.97101937 | 0.58943751 |
| MARE1_MOUSE | 1.04629156 | 0.30955014 |
| MARE2_MOUSE | 0.9926666 | 0.77606803 |
| MARE3_MOUSE | 0.98540773 | 0.58928631 |
| MARK2_MOUSE | 1.02383251 | 0.85802327 |
| MAT2B_MOUSE | 0.93985703 | 0.23022174 |
| MBLC2_MOUSE | 0.90343498 | 0.07978019 |
| MBP_MOUSE | 0.92328741 | 0.4454055 |
| MCAT_MOUSE | 1.04065071 | 0.47650239 |
| MCCA_MOUSE | 1.02846836 | 0.77068371 |
| MCCB_MOUSE | 1.07818015 | 0.362468 |
| MCTS1_MOUSE | 1.2107085 | 0.50599744 |
| MCU_MOUSE | 1.0260794 | 0.66581334 |
| MDHC_MOUSE | 0.97076836 | 0.37386438 |
| MDHM_MOUSE | 1.00730397 | 0.96203795 |
| MECR_MOUSE | 1.27092876 | 0.0299552 |
| METK2_MOUSE | 1.05221617 | 0.98342534 |
| MFF_MOUSE | 1.1429367 | 0.08036632 |
| MFN2_MOUSE | 1.01648063 | 0.95663788 |
| MFR1L_MOUSE | 0.94501509 | 0.19059654 |
| MGLL_MOUSE | 1.08016729 | 0.50201274 |
| MGST3_MOUSE | 1.1904103 | 0.15188327 |
| MIA40_MOUSE | 0.99153095 | 0.74890825 |
| MIC13_MOUSE | 1.11846878 | 0.18306348 |

| | | |
|---|---|---|
| MIC19_MOUSE | 0.99062762 | 0.62120066 |
| MIC25_MOUSE | 1.04625483 | 0.74891666 |
| MIC26_MOUSE | 1.02690877 | 0.89991855 |
| MIC27_MOUSE | 1.0358291 | 0.2455327 |
| MIC60_MOUSE | 0.96442481 | 0.33929642 |
| MICA3_MOUSE | 0.88265171 | 0.04276538 |
| MICU1_MOUSE | 0.96058267 | 0.37202076 |
| MICU3_MOUSE | 1.03362737 | 0.7407846 |
| MIF_MOUSE | 1.01136338 | 0.8920451 |
| MINK1_MOUSE | 1.05338813 | 0.26692958 |
| MIRO1_MOUSE | 0.94261506 | 0.1258291 |
| MIRO2_MOUSE | 1.12920101 | 0.06287708 |
| MK01_MOUSE | 1.01717077 | 0.83954249 |
| MK03_MOUSE | 0.99686119 | 0.88195385 |
| ML12B_MOUSE | 0.90102778 | 0.07550757 |
| MLC1_MOUSE | 1.08359027 | 0.75267065 |
| MLEC_MOUSE | 1.00699818 | 0.8871345 |
| MMSA_MOUSE | 1.07307357 | 0.14809918 |
| MOES_MOUSE | 1.0951602 | 0.4149014 |
| MOG_MOUSE | 0.81062557 | 0.04203528 |
| MOT1_MOUSE | 1.11464269 | 0.75318346 |
| MP2K1_MOUSE | 0.98046129 | 0.58968413 |
| MP2K2_MOUSE | 0.9120673 | 0.03546896 |
| MP2K4_MOUSE | 1.02065888 | 0.92241121 |
| MPC2_MOUSE | 1.32217013 | 0.00070535 |
| MPCP_MOUSE | 1.06910064 | 0.12559273 |
| MPI_MOUSE | 0.97964985 | 0.65587716 |
| MPP2_MOUSE | 1.11963475 | 0.04303153 |
| MPP3_MOUSE | 0.94265337 | 0.10859245 |
| MPP6_MOUSE | 1.05433544 | 0.54897412 |
| MPPA_MOUSE | 1.02924742 | 0.74045795 |
| MPPB_MOUSE | 1.12859517 | 0.74639868 |
| MRCKB_MOUSE | 0.93370468 | 0.03820849 |
| MSRA_MOUSE | 1.56066202 | 0.24394602 |
| MTAP2_MOUSE | 0.96096004 | 0.29523747 |
| MTCH1_MOUSE | 0.97208549 | 0.42474593 |
| MTCH2_MOUSE | 1.02016669 | 0.75168631 |
| MTEF2_MOUSE | 1.17645415 | 0.24683986 |
| MTMR1_MOUSE | 0.93901725 | 0.14697626 |

| | | |
|---|---|---|
| MTMR2_MOUSE | 0.90768323 | 0.16139349 |
| MTMR5_MOUSE | 0.97853735 | 0.46341999 |
| MTOR_MOUSE | 1.03595867 | 0.8085487 |
| MTPN_MOUSE | 0.95693534 | 0.2902891 |
| MTX1_MOUSE | 1.00711524 | 0.79443812 |
| MTX2_MOUSE | 1.03237818 | 0.64176554 |
| MUTA_MOUSE | 1.00531712 | 0.95113874 |
| MY18A_MOUSE | 1.02635245 | 0.86903555 |
| MYCT_MOUSE | 0.95986112 | 0.49220702 |
| MYH10_MOUSE | 0.9592996 | 0.36358753 |
| MYH14_MOUSE | 0.9453957 | 0.30542759 |
| MYH9_MOUSE | 0.92656046 | 0.30806453 |
| MYL6_MOUSE | 0.90022221 | 0.04429835 |
| MYO1D_MOUSE | 0.87790012 | 0.13191371 |
| MYO5A_MOUSE | 0.99772778 | 0.83136478 |
| MYO6_MOUSE | 0.92565205 | 0.16830803 |
| MYPR_MOUSE | 0.89549461 | 0.05094985 |
| NAC1_MOUSE | 1.06939221 | 0.26206924 |
| NAC2_MOUSE | 1.00869748 | 0.96067254 |
| NACA_MOUSE | 0.86900417 | 0.02119228 |
| NAGAB_MOUSE | 0.85394623 | 0.02971335 |
| NAGK_MOUSE | 1.10209212 | 0.68275629 |
| NAKD2_MOUSE | 1.03332278 | 0.30706702 |
| NAMPT_MOUSE | 1.11718972 | 0.43285243 |
| NB5R1_MOUSE | 1.09370359 | 0.24967383 |
| NB5R3_MOUSE | 1.0399061 | 0.25344097 |
| NBEA_MOUSE | 1.02745106 | 0.73967195 |
| NCALD_MOUSE | 0.8888428 | 0.05941141 |
| NCAM1_MOUSE | 1.00922019 | 0.93701912 |
| NCAM2_MOUSE | 1.06230671 | 0.06651004 |
| NCAN_MOUSE | 0.92976484 | 0.26067608 |
| NCDN_MOUSE | 0.97663871 | 0.29288803 |
| NCEH1_MOUSE | 1.01867947 | 0.63870012 |
| NCKP1_MOUSE | 1.02066555 | 0.68898707 |
| NCPR_MOUSE | 1.04230958 | 0.55276667 |
| NCS1_MOUSE | 1.02989954 | 0.90692259 |
| NDKA_MOUSE | 1.058308 | 0.16941974 |
| NDKB_MOUSE | 1.12502218 | 0.40143463 |
| NDRG1_MOUSE | 0.94785859 | 0.442828 |

| | | |
|---|---|---|
| NDRG2_MOUSE | 0.99018196 | 0.64431898 |
| NDRG3_MOUSE | 0.89548669 | 0.08100714 |
| NDRG4_MOUSE | 0.95069216 | 0.27890409 |
| NDUA1_MOUSE | 1.0132651 | 0.88219742 |
| NDUA2_MOUSE | 1.03051931 | 0.95158121 |
| NDUA4_MOUSE | 1.14503956 | 0.00412785 |
| NDUA5_MOUSE | 1.12733723 | 0.27634888 |
| NDUA6_MOUSE | 1.08844412 | 0.17983409 |
| NDUA7_MOUSE | 0.98251045 | 0.64620272 |
| NDUA8_MOUSE | 1.07774647 | 0.28945013 |
| NDUA9_MOUSE | 1.15779457 | 0.16088299 |
| NDUAA_MOUSE | 1.06895499 | 0.30846433 |
| NDUAB_MOUSE | 1.12938014 | 0.12287275 |
| NDUAC_MOUSE | 1.0793315 | 0.25434492 |
| NDUAD_MOUSE | 1.0688768 | 0.20339444 |
| NDUB1_MOUSE | 1.12926102 | 0.13593936 |
| NDUB3_MOUSE | 1.08217393 | 0.29072781 |
| NDUB4_MOUSE | 1.07374921 | 0.379376 |
| NDUB5_MOUSE | 0.97754016 | 0.48708157 |
| NDUB6_MOUSE | 1.06170335 | 0.45320348 |
| NDUB7_MOUSE | 1.10087822 | 0.22268906 |
| NDUB8_MOUSE | 0.91689878 | 0.03792883 |
| NDUB9_MOUSE | 1.31423046 | 0.17222648 |
| NDUBA_MOUSE | 0.98479154 | 0.55529063 |
| NDUBB_MOUSE | 1.00731954 | 0.95369061 |
| NDUC2_MOUSE | 1.06758252 | 0.44422729 |
| NDUF3_MOUSE | 1.07974234 | 0.42007626 |
| NDUS1_MOUSE | 0.96650326 | 0.28283496 |
| NDUS2_MOUSE | 1.0630756 | 0.73636905 |
| NDUS3_MOUSE | 1.02627792 | 0.83733642 |
| NDUS4_MOUSE | 1.00218457 | 0.89041652 |
| NDUS5_MOUSE | 1.01526487 | 0.88259846 |
| NDUS6_MOUSE | 1.06807777 | 0.51588702 |
| NDUS7_MOUSE | 1.09640032 | 0.3709634 |
| NDUS8_MOUSE | 0.99676192 | 0.76320379 |
| NDUV1_MOUSE | 1.07947914 | 0.33487841 |
| NDUV2_MOUSE | 0.99474737 | 0.68649134 |
| NEB2_MOUSE | 0.93550828 | 0.21287939 |
| NECA2_MOUSE | 0.87940408 | 0.03012099 |

| | | |
|---|---|---|
| NECP1_MOUSE | 0.95406575 | 0.21445126 |
| NEDD4_MOUSE | 0.84642226 | 0.12338064 |
| NEGR1_MOUSE | 0.94515288 | 0.1781445 |
| NEO1_MOUSE | 0.97520147 | 0.69255203 |
| NEUL_MOUSE | 0.91627141 | 0.05404982 |
| NEUM_MOUSE | 0.95603322 | 0.32362414 |
| NF1_MOUSE | 0.95071121 | 0.10359784 |
| NFASC_MOUSE | 1.02754702 | 0.74451862 |
| NFL_MOUSE | 1.20207472 | 0.0390092 |
| NFM_MOUSE | 1.20289993 | 0.05015881 |
| NFS1_MOUSE | 1.05719788 | 0.62294388 |
| NFU1_MOUSE | 0.89839858 | 0.06465322 |
| NGEF_MOUSE | 1.02519579 | 0.4896533 |
| NHRF1_MOUSE | 1.04269501 | 0.72023942 |
| NICA_MOUSE | 1.04294497 | 0.39422951 |
| NIF3L_MOUSE | 1.09041936 | 0.18156027 |
| NIPS1_MOUSE | 1.06533237 | 0.51417049 |
| NIPS2_MOUSE | 1.04139721 | 0.69284516 |
| NIT1_MOUSE | 1.10078875 | 0.23534919 |
| NIT2_MOUSE | 1.13577144 | 0.16474237 |
| NLGN3_MOUSE | 1.05900744 | 0.17582176 |
| NLTP_MOUSE | 1.06173948 | 0.1235345 |
| NMDE1_MOUSE | 0.97037595 | 0.38307683 |
| NMDE2_MOUSE | 0.95183862 | 0.33109324 |
| NMDZ1_MOUSE | 0.95947969 | 0.27012383 |
| NMT2_MOUSE | 1.06267801 | 0.901335 |
| NNRD_MOUSE | 0.98494453 | 0.5806195 |
| NNRE_MOUSE | 1.00790381 | 0.88367596 |
| NNTM_MOUSE | 6.67564333 | 0.05216051 |
| NOE1_MOUSE | 0.92599746 | 0.06978157 |
| NOMO1_MOUSE | 0.97066183 | 0.46179249 |
| NP1L1_MOUSE | 0.88747556 | 0.02188194 |
| NP1L4_MOUSE | 0.85882754 | 0.01284868 |
| NPL4_MOUSE | 0.97171711 | 0.62481358 |
| NPS3B_MOUSE | 1.13234199 | 0.11440893 |
| NPTN_MOUSE | 1.03143162 | 0.66149391 |
| NPTX1_MOUSE | 0.90833515 | 0.01447777 |
| NPTXR_MOUSE | 0.88738283 | 0.07150321 |
| NRCAM_MOUSE | 1.14839269 | 0.31441831 |

| | | |
|---|---|---|
| NRX1A_MOUSE | 1.01682435 | 0.94817938 |
| NRX3A_MOUSE | 1.00566944 | 0.95306512 |
| NSF_MOUSE | 0.99322478 | 0.65001745 |
| NSF1C_MOUSE | 0.95379788 | 0.26623312 |
| NT5D3_MOUSE | 1.05916555 | 0.21744058 |
| NTF2_MOUSE | 1.1785295 | 0.05905976 |
| NTRI_MOUSE | 0.98825748 | 0.57368287 |
| NTRK2_MOUSE | 1.09437487 | 0.3594508 |
| NU1M_MOUSE | 1.15263395 | 0.39224592 |
| NU5M_MOUSE | 1.16804406 | 0.24218301 |
| NUCG_MOUSE | 0.98424953 | 0.65962729 |
| NUDC_MOUSE | 1.2086636 | 0.00217095 |
| NUDT3_MOUSE | 0.92530622 | 0.06406152 |
| OAT_MOUSE | 1.12313648 | 0.24280878 |
| OCAD1_MOUSE | 0.92184018 | 0.08040909 |
| ODB2_MOUSE | 1.1042464 | 0.07019949 |
| ODBA_MOUSE | 0.99953326 | 0.83828165 |
| ODBB_MOUSE | 1.00248265 | 0.63956314 |
| ODO1_MOUSE | 1.02134296 | 0.86854725 |
| ODO2_MOUSE | 0.94457978 | 0.09527402 |
| ODP2_MOUSE | 0.96444011 | 0.20698769 |
| ODPA_MOUSE | 1.0249564 | 0.66129171 |
| ODPB_MOUSE | 0.99904408 | 0.76815612 |
| ODPX_MOUSE | 0.95428042 | 0.03605176 |
| OGA_MOUSE | 1.05758816 | 0.37620006 |
| OGT1_MOUSE | 0.99876822 | 0.84780331 |
| OLA1_MOUSE | 1.10552791 | 0.2503684 |
| OMGP_MOUSE | 0.94813176 | 0.40692267 |
| OMP_MOUSE | 0.75077186 | 0.07260789 |
| OPA1_MOUSE | 0.97592061 | 0.32270261 |
| OSB10_MOUSE | 0.98453597 | 0.68213975 |
| OSBL1_MOUSE | 1.02333705 | 0.89751222 |
| OSBL8_MOUSE | 0.99106234 | 0.65315863 |
| OSCP1_MOUSE | 1.05345987 | 0.61866299 |
| OST48_MOUSE | 1.01504547 | 0.97785333 |
| OTUB1_MOUSE | 0.95155946 | 0.36339172 |
| OX2G_MOUSE | 1.04870317 | 0.55787902 |
| OXR1_MOUSE | 0.96822046 | 0.37130483 |
| OXSR1_MOUSE | 0.90056682 | 0.10668183 |

| | | |
|---|---|---|
| P5CR2_MOUSE | 1.02113577 | 0.86756015 |
| P5CS_MOUSE | 0.95373377 | 0.2186305 |
| PA1B2_MOUSE | 1.02100765 | 0.79967824 |
| PA2G4_MOUSE | 0.94860807 | 0.05160568 |
| PABP1_MOUSE | 0.95953523 | 0.19427743 |
| PACN1_MOUSE | 0.85681121 | 0.03486825 |
| PACN2_MOUSE | 0.9578468 | 0.46600372 |
| PACS1_MOUSE | 0.97262035 | 0.30162945 |
| PACS2_MOUSE | 1.0461893 | 0.91913398 |
| PADI2_MOUSE | 0.9481439 | 0.45997087 |
| PAK1_MOUSE | 0.94763079 | 0.25417587 |
| PALM_MOUSE | 0.93079139 | 0.2173068 |
| PALM2_MOUSE | 1.10413941 | 0.88475218 |
| PARK7_MOUSE | 0.95404056 | 0.18930165 |
| PCBP1_MOUSE | 1.00713177 | 0.95802775 |
| PCBP2_MOUSE | 0.93977297 | 0.08755091 |
| PCCA_MOUSE | 0.95424421 | 0.20346456 |
| PCCB_MOUSE | 1.00090596 | 0.81193898 |
| PCKGM_MOUSE | 0.95715302 | 0.25383051 |
| PCLO_MOUSE | 1.01693286 | 0.94870891 |
| PCSK1_MOUSE | 0.91762521 | 0.21548108 |
| PCY2_MOUSE | 0.96174223 | 0.37457413 |
| PCYOX_MOUSE | 1.15507254 | 0.17640501 |
| PDC6I_MOUSE | 0.95421676 | 0.24845789 |
| PDCD5_MOUSE | 0.76320979 | 0.00275629 |
| PDCD6_MOUSE | 0.97101167 | 0.36312258 |
| PDE10_MOUSE | 1.11043813 | 0.08992869 |
| PDE1A_MOUSE | 0.89786772 | 0.04629677 |
| PDE1B_MOUSE | 1.00526591 | 0.93548675 |
| PDE2A_MOUSE | 1.04812793 | 0.33920883 |
| PDIA1_MOUSE | 1.00575048 | 0.94270369 |
| PDIA3_MOUSE | 0.9756158 | 0.57172246 |
| PDIA4_MOUSE | 0.95474027 | 0.28739388 |
| PDIA6_MOUSE | 0.99920898 | 0.83359917 |
| PDIP2_MOUSE | 0.9673986 | 0.44706677 |
| PDK1_MOUSE | 0.93729388 | 0.14992757 |
| PDK2_MOUSE | 0.93077913 | 0.27097819 |
| PDK3_MOUSE | 0.93361175 | 0.15337018 |
| PDP1_MOUSE | 1.06257359 | 0.28100377 |

| | | |
|---|---|---|
| PDPR_MOUSE | 1.00762373 | 0.96004196 |
| PDXK_MOUSE | 1.10611712 | 0.36480152 |
| PEA15_MOUSE | 1.36824969 | 0.61901669 |
| PEBP1_MOUSE | 0.93651747 | 0.0171186 |
| PEX5R_MOUSE | 0.97945021 | 0.47805376 |
| PFD3_MOUSE | 1.02434824 | 0.92730139 |
| PFD5_MOUSE | 0.96707755 | 0.3138578 |
| PFD6_MOUSE | 0.96528723 | 0.36871092 |
| PFKAL_MOUSE | 1.05272213 | 0.32877523 |
| PFKAM_MOUSE | 0.98169945 | 0.58441908 |
| PFKAP_MOUSE | 1.02725679 | 0.61691882 |
| PGAM1_MOUSE | 0.99935061 | 0.81874243 |
| PGAM5_MOUSE | 0.92499511 | 0.03368299 |
| PGCB_MOUSE | 1.13612543 | 0.30486573 |
| PGES2_MOUSE | 0.96041077 | 0.45285622 |
| PGFS_MOUSE | 1.06252775 | 0.18184908 |
| PGK1_MOUSE | 1.03335994 | 0.65176088 |
| PGM1_MOUSE | 1.0370261 | 0.64439547 |
| PGM2L_MOUSE | 0.94145415 | 0.07355529 |
| PGP_MOUSE | 0.93352441 | 0.06661916 |
| PGPS1_MOUSE | 1.05431775 | 0.45114758 |
| PGRC1_MOUSE | 0.96979417 | 0.50456131 |
| PGRC2_MOUSE | 0.89370458 | 0.06610713 |
| PGTA_MOUSE | 1.02648338 | 0.77741789 |
| PHAR1_MOUSE | 0.93084472 | 0.05922947 |
| PHB_MOUSE | 0.9865244 | 0.48785354 |
| PHB2_MOUSE | 1.06724223 | 0.33035951 |
| PHF24_MOUSE | 1.07333765 | 0.56219408 |
| PHIPL_MOUSE | 0.95514206 | 0.23890019 |
| PHP14_MOUSE | 0.9886719 | 0.595984 |
| PHYIP_MOUSE | 0.96376064 | 0.4341621 |
| PI3R4_MOUSE | 0.89167808 | 0.02270654 |
| PI42A_MOUSE | 0.94273072 | 0.10806918 |
| PI42B_MOUSE | 0.97617478 | 0.48190734 |
| PI42C_MOUSE | 1.00358304 | 0.80108792 |
| PI4KA_MOUSE | 1.05584289 | 0.3535877 |
| PI51C_MOUSE | 0.96523372 | 0.50100504 |
| PICAL_MOUSE | 0.96834221 | 0.38879483 |
| PIMT_MOUSE | 0.92375817 | 0.08782141 |

| | | |
|---|---|---|
| PIN1_MOUSE | 1.06597268 | 0.44592309 |
| PIPNA_MOUSE | 0.98535021 | 0.61805095 |
| PIPNB_MOUSE | 1.02701707 | 0.88147985 |
| PITM1_MOUSE | 0.94288814 | 0.16538686 |
| PK3C3_MOUSE | 1.04634065 | 0.73536577 |
| PKP4_MOUSE | 1.03996398 | 0.57660734 |
| PLAP_MOUSE | 0.9774675 | 0.57516115 |
| PLBL2_MOUSE | 0.84508706 | 0.01699015 |
| PLCB1_MOUSE | 1.00613749 | 0.77020676 |
| PLCG1_MOUSE | 1.29537503 | 0.08387851 |
| PLCX3_MOUSE | 1.14424476 | 0.12723393 |
| PLD3_MOUSE | 0.97168916 | 0.48260632 |
| PLEC_MOUSE | 1.07899567 | 0.2027478 |
| PLPL8_MOUSE | 1.05821826 | 0.20074931 |
| PLPP_MOUSE | 1.04769601 | 0.42258384 |
| PLPP3_MOUSE | 1.15348835 | 0.03520309 |
| PLPR4_MOUSE | 0.98114661 | 0.61911718 |
| PLRKT_MOUSE | 0.99214784 | 0.75749344 |
| PLSL_MOUSE | 0.92644802 | 0.13896414 |
| PLST_MOUSE | 0.97279865 | 0.46286432 |
| PLXA1_MOUSE | 1.02748703 | 0.5906007 |
| PLXA4_MOUSE | 1.00085776 | 0.94848269 |
| PLXB2_MOUSE | 1.17344504 | 0.33905113 |
| PMM1_MOUSE | 0.93177184 | 0.31012612 |
| PNPH_MOUSE | 0.98586192 | 0.71419673 |
| PNPT1_MOUSE | 0.96942518 | 0.43589907 |
| PP1A_MOUSE | 1.0420291 | 0.44524157 |
| PP1R7_MOUSE | 0.9492785 | 0.23359999 |
| PP2AA_MOUSE | 0.98023118 | 0.57012664 |
| PP2BA_MOUSE | 0.95650745 | 0.1996156 |
| PP2BB_MOUSE | 0.92236217 | 0.06922296 |
| PPAC_MOUSE | 0.89862769 | 0.08841646 |
| PPCE_MOUSE | 0.88941128 | 0.11350971 |
| PPCEL_MOUSE | 0.91241081 | 0.20774792 |
| PPGB_MOUSE | 0.99277351 | 0.73645119 |
| PPIA_MOUSE | 0.9050587 | 0.01508338 |
| PPIB_MOUSE | 1.00714709 | 0.9466152 |
| PPID_MOUSE | 1.01164918 | 0.71325911 |
| PPM1A_MOUSE | 1.05648944 | 0.86269732 |

| | | |
|---|---|---|
| PPM1H_MOUSE | 1.04288989 | 0.74915372 |
| PPME1_MOUSE | 0.94068711 | 0.10783277 |
| PPP5_MOUSE | 0.98668627 | 0.6034343 |
| PPR21_MOUSE | 0.91015258 | 0.24976518 |
| PPR29_MOUSE | 0.95123548 | 0.15439529 |
| PPTC7_MOUSE | 0.99441218 | 0.69922397 |
| PRAF3_MOUSE | 1.22893042 | 0.00097627 |
| PRDX1_MOUSE | 0.99648674 | 0.7490284 |
| PRDX2_MOUSE | 1.0048277 | 0.94308113 |
| PRDX3_MOUSE | 1.04002467 | 0.68350605 |
| PRDX5_MOUSE | 0.99238561 | 0.64325564 |
| PRDX6_MOUSE | 1.06887752 | 0.07476094 |
| PREB_MOUSE | 1.0305386 | 0.76752477 |
| PREP_MOUSE | 1.00554792 | 0.96084984 |
| PREX1_MOUSE | 1.07314831 | 0.27822383 |
| PRIO_MOUSE | 1.2152459 | 0.31692789 |
| PROD_MOUSE | 1.33184912 | 0.00683403 |
| PROF1_MOUSE | 0.99089886 | 0.68098301 |
| PROF2_MOUSE | 0.97319225 | 0.32864379 |
| PRPS1_MOUSE | 1.19313586 | 0.23268953 |
| PRPTZ_MOUSE | 0.94046504 | 0.32248869 |
| PRRT2_MOUSE | 0.93747582 | 0.10437303 |
| PRRT3_MOUSE | 0.98371067 | 0.59608915 |
| PRS10_MOUSE | 0.93512359 | 0.29038965 |
| PRS4_MOUSE | 1.02088613 | 0.85935263 |
| PRS6A_MOUSE | 0.91731684 | 0.17566909 |
| PRS6B_MOUSE | 1.00362165 | 0.86879271 |
| PRS7_MOUSE | 1.0002942 | 0.78615279 |
| PRS8_MOUSE | 1.02220178 | 0.75874328 |
| PSA_MOUSE | 1.00303462 | 0.89270577 |
| PSA1_MOUSE | 1.02998195 | 0.69003613 |
| PSA2_MOUSE | 1.08071953 | 0.61203594 |
| PSA3_MOUSE | 1.03955006 | 0.41789878 |
| PSA4_MOUSE | 0.97312791 | 0.42161853 |
| PSA5_MOUSE | 1.02069557 | 0.43074498 |
| PSA6_MOUSE | 0.97606106 | 0.35994854 |
| PSA7_MOUSE | 0.95031584 | 0.30712505 |
| PSB1_MOUSE | 0.97220788 | 0.53312792 |
| PSB2_MOUSE | 1.02259758 | 0.70455866 |

| | | |
|---|---|---|
| PSB3_MOUSE | 1.03650417 | 0.60703967 |
| PSB4_MOUSE | 1.09031281 | 0.3767386 |
| PSB5_MOUSE | 1.06300824 | 0.54682094 |
| PSB7_MOUSE | 1.14197811 | 0.2033687 |
| PSD11_MOUSE | 0.98031312 | 0.55340841 |
| PSD12_MOUSE | 0.95514434 | 0.41985889 |
| PSD13_MOUSE | 1.02813493 | 0.65293081 |
| PSD3_MOUSE | 0.96496766 | 0.20095554 |
| PSMD1_MOUSE | 1.016873 | 0.79713294 |
| PSMD2_MOUSE | 1.00111208 | 0.88348815 |
| PSMD3_MOUSE | 1.02822311 | 0.36964501 |
| PSMD4_MOUSE | 0.90764278 | 0.09602632 |
| PSMD5_MOUSE | 1.09648268 | 0.71265602 |
| PSMD6_MOUSE | 1.10318572 | 0.03722242 |
| PSMD8_MOUSE | 0.96818102 | 0.56338811 |
| PSMD9_MOUSE | 1.00634363 | 0.87850663 |
| PSME1_MOUSE | 0.85349389 | 0.02720591 |
| PTCD3_MOUSE | 0.92077942 | 0.07914372 |
| PTGR3_MOUSE | 0.98930205 | 0.77100447 |
| PTH2_MOUSE | 1.40128853 | 0.03860765 |
| PTN11_MOUSE | 1.11394948 | 0.35950536 |
| PTN5_MOUSE | 1.02792282 | 0.75957576 |
| PTPA_MOUSE | 0.91739892 | 0.01995569 |
| PTPR2_MOUSE | 1.05497927 | 0.21468762 |
| PTPRA_MOUSE | 0.95736181 | 0.36575842 |
| PTPRD_MOUSE | 1.00970418 | 0.98280693 |
| PTPRS_MOUSE | 0.97560704 | 0.46096821 |
| PUR6_MOUSE | 0.96800801 | 0.44984594 |
| PUR9_MOUSE | 1.04132235 | 0.84339459 |
| PURA_MOUSE | 0.9363025 | 0.20114416 |
| PURB_MOUSE | 1.05287647 | 0.71801766 |
| PYC_MOUSE | 1.08013427 | 0.13587142 |
| PYGB_MOUSE | 1.08392465 | 0.01500789 |
| PYGM_MOUSE | 1.112087 | 0.02193498 |
| QCR1_MOUSE | 1.01169733 | 0.98801884 |
| QCR2_MOUSE | 0.98791128 | 0.61404823 |
| QCR7_MOUSE | 0.95733574 | 0.25947284 |
| QCR8_MOUSE | 1.09664253 | 0.65418238 |
| QCR9_MOUSE | 1.15712494 | 0.12183419 |

| | | |
|---|---|---|
| QORL2_MOUSE | 1.44243371 | 0.1082091 |
| RAB10_MOUSE | 1.20621044 | 0.09095565 |
| RAB12_MOUSE | 1.08023048 | 0.04125747 |
| RAB14_MOUSE | 0.99635397 | 0.79720283 |
| RAB18_MOUSE | 1.00468172 | 0.70458126 |
| RAB1A_MOUSE | 0.90731352 | 0.06284606 |
| RAB1B_MOUSE | 0.95186358 | 0.18105308 |
| RAB21_MOUSE | 1.07540456 | 0.5658942 |
| RAB23_MOUSE | 1.01080441 | 0.91200033 |
| RAB2A_MOUSE | 0.98936085 | 0.59449412 |
| RAB35_MOUSE | 0.99362191 | 0.77694908 |
| RAB3A_MOUSE | 0.97120894 | 0.49888111 |
| RAB3B_MOUSE | 1.07653686 | 0.68951256 |
| RAB3C_MOUSE | 1.0447865 | 0.98895973 |
| RAB4B_MOUSE | 0.97944705 | 0.57335125 |
| RAB5A_MOUSE | 1.08860547 | 0.27483661 |
| RAB5B_MOUSE | 0.94031537 | 0.1614421 |
| RAB5C_MOUSE | 1.01405013 | 0.83503063 |
| RAB6A_MOUSE | 0.97803469 | 0.57123781 |
| RAB6B_MOUSE | 0.96473644 | 0.38382059 |
| RAB7A_MOUSE | 1.00676915 | 0.83160283 |
| RAB8A_MOUSE | 1.67570226 | 0.06205517 |
| RABE1_MOUSE | 0.85418023 | 0.05176806 |
| RABL6_MOUSE | 0.94818092 | 0.42021216 |
| RAC1_MOUSE | 1.0583443 | 0.83299825 |
| RACK1_MOUSE | 0.97590529 | 0.59709999 |
| RADI_MOUSE | 1.01189008 | 0.87431125 |
| RALA_MOUSE | 1.08575762 | 0.46861418 |
| RAN_MOUSE | 0.98695635 | 0.5309179 |
| RANG_MOUSE | 0.88422938 | 0.05793791 |
| RAP2A_MOUSE | 1.03196459 | 0.66591793 |
| RAP2B_MOUSE | 0.97326508 | 0.45540521 |
| RASH_MOUSE | 0.95613379 | 0.38487808 |
| RASL1_MOUSE | 0.99970923 | 0.89218374 |
| RASM_MOUSE | 1.10863888 | 0.14557324 |
| RB11B_MOUSE | 1.04332353 | 0.32837907 |
| RB39B_MOUSE | 0.99731684 | 0.7510797 |
| RB6I2_MOUSE | 0.92540326 | 0.11513112 |
| RBBP9_MOUSE | 1.05063415 | 0.73727939 |

| | | |
|---|---|---|
| RBGPR_MOUSE | 0.95931623 | 0.28753297 |
| RCN2_MOUSE | 0.86323993 | 0.19989303 |
| RD23B_MOUSE | 1.03184009 | 0.88407089 |
| RDH14_MOUSE | 1.01932945 | 0.81068441 |
| RENT1_MOUSE | 1.13651544 | 0.32414873 |
| RFIP2_MOUSE | 0.97744292 | 0.53541153 |
| RFIP5_MOUSE | 0.82827197 | 0.004972 |
| RFTN2_MOUSE | 1.18123722 | 0.47660201 |
| RGRF2_MOUSE | 0.95621213 | 0.45574269 |
| RGS7_MOUSE | 1.06613564 | 0.45796778 |
| RHG01_MOUSE | 1.00414306 | 0.97513123 |
| RHG32_MOUSE | 0.99949117 | 0.83417063 |
| RHG35_MOUSE | 1.12397969 | 0.1681792 |
| RHG44_MOUSE | 0.99536762 | 0.74758901 |
| RHOA_MOUSE | 1.17126322 | 0.02563162 |
| RHOB_MOUSE | 1.13512493 | 0.03238656 |
| RHOG_MOUSE | 0.93688861 | 0.36250404 |
| RIMB2_MOUSE | 0.91544582 | 0.01857911 |
| RIMS1_MOUSE | 0.99564106 | 0.80008711 |
| RINI_MOUSE | 0.95266173 | 0.42019188 |
| RL10_MOUSE | 0.96034669 | 0.3751769 |
| RL10A_MOUSE | 1.00142439 | 0.98769481 |
| RL12_MOUSE | 0.95056575 | 0.34171317 |
| RL13_MOUSE | 1.28649532 | 0.00542686 |
| RL14_MOUSE | 1.00173738 | 0.93064614 |
| RL17_MOUSE | 1.13475299 | 0.01307892 |
| RL21_MOUSE | 0.9770437 | 0.56401543 |
| RL3_MOUSE | 1.05475442 | 0.1105932 |
| RL38_MOUSE | 0.96130319 | 0.51849438 |
| RL4_MOUSE | 1.02406662 | 0.76887408 |
| RL5_MOUSE | 1.01958825 | 0.87470122 |
| RL6_MOUSE | 1.04220487 | 0.48659376 |
| RL7_MOUSE | 1.0676172 | 0.40007301 |
| RL7A_MOUSE | 0.96774819 | 0.44636191 |
| RL8_MOUSE | 1.0821383 | 0.34926238 |
| RLA0_MOUSE | 0.8992624 | 0.01106995 |
| RLA1_MOUSE | 1.01523076 | 0.88680788 |
| RLA2_MOUSE | 0.87789947 | 0.16794652 |
| RM01_MOUSE | 1.2219823 | 0.04654935 |

| | | |
|---|---|---|
| RM12_MOUSE | 0.85648028 | 0.0520232 |
| RM41_MOUSE | 0.87317058 | 0.03167622 |
| RM50_MOUSE | 0.89149707 | 0.03190142 |
| RMD1_MOUSE | 0.92883811 | 0.15227402 |
| RMD3_MOUSE | 0.94254166 | 0.15314214 |
| RO60_MOUSE | 1.11781037 | 0.68018988 |
| ROBO2_MOUSE | 0.82465028 | 0.0682884 |
| ROCK2_MOUSE | 0.95463743 | 0.39349243 |
| ROGDI_MOUSE | 1.0133779 | 0.99429488 |
| RP3A_MOUSE | 1.04420423 | 0.78576917 |
| RPGF2_MOUSE | 0.8806987 | 0.00810043 |
| RPGF4_MOUSE | 0.99700422 | 0.83229983 |
| RPGP1_MOUSE | 1.00151786 | 0.65251196 |
| RPGP2_MOUSE | 0.95419559 | 0.2416302 |
| RPN1_MOUSE | 1.05715667 | 0.595989 |
| RPN2_MOUSE | 1.04552002 | 0.35340898 |
| RRAS2_MOUSE | 0.98899762 | 0.6486666 |
| RRFM_MOUSE | 1.01450603 | 0.90289726 |
| RS10_MOUSE | 1.06915766 | 0.43863917 |
| RS11_MOUSE | 1.05037568 | 0.50982213 |
| RS13_MOUSE | 1.10662193 | 0.23947258 |
| RS14_MOUSE | 1.01254505 | 0.94049481 |
| RS15A_MOUSE | 1.16267155 | 0.03871847 |
| RS18_MOUSE | 1.03483303 | 0.67342642 |
| RS19_MOUSE | 0.99985735 | 0.72847647 |
| RS2_MOUSE | 1.03587276 | 0.43573595 |
| RS21_MOUSE | 0.92580756 | 0.17150523 |
| RS24_MOUSE | 0.86541691 | 0.05371257 |
| RS27A_MOUSE | 0.98427149 | 0.58368844 |
| RS27L_MOUSE | 1.07386022 | 0.19775328 |
| RS3_MOUSE | 0.99402437 | 0.68947858 |
| RS3A_MOUSE | 0.97666258 | 0.58078618 |
| RS4X_MOUSE | 1.1468839 | 0.03459434 |
| RS6_MOUSE | 1.02620985 | 0.75940861 |
| RS7_MOUSE | 1.31546195 | 0.06390383 |
| RS8_MOUSE | 0.95447842 | 0.22542809 |
| RS9_MOUSE | 1.13896196 | 0.02750796 |
| RSSA_MOUSE | 0.99236724 | 0.70690093 |
| RT23_MOUSE | 1.01594625 | 0.82428655 |

| | | |
|---|---|---|
| RT27_MOUSE | 0.97009438 | 0.57369042 |
| RT29_MOUSE | 1.39907581 | 0.10900803 |
| RT34_MOUSE | 0.82251234 | 0.06443731 |
| RT36_MOUSE | 1.14315833 | 0.05350938 |
| RT4I1_MOUSE | 1.15681922 | 0.1500766 |
| RTCB_MOUSE | 1.01697094 | 0.59779571 |
| RTN1_MOUSE | 0.9446657 | 0.24741342 |
| RTN3_MOUSE | 0.88792286 | 0.01867472 |
| RTN4_MOUSE | 0.93562716 | 0.19360728 |
| RUFY3_MOUSE | 0.90360467 | 0.03603376 |
| RUVB1_MOUSE | 0.85495002 | 0.12017311 |
| RYR2_MOUSE | 0.96981415 | 0.45958464 |
| S100B_MOUSE | 1.07615757 | 0.79393762 |
| S10AD_MOUSE | 0.98142271 | 0.58792829 |
| S12A2_MOUSE | 0.96411232 | 0.36278421 |
| S12A5_MOUSE | 1.08186295 | 0.21279942 |
| S14L2_MOUSE | 1.31982417 | 0.03278987 |
| S1PR1_MOUSE | 1.07335572 | 0.6308228 |
| S20A2_MOUSE | 1.18114112 | 0.02850308 |
| S2542_MOUSE | 1.23876352 | 0.03489096 |
| S2546_MOUSE | 1.05755188 | 0.36558275 |
| S2551_MOUSE | 1.02868958 | 0.58016194 |
| S27A4_MOUSE | 0.99510606 | 0.83680909 |
| S38A3_MOUSE | 1.05171259 | 0.70640212 |
| S39AA_MOUSE | 0.93959314 | 0.40388843 |
| S4A10_MOUSE | 1.08297921 | 0.0026341 |
| S4A4_MOUSE | 1.03002366 | 0.45470768 |
| S6A11_MOUSE | 1.04479066 | 0.67881269 |
| S6A17_MOUSE | 0.98198456 | 0.55952686 |
| S7A14_MOUSE | 1.07678813 | 0.41793214 |
| SAC1_MOUSE | 1.05463631 | 0.25844817 |
| SAHH_MOUSE | 0.97767876 | 0.4836327 |
| SAHH2_MOUSE | 1.04947005 | 0.23387923 |
| SAHH3_MOUSE | 1.20571249 | 0.00609146 |
| SAM50_MOUSE | 1.01565246 | 0.8108083 |
| SAR1B_MOUSE | 1.01195537 | 0.90494495 |
| SATT_MOUSE | 1.27179318 | 0.00018258 |
| SC22B_MOUSE | 1.04644217 | 0.84820811 |
| SC23A_MOUSE | 1.02069283 | 0.70997051 |

| | | |
|---|---|---|
| SC6A1_MOUSE | 1.04704687 | 0.44402493 |
| SC6A9_MOUSE | 1.04501706 | 0.99659996 |
| SCAI_MOUSE | 1.00814605 | 0.96001856 |
| SCAM1_MOUSE | 0.91149406 | 0.05312665 |
| SCAM3_MOUSE | 0.95695415 | 0.37448447 |
| SCAM5_MOUSE | 1.00947592 | 0.97083037 |
| SCFD1_MOUSE | 0.90936702 | 0.10148419 |
| SCMC2_MOUSE | 1.15307934 | 0.00565949 |
| SCMC3_MOUSE | 0.99125703 | 0.6576777 |
| SCN1A_MOUSE | 0.96107431 | 0.32573638 |
| SCN2B_MOUSE | 0.91962198 | 0.04399186 |
| SCN9A_MOUSE | 0.98267841 | 0.54947868 |
| SCOT1_MOUSE | 0.97196405 | 0.32820464 |
| SCPDL_MOUSE | 0.96689258 | 0.38797649 |
| SCRN1_MOUSE | 0.89931972 | 0.01521 |
| SDHA_MOUSE | 1.01064141 | 0.97560476 |
| SDHB_MOUSE | 1.01242014 | 0.87286453 |
| SEM4A_MOUSE | 1.2110218 | 0.22946015 |
| SEPT11_MOUSE | 1.02676427 | 0.89044176 |
| SEPT2_MOUSE | 1.10808339 | 0.03104252 |
| SEPT3_MOUSE | 0.98088199 | 0.43153585 |
| SEPT4_MOUSE | 0.93352012 | 0.34317225 |
| SEPT5_MOUSE | 0.98490978 | 0.53662693 |
| SEPT6_MOUSE | 0.98069654 | 0.49533197 |
| SEPT7_MOUSE | 0.97871305 | 0.56888885 |
| SEPT8_MOUSE | 1.05738837 | 0.3594926 |
| SEPT9_MOUSE | 0.96973026 | 0.16991516 |
| SERA_MOUSE | 0.92761969 | 0.08067915 |
| SERC_MOUSE | 0.98622933 | 0.66072066 |
| SFXN1_MOUSE | 1.06569193 | 0.36873902 |
| SFXN3_MOUSE | 0.98838765 | 0.60074401 |
| SFXN5_MOUSE | 1.05890098 | 0.59976321 |
| SGIP1_MOUSE | 1.02558421 | 0.63590875 |
| SGSM1_MOUSE | 1.01947642 | 0.8455668 |
| SGT1_MOUSE | 1.00218841 | 0.70255641 |
| SGTB_MOUSE | 1.04878091 | 0.50861808 |
| SH3G1_MOUSE | 0.81574453 | 0.05437591 |
| SH3G2_MOUSE | 0.84639204 | 0.01048259 |
| SH3G3_MOUSE | 0.82058712 | 0.01296663 |

| | | |
|---|---|---|
| SH3K1_MOUSE | 0.85272449 | 0.10281332 |
| SH3L3_MOUSE | 1.07402202 | 0.66597029 |
| SHAN1_MOUSE | 0.98526062 | 0.57276709 |
| SHAN2_MOUSE | 0.96519912 | 0.36678548 |
| SHAN3_MOUSE | 0.96995783 | 0.4804189 |
| SHLB1_MOUSE | 0.90725642 | 0.06069977 |
| SHLB2_MOUSE | 0.92642359 | 0.11529593 |
| SHPS1_MOUSE | 1.07856662 | 0.152269 |
| SHSA7_MOUSE | 1.00105769 | 0.78460967 |
| SI1L1_MOUSE | 0.957487 | 0.3542715 |
| SIR2_MOUSE | 0.95892558 | 0.57611729 |
| SIR3_MOUSE | 1.12625971 | 0.27961822 |
| SIR5_MOUSE | 1.09770497 | 0.57014935 |
| SKP1_MOUSE | 0.91001624 | 0.09055613 |
| SKT_MOUSE | 0.99757909 | 0.71681481 |
| SL9A1_MOUSE | 1.03318016 | 0.69987297 |
| SLIRP_MOUSE | 1.03305584 | 0.51598989 |
| SLK_MOUSE | 0.99291245 | 0.64218009 |
| SMAP2_MOUSE | 0.91995148 | 0.04765165 |
| SNAA_MOUSE | 0.93173392 | 0.19459389 |
| SNAB_MOUSE | 0.97627099 | 0.49522152 |
| SNAG_MOUSE | 0.92826636 | 0.06734131 |
| SND1_MOUSE | 1.00211889 | 0.82368544 |
| SNG1_MOUSE | 0.93829235 | 0.08963179 |
| SNG3_MOUSE | 1.11486259 | 0.15167164 |
| SNP25_MOUSE | 0.8594587 | 0.05255507 |
| SNP29_MOUSE | 1.03890399 | 0.483857 |
| SNP47_MOUSE | 1.08553901 | 0.50412404 |
| SNPH_MOUSE | 0.94173791 | 0.14161218 |
| SNX1_MOUSE | 0.92577111 | 0.18448065 |
| SNX12_MOUSE | 1.00808218 | 0.85986006 |
| SNX2_MOUSE | 0.98818691 | 0.66347421 |
| SNX27_MOUSE | 0.95699162 | 0.27947091 |
| SNX3_MOUSE | 1.1333266 | 0.06478907 |
| SNX30_MOUSE | 2.21771871 | 0.0721492 |
| SNX4_MOUSE | 0.97005969 | 0.47716903 |
| SNX5_MOUSE | 1.02008195 | 0.81734289 |
| SODM_MOUSE | 1.08063166 | 0.477939 |
| SOGA3_MOUSE | 0.92122949 | 0.16849101 |

| | | |
|---|---|---|
| SORCN_MOUSE | 0.94608544 | 0.14819572 |
| SPB6_MOUSE | 1.19075801 | 0.07879967 |
| SPCS2_MOUSE | 1.06168626 | 0.72908468 |
| SPG7_MOUSE | 1.0479023 | 0.50632023 |
| SPN90_MOUSE | 0.93930708 | 0.12426371 |
| SPRE_MOUSE | 0.97061794 | 0.33119798 |
| SPRL1_MOUSE | 1.0922769 | 0.94911159 |
| SPRY4_MOUSE | 0.91714582 | 0.18315361 |
| SPTB1_MOUSE | 0.94671099 | 0.29081609 |
| SPTB2_MOUSE | 0.99272696 | 0.68333013 |
| SPTN1_MOUSE | 0.98083127 | 0.5849659 |
| SRBS1_MOUSE | 0.98999119 | 0.61263145 |
| SRBS2_MOUSE | 1.05058638 | 0.56344619 |
| SRC_MOUSE | 0.99456918 | 0.64609853 |
| SRC8_MOUSE | 0.9315716 | 0.09193389 |
| SRCN1_MOUSE | 0.9731187 | 0.40206964 |
| SRGP2_MOUSE | 0.93186495 | 0.17842562 |
| SRGP3_MOUSE | 0.9410716 | 0.19740883 |
| SRR_MOUSE | 1.00273983 | 0.78911629 |
| SSBP_MOUSE | 1.02858033 | 0.96214633 |
| SSDH_MOUSE | 1.05336459 | 0.48457141 |
| ST4A1_MOUSE | 1.24981809 | 0.14885465 |
| STAM1_MOUSE | 0.96401984 | 0.28355118 |
| STB5L_MOUSE | 1.03537653 | 0.81945629 |
| STIP1_MOUSE | 0.90087173 | 0.02015739 |
| STK24_MOUSE | 1.09275702 | 0.50911679 |
| STK39_MOUSE | 0.9710908 | 0.43591398 |
| STML2_MOUSE | 1.03179651 | 0.87648485 |
| STMN1_MOUSE | 0.72510624 | 1.2867E-07 |
| STRAP_MOUSE | 0.94190064 | 0.36226366 |
| STRN_MOUSE | 0.94415223 | 0.23880527 |
| STRN3_MOUSE | 0.87127657 | 0.0786812 |
| STRN4_MOUSE | 0.94730094 | 0.27475718 |
| STX12_MOUSE | 0.90608217 | 0.18293903 |
| STX1A_MOUSE | 0.94308441 | 0.27267773 |
| STX1B_MOUSE | 0.92815793 | 0.21488608 |
| STX7_MOUSE | 0.96880581 | 0.47772908 |
| STXB1_MOUSE | 1.05753936 | 0.35767935 |
| STXB3_MOUSE | 0.893382 | 0.24011521 |

| | | |
|---|---|---|
| STXB5_MOUSE | 0.99450876 | 0.71058841 |
| SUCA_MOUSE | 1.06217273 | 0.5478979 |
| SUCB1_MOUSE | 1.021981 | 0.77281985 |
| SUCB2_MOUSE | 1.05904375 | 0.26430523 |
| SV2A_MOUSE | 1.024776 | 0.81296736 |
| SV2B_MOUSE | 0.95634854 | 0.27412844 |
| SVOP_MOUSE | 0.94044311 | 0.25413697 |
| SYAC_MOUSE | 0.92403912 | 0.02220938 |
| SYDC_MOUSE | 1.04803505 | 0.79129583 |
| SYEP_MOUSE | 0.93075566 | 0.09574481 |
| SYGP1_MOUSE | 0.95533963 | 0.29042545 |
| SYHC_MOUSE | 0.90943028 | 0.07775275 |
| SYIM_MOUSE | 1.03859941 | 0.58928987 |
| SYJ2B_MOUSE | 0.98332061 | 0.5946796 |
| SYK_MOUSE | 1.33450634 | 0.61688277 |
| SYLM_MOUSE | 0.97177975 | 0.47096078 |
| SYMM_MOUSE | 1.09149155 | 0.16359392 |
| SYN1_MOUSE | 1.03637882 | 0.58926886 |
| SYN2_MOUSE | 1.0596935 | 0.64906204 |
| SYN3_MOUSE | 1.07163465 | 0.16762455 |
| SYNC_MOUSE | 0.94703939 | 0.26854893 |
| SYNE1_MOUSE | 0.92456657 | 0.072385 |
| SYNJ1_MOUSE | 1.02113633 | 0.85137508 |
| SYNPO_MOUSE | 0.99553727 | 0.84741596 |
| SYNPR_MOUSE | 0.96311067 | 0.36526735 |
| SYPH_MOUSE | 0.9845774 | 0.49400302 |
| SYRC_MOUSE | 1.0973343 | 0.20710731 |
| SYSC_MOUSE | 1.03284382 | 0.71494256 |
| SYSM_MOUSE | 1.04989415 | 0.71792161 |
| SYT1_MOUSE | 0.98533916 | 0.52971826 |
| SYT12_MOUSE | 0.92375458 | 0.03499333 |
| SYT2_MOUSE | 0.9861662 | 0.76967211 |
| SYT3_MOUSE | 0.98845584 | 0.73235031 |
| SYT7_MOUSE | 1.04949445 | 0.61120816 |
| SYTC_MOUSE | 1.00540656 | 0.84947119 |
| SYTM_MOUSE | 1.23043861 | 0.36645254 |
| SYUA_MOUSE | 0.85037165 | 0.01035041 |
| SYUB_MOUSE | 0.8592758 | 0.01747851 |
| SYVC_MOUSE | 0.94623633 | 0.45948154 |

| | | |
|---|---|---|
| SYWC_MOUSE | 1.04217102 | 0.67100351 |
| SYYM_MOUSE | 1.13322252 | 0.05768868 |
| T11L1_MOUSE | 0.9461901 | 0.42538745 |
| TACO1_MOUSE | 1.06659555 | 0.68833384 |
| TAGL3_MOUSE | 1.01180859 | 0.85287693 |
| TALDO_MOUSE | 0.88202615 | 0.02335911 |
| TANC2_MOUSE | 0.94793374 | 0.33942391 |
| TAU_MOUSE | 0.92624884 | 0.22015272 |
| TB10B_MOUSE | 0.94753702 | 0.3484834 |
| TBA4A_MOUSE | 0.94867859 | 0.31322753 |
| TBB3_MOUSE | 1.09699947 | 0.6596472 |
| TBB4A_MOUSE | 0.90742209 | 0.16494622 |
| TBB5_MOUSE | 0.88218878 | 0.01988568 |
| TBRG4_MOUSE | 0.99649289 | 0.86518266 |
| TCPA_MOUSE | 0.98982054 | 0.69752925 |
| TCPB_MOUSE | 0.98585037 | 0.66673882 |
| TCPD_MOUSE | 1.00836233 | 0.88921639 |
| TCPE_MOUSE | 1.04022588 | 0.49950504 |
| TCPG_MOUSE | 0.99198598 | 0.72968721 |
| TCPH_MOUSE | 0.97792546 | 0.41557343 |
| TCPQ_MOUSE | 0.98445085 | 0.47690159 |
| TCPR1_MOUSE | 0.95944132 | 0.22282383 |
| TCPZ_MOUSE | 0.99504278 | 0.7495115 |
| TCTP_MOUSE | 0.89399001 | 0.16265678 |
| TDRKH_MOUSE | 0.96790514 | 0.37029702 |
| TEBP_MOUSE | 1.00688019 | 0.96098863 |
| TECR_MOUSE | 0.94458914 | 0.22841347 |
| TENA_MOUSE | 1.22311041 | 0.50258319 |
| TENR_MOUSE | 0.93551093 | 0.15694335 |
| TERA_MOUSE | 1.01049795 | 0.95930683 |
| TFAM_MOUSE | 0.92913588 | 0.26154215 |
| TFR1_MOUSE | 0.87528874 | 0.02695667 |
| THEM4_MOUSE | 1.00142319 | 0.85353249 |
| THEM6_MOUSE | 1.11408056 | 0.97465368 |
| THIC_MOUSE | 1.00423726 | 0.99748874 |
| THIKA_MOUSE | 1.04476178 | 0.44730613 |
| THIL_MOUSE | 0.98463867 | 0.55561864 |
| THIM_MOUSE | 1.02418541 | 0.91047678 |
| THNS1_MOUSE | 0.96415427 | 0.37261571 |

| | | |
|---|---|---|
| THOP1_MOUSE | 0.90023 | 0.1180073 |
| THTM_MOUSE | 1.04462611 | 0.40216014 |
| THTR_MOUSE | 1.1169525 | 0.22702532 |
| THY1_MOUSE | 1.06909276 | 0.54064922 |
| TIM10_MOUSE | 0.96762335 | 0.56359936 |
| TIM13_MOUSE | 0.90944571 | 0.01948205 |
| TIM29_MOUSE | 1.11820605 | 0.06579198 |
| TIM44_MOUSE | 0.9690974 | 0.40315824 |
| TIM50_MOUSE | 0.94690242 | 0.05604746 |
| TKT_MOUSE | 0.9640878 | 0.26778813 |
| TLN2_MOUSE | 0.92267038 | 0.02838861 |
| TM160_MOUSE | 1.06120632 | 0.28164074 |
| TM1L2_MOUSE | 0.91096165 | 0.04149846 |
| TMM11_MOUSE | 0.99971579 | 0.9235086 |
| TMM33_MOUSE | 1.11270596 | 0.12955369 |
| TMM65_MOUSE | 0.99989303 | 0.76880802 |
| TMOD2_MOUSE | 1.00595735 | 0.93766663 |
| TMX2_MOUSE | 0.99040111 | 0.77287943 |
| TMX3_MOUSE | 1.00623572 | 0.95218588 |
| TMX4_MOUSE | 1.02786535 | 0.99878993 |
| TNPO3_MOUSE | 1.08265063 | 0.51795623 |
| TOLIP_MOUSE | 0.88994856 | 0.02700395 |
| TOM1_MOUSE | 1.0782633 | 0.30007052 |
| TOM22_MOUSE | 0.87148339 | 0.00639301 |
| TOM40_MOUSE | 0.92871867 | 0.31824035 |
| TOM70_MOUSE | 0.93413348 | 0.0500182 |
| TP4A2_MOUSE | 1.26450248 | 0.08853068 |
| TPC_MOUSE | 0.85941666 | 0.03571952 |
| TPD52_MOUSE | 0.88589387 | 0.05024765 |
| TPD54_MOUSE | 0.97256609 | 0.50168166 |
| TPIS_MOUSE | 0.96156479 | 0.30606936 |
| TPM1_MOUSE | 0.82158649 | 0.07685962 |
| TPM3_MOUSE | 0.83181788 | 0.12636424 |
| TPP2_MOUSE | 0.96435444 | 0.31314126 |
| TPPP_MOUSE | 0.98983512 | 0.74011714 |
| TPPP3_MOUSE | 0.87549819 | 0.00034584 |
| TPRGL_MOUSE | 1.00638025 | 0.93669913 |
| TRAP1_MOUSE | 1.0252592 | 0.79771354 |
| TRFE_MOUSE | 0.86161976 | 0.07740691 |

| | | |
|---|---|---|
| TRIM2_MOUSE | 0.99869517 | 0.76358175 |
| TRIM3_MOUSE | 0.90037685 | 0.00616791 |
| TRIO_MOUSE | 0.98451337 | 0.57920402 |
| TRPV2_MOUSE | 1.06253942 | 0.75566225 |
| TRXR1_MOUSE | 1.182234 | 0.09563947 |
| TRXR2_MOUSE | 1.10992567 | 0.30197998 |
| TS101_MOUSE | 0.9987235 | 0.8092808 |
| TSN_MOUSE | 0.85806948 | 0.01495171 |
| TSN2_MOUSE | 0.78755327 | 0.01246341 |
| TSN7_MOUSE | 1.02855261 | 0.94388463 |
| TSR2_MOUSE | 0.78993627 | 0.01591154 |
| TTC19_MOUSE | 1.08804145 | 0.48822944 |
| TTC7B_MOUSE | 1.03321879 | 0.38739963 |
| TTYH1_MOUSE | 1.09244013 | 0.35325209 |
| TTYH3_MOUSE | 1.98762773 | 0.06407048 |
| TWF1_MOUSE | 0.96596414 | 0.3479013 |
| TWF2_MOUSE | 1.0414985 | 0.28373539 |
| TXD12_MOUSE | 0.90612918 | 0.21537386 |
| TXND5_MOUSE | 0.95008826 | 0.27725032 |
| TXNL1_MOUSE | 0.96188283 | 0.29792632 |
| TXTP_MOUSE | 0.99610194 | 0.83089066 |
| TY3H_MOUSE | 0.96575574 | 0.52756912 |
| UB2L3_MOUSE | 1.05690793 | 0.3724886 |
| UB2V2_MOUSE | 1.04748754 | 0.60753069 |
| UBA1_MOUSE | 0.95168029 | 0.28156258 |
| UBA5_MOUSE | 0.94309139 | 0.23308059 |
| UBC12_MOUSE | 0.95529765 | 0.14945501 |
| UBE2N_MOUSE | 0.99700888 | 0.79835596 |
| UBE2O_MOUSE | 0.99411847 | 0.84077818 |
| UBP14_MOUSE | 0.96358024 | 0.2858734 |
| UBP5_MOUSE | 0.97341633 | 0.38453958 |
| UBP7_MOUSE | 0.95512982 | 0.51509324 |
| UBQL1_MOUSE | 0.91287205 | 0.16683312 |
| UBQL2_MOUSE | 0.96792371 | 0.37771157 |
| UBR4_MOUSE | 1.07291953 | 0.30335954 |
| UBXN6_MOUSE | 1.09404209 | 0.43693386 |
| UCHL1_MOUSE | 0.94324162 | 0.13856474 |
| UCHL3_MOUSE | 0.97188485 | 0.48368161 |
| UCRI_MOUSE | 1.11489258 | 0.08478631 |

| | | |
|---|---|---|
| UGPA_MOUSE | 1.0308431 | 0.88989633 |
| ULA1_MOUSE | 0.98113754 | 0.54432128 |
| UN13A_MOUSE | 1.01711656 | 0.60763339 |
| UQCC1_MOUSE | 0.92407424 | 0.02441843 |
| USMG5_MOUSE | 1.02403423 | 0.93235496 |
| USO1_MOUSE | 0.95990583 | 0.34881409 |
| USP9X_MOUSE | 0.9799593 | 0.40949594 |
| VA0D1_MOUSE | 1.024633 | 0.77378582 |
| VAC14_MOUSE | 1.11176354 | 0.31734126 |
| VAMP1_MOUSE | 0.89368993 | 0.07889834 |
| VAMP2_MOUSE | 0.87252571 | 0.08498467 |
| VAPA_MOUSE | 1.14396064 | 0.08326456 |
| VAPB_MOUSE | 0.97259054 | 0.42178386 |
| VAS1_MOUSE | 1.0109867 | 0.93260932 |
| VAT1_MOUSE | 1.12942398 | 0.02496447 |
| VAT1L_MOUSE | 1.08616339 | 0.0217056 |
| VATA_MOUSE | 0.97969198 | 0.53996233 |
| VATB2_MOUSE | 1.03428276 | 0.63523004 |
| VATC1_MOUSE | 1.01698142 | 0.7358551 |
| VATD_MOUSE | 1.01549774 | 0.91317265 |
| VATE1_MOUSE | 0.9533422 | 0.18390816 |
| VATF_MOUSE | 0.975514 | 0.31551618 |
| VATG1_MOUSE | 0.91381164 | 0.12155224 |
| VATG2_MOUSE | 0.92537756 | 0.12362507 |
| VATH_MOUSE | 1.04035333 | 0.51400887 |
| VATL_MOUSE | 1.43817644 | 0.0089681 |
| VCAM1_MOUSE | 1.1060851 | 0.154121 |
| VCIP1_MOUSE | 0.89452199 | 0.02421264 |
| VDAC1_MOUSE | 0.96794906 | 0.35573662 |
| VDAC2_MOUSE | 0.94635447 | 0.05771806 |
| VDAC3_MOUSE | 0.96134227 | 0.23916539 |
| VGLU1_MOUSE | 1.12942693 | 0.27134059 |
| VGLU2_MOUSE | 0.95270776 | 0.29404936 |
| VIAAT_MOUSE | 1.08785407 | 0.19056714 |
| VIME_MOUSE | 0.82043361 | 0.00221274 |
| VINC_MOUSE | 1.00728389 | 0.9974669 |
| VISL1_MOUSE | 0.91172632 | 0.03859952 |
| VMA5A_MOUSE | 1.04581816 | 0.84178722 |
| VP13A_MOUSE | 1.04297438 | 0.61735275 |

| | | |
|---|---|---|
| VP13C_MOUSE | 0.92050447 | 0.236069 |
| VP26A_MOUSE | 1.01525119 | 0.95772505 |
| VP26B_MOUSE | 1.01425214 | 0.84112908 |
| VPP1_MOUSE | 0.99153302 | 0.66960457 |
| VPS16_MOUSE | 1.10985587 | 0.3395319 |
| VPS29_MOUSE | 1.07236188 | 0.55709807 |
| VPS35_MOUSE | 0.99336806 | 0.67153648 |
| VPS45_MOUSE | 1.07015371 | 0.0585805 |
| VPS50_MOUSE | 1.02366727 | 0.78410892 |
| VPS51_MOUSE | 1.03896638 | 0.61273325 |
| VPS53_MOUSE | 1.20042907 | 0.18981004 |
| VTA1_MOUSE | 0.92807145 | 0.03330006 |
| VWA8_MOUSE | 1.01780504 | 0.90252564 |
| WASF1_MOUSE | 0.96679698 | 0.28621079 |
| WASF3_MOUSE | 0.93731895 | 0.41155407 |
| WASL_MOUSE | 1.24999272 | 0.0501783 |
| WBP2_MOUSE | 1.0204201 | 0.9553125 |
| WDFY3_MOUSE | 1.10767713 | 0.26354883 |
| WDR1_MOUSE | 1.03237583 | 0.56281059 |
| WDR37_MOUSE | 1.05139273 | 0.3096423 |
| WDR44_MOUSE | 0.87542043 | 0.18360738 |
| WDR47_MOUSE | 1.00684118 | 0.93506322 |
| WDR48_MOUSE | 1.01385339 | 0.85738402 |
| WDR7_MOUSE | 0.98605399 | 0.64379707 |
| WFS1_MOUSE | 1.01617449 | 0.87820962 |
| WIPI2_MOUSE | 1.03539309 | 0.4650419 |
| WNK2_MOUSE | 1.12406668 | 0.17475338 |
| XPO1_MOUSE | 1.07081961 | 0.58406158 |
| XPO2_MOUSE | 0.99901198 | 0.81706332 |
| XPO7_MOUSE | 1.06797572 | 0.83036433 |
| XPP1_MOUSE | 0.92329385 | 0.03712718 |
| YKT6_MOUSE | 0.96367688 | 0.32493949 |
| ZC21A_MOUSE | 0.97277742 | 0.48864905 |
| ZNT3_MOUSE | 0.9808544 | 0.46353926 |
| ZNT9_MOUSE | 1.02329844 | 0.64256207 |
| ZO1_MOUSE | 1.11821229 | 0.04610989 |
| ZO2_MOUSE | 0.88341097 | 0.02445254 |

**Table S2. Proteins exhibiting significant changes in WT versus PPT1 KO synaptic palmitome.** These data correspond to **Figure 2B** and are the expression ratios for proteins that exhibit significant changes in the palmitome (n = 242). Proteins marked with an asterisk (*) were removed from subsequent consideration as PPT1 substrates due to decreased palmitoylation (n = 4: HDHD2, S1PR1, SNX1, UBQL2 ; blue <1.5-fold change), presence in the CRAPome (n = 9: 1433E, COF1, HNRPU, KPYM, RL13, RL22, RL4, RLA0, TCPD), lack of a cysteine residue (n = 2: NDUA6, SYUA), increased palmitoylation and protein expression (n = 4: ASAH1, CATD, SCRB2, and TPP1), or lack of detection in the synaptic proteome (n = 19: CKAP4, CPNE1, FHL1, GPC5B, ITM2B, ITM2C, LGI2, MAGI1, NSMA2, PP1G, PRDX4, R7BP, RB3GP, S39AC, TPPC3, VAMP7, XKR4, S1PR1, MYO6, PDPR; removed proteins S1PR1, HNRPU, and RL22 also fall in this category). The remaining 204 proteins are the final list of putative PPT1 substrates prior to the validation screen. P-values were calculated using a two-tailed t-test. Red denotes significantly increased expression in the palmitome (>1.5-fold).

| Uniprot ID | Average Ratio (KO/WT) | P value |
|---|---|---|
| HDHD2_MOUSE* | 0.65809308 | 0.02504326 |
| S1PR1_MOUSE* | 0.43333627 | 0.01690137 |
| SNX1_MOUSE* | 0.55468914 | 0.0240519 |
| UBQL2_MOUSE* | 0.57153189 | 0.01479205 |
| 1433B_MOUSE | 6.16513858 | 0.03447738 |
| 1433E_MOUSE* | 1.7373229 | 0.04223438 |
| 6PGL_MOUSE | 37.011627 | 0.04825557 |
| ABHGA_MOUSE | 1.73812986 | 0.02325394 |
| ABR_MOUSE | 2.06555242 | 0.0190207 |
| ACON_MOUSE | 2.37614763 | 0.0264551 |
| ACSF2_MOUSE | 6.85911933 | 0.04305497 |
| ACTN1_MOUSE | 2.94150451 | 0.00907073 |
| ADDG_MOUSE | 2.02630442 | 0.0064112 |
| AIFM1_MOUSE | 2.81649658 | 0.04296289 |
| AL1L1_MOUSE | 3.12190544 | 0.01219849 |
| AL3A2_MOUSE | 2.23955338 | 0.02893705 |
| ANK3_MOUSE | 2.44174507 | 0.02532274 |
| AP2A1_MOUSE | 1.65944151 | 0.02989611 |
| AP2B1_MOUSE | 2.39035809 | 0.02267046 |
| AP2M1_MOUSE | 2.50176909 | 0.04655494 |
| ASAH1_MOUSE* | 5.01610876 | 0.00167502 |
| ASTN1_MOUSE | 1.66472933 | 0.01603359 |
| AT1A1_MOUSE | 1.83121203 | 0.00714207 |
| AT1A2_MOUSE | 1.74425582 | 0.0026334 |
| AT1A3_MOUSE | 1.74595803 | 0.02985501 |
| AT1B1_MOUSE | 2.17994482 | 0.01875778 |
| AT1B2_MOUSE | 6.85159323 | 0.01769292 |

| | | |
|---|---|---|
| AT2B4_MOUSE | 1.73884867 | 0.01144252 |
| ATAD3_MOUSE | 1.77376102 | 0.03722607 |
| ATP5E_MOUSE | 2.93731529 | 0.03721463 |
| ATPG_MOUSE | 3.39113512 | 0.03156482 |
| ATPO_MOUSE | 6.13811283 | 0.04397278 |
| BAIP2_MOUSE | 2.59538287 | 0.04601941 |
| BASI_MOUSE | 2.69458393 | 0.0156804 |
| BRSK2_MOUSE | 1.84487865 | 0.01505495 |
| CA2D1_MOUSE | 3.16785152 | 0.01823684 |
| CAC1E_MOUSE | 2.08654395 | 0.00724175 |
| CADM2_MOUSE | 1.97486452 | 0.00348435 |
| CAPS1_MOUSE | 2.47981609 | 0.04828016 |
| CATD_MOUSE* | 8.6144266 | 0.00377278 |
| CBPE_MOUSE | 3.12398966 | 0.04667749 |
| CCG8_MOUSE | 3.17187285 | 0.02036904 |
| CCNY_MOUSE | 3.50274587 | 0.01081474 |
| CD47_MOUSE | 2.98286693 | 0.00840764 |
| CD81_MOUSE | 2.55345522 | 0.04181171 |
| CISD1_MOUSE | 1.91992184 | 0.0499723 |
| CISY_MOUSE | 2.97152825 | 0.02582269 |
| CKAP4_MOUSE* | 1.8302586 | 0.03220663 |
| CLH1_MOUSE | 2.80442701 | 0.02324201 |
| CLUS_MOUSE | 7.87342416 | 0.02781312 |
| CNDP2_MOUSE | 1.57796766 | 0.03215849 |
| COF1_MOUSE* | 3.70220246 | 0.02701788 |
| COTL1_MOUSE | 2.24339821 | 0.02674421 |
| COX2_MOUSE | 1.66755743 | 0.04724754 |
| COX41_MOUSE | 8.04690807 | 0.02848797 |
| CPNE1_MOUSE* | 3.37323728 | 0.04085177 |
| CRIP2_MOUSE | 6.82769925 | 0.04955228 |
| CSPG2_MOUSE | 6.24611017 | 0.04521241 |
| CTNA2_MOUSE | 1.87486094 | 0.04632975 |
| CXA1_MOUSE | 2.91852558 | 0.03654657 |
| CYFP1_MOUSE | 11.9840583 | 0.04398267 |
| DCE1_MOUSE | 2.84288313 | 0.04296491 |
| DCE2_MOUSE | 1.87714304 | 0.0499237 |
| DCLK1_MOUSE | 2.62481528 | 0.02118323 |
| DCTN1_MOUSE | 1.56010131 | 0.01730595 |
| DDB1_MOUSE | 2.72411734 | 0.00475071 |

| | | |
|---|---|---|
| DEST_MOUSE | 4.34731089 | 0.02196256 |
| DHB4_MOUSE | 1.69748231 | 0.04491371 |
| DIP2B_MOUSE | 1.96580728 | 0.03076635 |
| DMXL2_MOUSE | 1.63516189 | 0.03312546 |
| DNJC5_MOUSE | 5.11996889 | 0.01085887 |
| DPP6_MOUSE | 1.5301643 | 0.01851725 |
| DYL2_MOUSE | 4.19224278 | 0.02707414 |
| DYN1_MOUSE | 2.31587103 | 0.04280089 |
| EFTU_MOUSE | 1.52726819 | 0.00581057 |
| ENTP2_MOUSE | 2.19407876 | 0.00625933 |
| EP15R_MOUSE | 1.80511174 | 0.04214984 |
| EPHA4_MOUSE | 2460.83085 | 0.01654109 |
| ERP29_MOUSE | 17.3706319 | 0.02648587 |
| F10A1_MOUSE | 2.09881456 | 0.03346201 |
| FA49B_MOUSE | 2.07208701 | 0.0411204 |
| FAS_MOUSE | 2.07741976 | 0.01219864 |
| FBX41_MOUSE | 2.70989241 | 0.0363276 |
| FHL1_MOUSE* | 3.34333736 | 0.03506652 |
| FUMH_MOUSE | 2.42120508 | 0.04220973 |
| GABR1_MOUSE | 3.28428604 | 0.01480859 |
| GABR2_MOUSE | 3.43425887 | 0.03273567 |
| GBB2_MOUSE | 2.96271577 | 0.00257017 |
| GBRG2_MOUSE | 8.89414293 | 0.03526663 |
| GNA13_MOUSE | 2.49947339 | 0.00290725 |
| GNAI1_MOUSE | 3.3179847 | 0.02874637 |
| GNAI2_MOUSE | 4.67638206 | 0.03740986 |
| GNAO_MOUSE | 4.80553608 | 0.02535557 |
| GNAQ_MOUSE | 3.93664545 | 0.03285901 |
| GNAZ_MOUSE | 2.89572628 | 0.02645697 |
| GPC5B_MOUSE* | 4.4881951 | 0.02189772 |
| GPDA_MOUSE | 2.07136789 | 0.04772962 |
| GPM6A_MOUSE | 5.62523104 | 0.00678947 |
| GRIA1_MOUSE | 1.86606467 | 0.04587879 |
| GRIA2_MOUSE | 2.00334698 | 0.03965339 |
| GRP78_MOUSE | 1.71353245 | 0.00515339 |
| GTR1_MOUSE | 2.82450062 | 0.04301984 |
| HCN1_MOUSE | 1.84696478 | 0.03383597 |
| HEXB_MOUSE | 2.25030845 | 0.04017271 |
| HNRPU_MOUSE* | 3.08409018 | 0.03077163 |

| | | |
|---|---|---|
| HS12A_MOUSE | 2.02599041 | 0.04575905 |
| HSP74_MOUSE | 2.09076683 | 0.02485589 |
| HXK1_MOUSE | 1.86417568 | 0.03223978 |
| IMPA1_MOUSE | 3.86612134 | 0.0392026 |
| ITM2B_MOUSE* | 7.39999446 | 0.00713478 |
| ITM2C_MOUSE* | 4.82790884 | 0.0202228 |
| ITSN1_MOUSE | 2.93919762 | 0.02534814 |
| IVD_MOUSE | 1.62426359 | 0.0289076 |
| KCC2D_MOUSE | 1.56455433 | 0.01699244 |
| KCND2_MOUSE | 3.5143829 | 0.00905985 |
| KCRB_MOUSE | 1.81458902 | 0.02929205 |
| KI21A_MOUSE | 1.84691714 | 0.0114136 |
| KIF5C_MOUSE | 1.66905138 | 0.03508453 |
| KPYM_MOUSE* | 2.01532364 | 0.01438118 |
| KTN1_MOUSE | 2.7855293 | 0.01301305 |
| LAT1_MOUSE | 4.00246603 | 0.03615308 |
| LDHB_MOUSE | 1.92755339 | 0.04998723 |
| LETM1_MOUSE | 1.85597866 | 0.04277988 |
| LGI1_MOUSE | 2.04735999 | 0.01299686 |
| LGI2_MOUSE* | 4.74635604 | 0.04678316 |
| LIPA3_MOUSE | 1.68592935 | 0.0239067 |
| LIS1_MOUSE | 2.0801692 | 0.02626258 |
| LRRC7_MOUSE | 1.93307508 | 5.4308E-05 |
| M2OM_MOUSE | 3.10655303 | 0.03588682 |
| MADD_MOUSE | 2.7802968 | 0.02157297 |
| MAGI1_MOUSE* | 2.71135344 | 0.00303765 |
| MBLC2_MOUSE | 7.94560385 | 0.01830526 |
| MIA40_MOUSE | 8.28682583 | 0.04383079 |
| MPP2_MOUSE | 2.37879293 | 0.02489248 |
| MPP6_MOUSE | 1.96903366 | 0.00068579 |
| MRCKB_MOUSE | 3.37056361 | 0.00453073 |
| MYO6_MOUSE* | 7.65775067 | 0.04038318 |
| NAC1_MOUSE | 1.60781165 | 0.0205655 |
| NCALD_MOUSE | 45.0986498 | 0.03333189 |
| NCKP1_MOUSE | 1.76101097 | 0.03686968 |
| NDKA_MOUSE | 5.18466229 | 0.01994241 |
| NDUA6_MOUSE* | 3.22566946 | 0.04169493 |
| NDUS1_MOUSE | 1.85921323 | 0.00543366 |
| NF1_MOUSE | 4.40251115 | 0.02493713 |

| | | |
|---|---|---|
| NFASC_MOUSE | 1.52392687 | 0.03292294 |
| NNTM_MOUSE* | 5.15295413 | 0.00064755 |
| NRCAM_MOUSE | 1.71358651 | 0.01137874 |
| NRX3A_MOUSE | 2.86290478 | 0.02537298 |
| NSF_MOUSE | 1.79680977 | 0.01105155 |
| NSMA2_MOUSE* | 3.0656357 | 0.01513815 |
| NTRI_MOUSE | 4.06026601 | 0.03255085 |
| OAT_MOUSE | 2.38337287 | 0.04077862 |
| ODO1_MOUSE | 2.73375853 | 0.0474292 |
| ODPX_MOUSE | 1.55516198 | 0.01906577 |
| OPA1_MOUSE | 1.90523472 | 0.04247727 |
| OXR1_MOUSE | 1.65508292 | 0.00273805 |
| PALM2_MOUSE | 4.02528407 | 0.01746552 |
| PCCA_MOUSE | 1.52423112 | 0.03218611 |
| PDE2A_MOUSE | 1.70966649 | 0.01001083 |
| PDIA4_MOUSE | 3.02013745 | 0.0411904 |
| PDPR_MOUSE* | 2.17601378 | 0.00014154 |
| PGM1_MOUSE | 1.97753773 | 0.00126391 |
| PI4KA_MOUSE | 2.83055274 | 0.0147451 |
| PLCB1_MOUSE | 1.99490751 | 0.01982732 |
| PLPR4_MOUSE | 2.50426577 | 0.01188047 |
| PNPT1_MOUSE | 47.0149712 | 0.03136547 |
| PP1G_MOUSE* | 4.42574217 | 0.04759373 |
| PP2BA_MOUSE | 1.76925852 | 0.02415911 |
| PRDX4_MOUSE* | 4.45364429 | 0.0123701 |
| PRDX6_MOUSE | 1.6130104 | 0.0298685 |
| PRRT3_MOUSE | 2.28770195 | 0.02904083 |
| PTPRD_MOUSE | 2.86420019 | 0.03072229 |
| PTPRS_MOUSE | 2.25908928 | 0.02607941 |
| PYGB_MOUSE | 3.99445536 | 0.03208737 |
| R7BP_MOUSE* | 12.474933 | 0.04004716 |
| RAB1A_MOUSE | 3.9013582 | 0.04863898 |
| RAB3B_MOUSE | 11.741844 | 0.01679947 |
| RAB7A_MOUSE | 1.84884089 | 0.03543951 |
| RAN_MOUSE | 2.49778976 | 0.02187895 |
| RAP2B_MOUSE | 8.62180744 | 0.02144014 |
| RASH_MOUSE | 3.37978611 | 0.01763677 |
| RB3GP_MOUSE* | 4.54828995 | 0.01411416 |
| RBGPR_MOUSE | 3.97162065 | 0.03226824 |

| | | |
|---|---|---|
| RHOG_MOUSE | 1.93802777 | 0.02717621 |
| RL13_MOUSE* | 4.26738647 | 0.01404968 |
| RL22_MOUSE* | 9.99948728 | 0.03805332 |
| RL4_MOUSE* | 2.71230165 | 0.03870093 |
| RL5_MOUSE | 2.24191057 | 0.0344874 |
| RLA0_MOUSE* | 8.16503281 | 0.03251642 |
| RMD3_MOUSE | 38.0510859 | 0.04528451 |
| RPGF4_MOUSE | 2.68455757 | 0.02402321 |
| RPN1_MOUSE | 4.6181334 | 0.01553258 |
| S12A2_MOUSE | 2.59519204 | 0.03958399 |
| S39AC_MOUSE* | 2.75310155 | 0.04764758 |
| SAHH2_MOUSE | 2.71243878 | 0.01356875 |
| SATT_MOUSE | 8.42850683 | 0.04403561 |
| SC6A1_MOUSE | 3.41449618 | 0.04661652 |
| SCAM1_MOUSE | 3.27589443 | 0.03360363 |
| SCN1A_MOUSE | 6.14798148 | 0.01534375 |
| SCRB2_MOUSE* | 3.72143443 | 0.00760874 |
| SDHB_MOUSE | 2.84421004 | 0.02049879 |
| SEPT8_MOUSE | 1.78476244 | 0.0198599 |
| SFXN1_MOUSE | 7.93326104 | 0.04248025 |
| SFXN3_MOUSE | 2.37253319 | 0.01622603 |
| SND1_MOUSE | 2.10071498 | 0.03792139 |
| SPB6_MOUSE | 2.54675778 | 0.02905788 |
| SPRL1_MOUSE | 1.70498504 | 0.03331293 |
| SRBS2_MOUSE | 2.21686344 | 0.04168358 |
| SSDH_MOUSE | 1.66631376 | 0.00500585 |
| STXB1_MOUSE | 1.75202274 | 0.03577848 |
| SUCB2_MOUSE | 3.5185261 | 0.04360612 |
| SYIM_MOUSE | 1.85343949 | 0.03479621 |
| SYNJ1_MOUSE | 2.17892072 | 0.02888376 |
| SYNPR_MOUSE | 6.44464358 | 0.01783558 |
| SYT2_MOUSE | 2.48418822 | 0.04940637 |
| SYT7_MOUSE | 2.72888773 | 0.01288555 |
| SYUA_MOUSE* | 8.72755643 | 0.04972285 |
| TCPD_MOUSE* | 1.7520083 | 0.01304949 |
| TCPH_MOUSE | 2.26558686 | 0.02937783 |
| TERA_MOUSE | 2.18260205 | 0.02571782 |
| THIKA_MOUSE | 13.1940085 | 0.03926997 |
| THY1_MOUSE | 3.3568611 | 0.02204422 |

| | | |
|---|---|---|
| TIM13_MOUSE | 3.78323177 | 0.02091121 |
| TPP1_MOUSE* | 3.44625467 | 0.00473966 |
| TPPC3_MOUSE* | 2.46087779 | 0.01246953 |
| UBA1_MOUSE | 2.30346159 | 0.0187566 |
| UBE2O_MOUSE | 2.14510163 | 0.03154608 |
| UCRI_MOUSE | 4.83957182 | 0.0492247 |
| USMG5_MOUSE | 2.37515292 | 0.03253737 |
| VA0D1_MOUSE | 4.63836049 | 0.03338637 |
| VAMP1_MOUSE | 6.08333941 | 0.01721828 |
| VAMP7_MOUSE* | 15.422196 | 0.03328643 |
| VATE1_MOUSE | 2.81657735 | 0.03904979 |
| VCIP1_MOUSE | 4.69443448 | 0.03133417 |
| VDAC2_MOUSE | 2.92874791 | 0.0286335 |
| VGLU2_MOUSE | 1.96685604 | 0.00333991 |
| VISL1_MOUSE | 2.45048957 | 0.03958976 |
| VPS29_MOUSE | 25.4882128 | 0.04026089 |
| VPS35_MOUSE | 2.28353437 | 0.01015621 |
| VTA1_MOUSE | 39.3736025 | 0.04203121 |
| WFS1_MOUSE | 5.30342783 | 0.04325322 |
| XKR4_MOUSE* | 3.20447199 | 0.01798265 |

**Table S3. Formerly palmitoylated peptides identified for putative PPT1 substrates**. [x] Carbamidomethyl indicates location of palmitoylated cysteine (C) in peptide sequence.

| Protein Name (Uniprot ID) | Peptide Sequence | Modifications |
|---|---|---|
| ACON_MOUSE | CTTDHISAAGPWLK | [1] Carbamidomethyl |
| ACON_MOUSE | DVGGIVLANACGPCIGQWDRK | [11] Carbamidomethyl \|[14] Carbamidomethyl |
| ACON_MOUSE | VAVPSTIHCDHLIEAQVGGEK | [9] Carbamidomethyl |
| ACON_MOUSE | VGLIGSCTNSSYEDMGR | [7] Carbamidomethyl |
| ACTN1_MOUSE | DGLGFCALIHR | [6] Carbamidomethyl |
| ACTN1_MOUSE | EGLLLWCQR | [7] Carbamidomethyl |
| AL1L1_MOUSE | ECDVLPDDTVSTLYNR | [2] Carbamidomethyl |
| AL1L1_MOUSE | IAVIGQSLFGQEVYCQLR | [15] Carbamidomethyl |
| AL1L1_MOUSE | LRGEDGESECVINYVEK | [10] Carbamidomethyl |
| AL1L1_MOUSE | SPLIIFADCDLNK | [9] Carbamidomethyl |
| AL1L1_MOUSE | VPGAWTEACGQK | [9] Carbamidomethyl |
| ANK3_MOUSE | DIEVLEGKPIYVDCYGNLAPLTK | [14] Carbamidomethyl |
| AP2A1_MOUSE | AADLLYAMCDR | [9] Carbamidomethyl |
| AP2A1_MOUSE | ACNQLGQFLQHR | [2] Carbamidomethyl |
| AP2A1_MOUSE | ALQVGCLLR | [6] Carbamidomethyl |
| AP2A1_MOUSE | HLCELLAQQF | [3] Carbamidomethyl |
| AP2A1_MOUSE | QSAALCLLR | [6] Carbamidomethyl |
| AP2A1_MOUSE | TVFEALQAPACHENMVK | [11] Carbamidomethyl |
| AP2A1_MOUSE | YLALESMCTLASSEFSHEAVK | [8] Carbamidomethyl |
| AP2B1_MOUSE | DVSSLFPDVVNCMQTDNLELK | [12] Carbamidomethyl |
| AP2M1_MOUSE | IPTPLNTSGVQVICMK | [14] Carbamidomethyl |
| AP2M1_MOUSE | MCDVMAAYFGK | [2] Carbamidomethyl |
| AP2M1_MOUSE | QSIAIDDCTFHQCVR | [8] Carbamidomethyl \|[13] Carbamidomethyl |
| AP2M1_MOUSE | SYLSGMPECK | [9] Carbamidomethyl |

| | | |
|---|---|---|
| ASTN1_MOUSE | TLDSLQGCNEK | [8] Carbamidomethyl |
| AT1A1_MOUSE | LIIVEGCQR | [7] Carbamidomethyl |
| AT1A1_MOUSE | NIAFFSTNCVEGTAR | [9] Carbamidomethyl |
| AT1A1_MOUSE | NLEAVETLGSTSTICSDK | [15] Carbamidomethyl |
| AT1A2_MOUSE | CIELSCGSVR | [1] Carbamidomethyl \|[6] Carbamidomethyl |
| AT1A2_MOUSE | CSTILVQGK | [1] Carbamidomethyl |
| AT1A2_MOUSE | LCFVGLMSMIDPPR | [2] Carbamidomethyl |
| AT1A2_MOUSE | LIIVEGCQR | [7] Carbamidomethyl |
| AT1A2_MOUSE | NICFFSTNCVEGTAR | [3] Carbamidomethyl \|[9] Carbamidomethyl |
| AT1A2_MOUSE | NLEAVETLGSTSTICSDK | [15] Carbamidomethyl |
| AT1A2_MOUSE | VLGFCQLNLPSGKFPR | [5] Carbamidomethyl |
| AT1A3_MOUSE | ACVIHGTDLK | [2] Carbamidomethyl |
| AT1A3_MOUSE | CATILLQGK | [1] Carbamidomethyl |
| AT1A3_MOUSE | CIELSSGSVK | [1] Carbamidomethyl |
| AT1A3_MOUSE | KYNTDCVQGLTHSK | [6] Carbamidomethyl |
| AT1A3_MOUSE | LIIVEGCQR | [7] Carbamidomethyl |
| AT1A3_MOUSE | MSVEEVCRK | [7] Carbamidomethyl |
| AT1A3_MOUSE | NITFFSTNCVEGTAR | [9] Carbamidomethyl |
| AT1A3_MOUSE | NLEAVETLGSTSTICSDK | [15] Carbamidomethyl |
| AT1A3_MOUSE | SPDCTHDNPLETR | [4] Carbamidomethyl |
| AT1A3_MOUSE | SSHTWVALSHIAGLCNR | [15] Carbamidomethyl |
| AT1A3_MOUSE | VLGFCHYYLPEEQFPK | [5] Carbamidomethyl |
| AT1A3_MOUSE | YNTDCVQGLTHSK | [5] Carbamidomethyl |
| AT1B1_MOUSE | DDMIFEDCGNVPSEPK | [8] Carbamidomethyl |
| AT1B1_MOUSE | DSAQKDDMIFEDCGNVPSEPK | [13] Carbamidomethyl |
| AT1B1_MOUSE | YNPNVLPVQCTGK | [10] Carbamidomethyl |
| AT1B2_MOUSE | FLNVTPNVEVNVECR | [14] Carbamidomethyl |
| AT1B2_MOUSE | KSCGQVVEEWK | [3] Carbamidomethyl |

| | | |
|---|---|---|
| AT1B2_MOUSE | SCGQVVEEWK | [2] Carbamidomethyl |
| AT1B2_MOUSE | TQLGDCSGIGDPTHYGYSTGQPCVFIK | [6] Carbamidomethyl \|[23] Carbamidomethyl |
| AT2B4_MOUSE | GADAVAQISAHYGGVQEICTR | [19] Carbamidomethyl |
| AT2B4_MOUSE | HLDACETMGNATAICSDK | [5] Carbamidomethyl \|[15] Carbamidomethyl |
| AT2B4_MOUSE | TECGLLGFVTDLK | [3] Carbamidomethyl |
| ATPG_MOUSE | GLCGAIHSSVAK | [3] Carbamidomethyl |
| ATPO_MOUSE | GEVPCTVTTASPLDDAVLSELK | [5] Carbamidomethyl |
| CADM2_MOUSE | IIPSTPFPQEGQALTLTCESK | [18] Carbamidomethyl |
| CAPS1_MOUSE | HGMDEFISSNPCNFDHASLFEMVQR | [12] Carbamidomethyl |
| CAPS1_MOUSE | IVYCTMEVEGGEK | [4] Carbamidomethyl |
| CAPS1_MOUSE | KFEHQLLYNACQLDNPDEQAAQIR | [11] Carbamidomethyl |
| CAPS1_MOUSE | LCSMEMGQEHQYHSK | [2] Carbamidomethyl |
| CAPS1_MOUSE | LMASDMIESCVK | [10] Carbamidomethyl |
| CATD_MOUSE | AIGAVPLIQGEYMIPCEK | [16] Carbamidomethyl |
| CATD_MOUSE | GGCEAIVDTGTSLLVGPVEEVK | [3] Carbamidomethyl |
| CATD_MOUSE | ILDIACWVHHK | [6] Carbamidomethyl |
| CBPE_MOUSE | SGTAHEYSSCPDDAIFQSLAR | [10] Carbamidomethyl |
| CD81_MOUSE | NSLCPSGGNILTPLLQQDCHQK | [4] Carbamidomethyl \|[19] Carbamidomethyl |
| CD81_MOUSE | TFHETLNCCGSNALTTLTTTILR | [8] Carbamidomethyl \|[9] Carbamidomethyl |
| CISD1_MOUSE | KFPFCDGAHIK | [5] Carbamidomethyl |
| CISY_MOUSE | GYSIPECQK | [7] Carbamidomethyl |
| CLH1_MOUSE | CNEPAVWSQLAK | [1] Carbamidomethyl |
| CLH1_MOUSE | EDKLECSEELGDLVK | [6] Carbamidomethyl |
| CLH1_MOUSE | EVCFACVDGK | [3] Carbamidomethyl \|[6] Carbamidomethyl |
| CLH1_MOUSE | EVCFACVDGK | [3] Nethylmaleimide+water \|[6] Carbamidomethyl |
| CLH1_MOUSE | EVCFACVDGK | [3] Carbamidomethyl \|[6] Nethylmaleimide+water |
| CLH1_MOUSE | GQCDLELINVCNENSLFK | [3] Carbamidomethyl \|[11] Carbamidomethyl |
| CLH1_MOUSE | HSSLAGCQIINYR | [7] Carbamidomethyl |

| | | |
|---|---|---|
| CLH1_MOUSE | IHEGCEEPATHNALAK | [5] Carbamidomethyl |
| CLH1_MOUSE | LECSEELGDLVK | [3] Carbamidomethyl |
| CLH1_MOUSE | RDPHLACVAYER | [7] Carbamidomethyl |
| CLH1_MOUSE | VIQCFAETGQVQK | [4] Carbamidomethyl |
| CLH1_MOUSE | YESLELCRPVLQQGR | [7] Carbamidomethyl |
| CRIP2_MOUSE | ASSVTTFTGEPNMCPR | [14] Carbamidomethyl |
| CSPG2_MOUSE | FTFEEAEAECTSR | [10] Carbamidomethyl |
| CTNA2_MOUSE | LESIISGAALMADSSCTR | [16] Carbamidomethyl |
| CYFP1_MOUSE | DCPDNAEEYER | [2] Carbamidomethyl |
| DCE1_MOUSE | NLLSCENSDQGAR | [5] Carbamidomethyl |
| DCE2_MOUSE | LCALLYGDSGKPAEGGGSVTSR | [2] Carbamidomethyl |
| DCE2_MOUSE | MMGVPLQCSALLVR | [8] Carbamidomethyl |
| DCTN1_MOUSE | QSCTILISTMNK | [3] Carbamidomethyl |
| DMXL2_MOUSE | EIAALHEICNHESVIK | [9] Carbamidomethyl |
| DMXL2_MOUSE | VGCPVLALEVLSK | [3] Carbamidomethyl |
| DMXL2_MOUSE | VINLSQYGPACFGQEHAR | [11] Carbamidomethyl |
| DPP6_MOUSE | GENQGQTFTCGSALSPITDFK | [10] Carbamidomethyl |
| DYL2_MOUSE | KYNPTWHCIVGR | [8] Carbamidomethyl |
| DYL2_MOUSE | NADMSEDMQQDAVDCATQAMEK | [15] Carbamidomethyl |
| DYN1_MOUSE | ENCLILAVSPANSDLANSDALK | [3] Carbamidomethyl |
| DYN1_MOUSE | QLELACETQEEVDSWK | [6] Carbamidomethyl |
| ENTP2_MOUSE | AGQSLVECLEQALR | [8] Carbamidomethyl |
| FAS_MOUSE | ACVDTALENLSTLK | [2] Carbamidomethyl |
| FAS_MOUSE | FVFTPHMEAECLSESTALQK | [11] Carbamidomethyl |
| FAS_MOUSE | GYDYGPQFQGICEATLEGEQGK | [12] Carbamidomethyl |
| FAS_MOUSE | LGMLSPDGTCR | [10] Carbamidomethyl |
| FAS_MOUSE | SSCTIIPLMK | [3] Carbamidomethyl |
| GABR2_MOUSE | VFCCAFEESMFGSK | [3] Carbamidomethyl \|[4] Carbamidomethyl |

| | | |
|---|---|---|
| GBB2_MOUSE | ACGDSTLTQITAGLDPVGR | [2] Carbamidomethyl |
| GBB2_MOUSE | ELPGHTGYLSCCR | [11] Carbamidomethyl \|[12] Carbamidomethyl |
| GBB2_MOUSE | KACGDSTLTQITAGLDPVGR | [3] Carbamidomethyl |
| GBB2_MOUSE | TFVSGACDASIK | [7] Carbamidomethyl |
| GBB2_MOUSE | VSCLGVTDDGMAVATGSWDSFLK | [3] Carbamidomethyl |
| GBRG2_MOUSE | DCASFFCCFEDCR | [2] Carbamidomethyl \|[7] Carbamidomethyl \|[8] Carbamidomethyl \|[12] Carbamidomethyl |
| GNAI1_MOUSE | DSGVQACFNR | [7] Carbamidomethyl |
| GNAI1_MOUSE | EIYTHFTCATDTK | [8] Carbamidomethyl |
| GNAI1_MOUSE | IIHEAGYSEEECK | [12] Carbamidomethyl |
| GNAI1_MOUSE | LFDSICNNK | [6] Carbamidomethyl |
| GNAI2_MOUSE | EIYTHFTCATDTK | [8] Carbamidomethyl |
| GNAI2_MOUSE | IIHEDGYSEEECR | [12] Carbamidomethyl |
| GNAI2_MOUSE | LFDSICNNK | [6] Carbamidomethyl |
| GNAI2_MOUSE | LWADHGVQACFGR | [10] Carbamidomethyl |
| GNAI2_MOUSE | RLWADHGVQACFGR | [11] Carbamidomethyl |
| GNAO_MOUSE | LWGDSGIQECFNR | [10] Carbamidomethyl |
| GNAO_MOUSE | MHESLMLFDSICNNK | [12] Carbamidomethyl |
| GNAO_MOUSE | MVCDVVSR | [3] Carbamidomethyl |
| GNAQ_MOUSE | IIYSHFTCATDTENIR | [8] Carbamidomethyl |
| GNAQ_MOUSE | SLWNDPGIQECYDR | [11] Carbamidomethyl |
| GNAQ_MOUSE | TLESIMACCLSEEAK | [8] Carbamidomethyl \|[9] Carbamidomethyl |
| GNAZ_MOUSE | LWADPGAQACFGR | [10] Carbamidomethyl |
| GPM6A_MOUSE | ICTASENFLR | [2] Carbamidomethyl |
| GPM6A_MOUSE | KICTASENFLR | [3] Carbamidomethyl |
| GRIA1_MOUSE | GFCLIPQQSINEAIR | [3] Carbamidomethyl |
| GRIA1_MOUSE | RGNAGDCLANPAVPWGQGIDIQR | [7] Carbamidomethyl |
| GRIA2_MOUSE | RGNAGDCLANPAVPWGQGVEIER | [7] Carbamidomethyl |
| HS12A_MOUSE | EPECIHVMR | [4] Carbamidomethyl |

| | | |
|---|---|---|
| HSP74_MOUSE | GCALQCAILSPAFK | [2] Carbamidomethyl \|[6] Carbamidomethyl |
| HSP74_MOUSE | SVMDATQIAGLNCLR | [13] Carbamidomethyl |
| HXK1_MOUSE | AAQLCGAGMAAVVEK | [5] Carbamidomethyl |
| HXK1_MOUSE | AILQQLGLNSTCDDSILVK | [12] Carbamidomethyl |
| HXK1_MOUSE | ATDCVGHDVATLLR | [4] Carbamidomethyl |
| HXK1_MOUSE | KLPVGFTFSFPCR | [12] Carbamidomethyl |
| HXK1_MOUSE | MPLGFTFSFPCK | [11] Carbamidomethyl |
| IMPA1_MOUSE | YPCHSFIGEESVAAGEK | [3] Carbamidomethyl |
| KCRB_MOUSE | FCTGLTQIETLFK | [2] Carbamidomethyl |
| KI21A_MOUSE | TVNTEPEMMQCLK | [11] Carbamidomethyl |
| KIF5C_MOUSE | SLEPCDNTPIIDNITPVVDGISAEK | [5] Carbamidomethyl |
| LDHB_MOUSE | VIGSGCNLDSAR | [6] Carbamidomethyl |
| LETM1_MOUSE | GEEITKEEIDILSDACSK | [16] Carbamidomethyl |
| LGI1_MOUSE | DFDCIITEFAK | [4] Carbamidomethyl |
| LIS1_MOUSE | LLASCSADMTIK | [5] Carbamidomethyl |
| LIS1_MOUSE | MVRPNQDGTLIASCSNDQTVR | [14] Carbamidomethyl |
| LRRC7_MOUSE | SMCAPLPVAAQSTTLPSLSGR | [3] Carbamidomethyl |
| MIA40_MOUSE | GSDCIDQFR | [4] Carbamidomethyl |
| MPP2_MOUSE | DLELTPTSGTLCGSLSGK | [12] Carbamidomethyl |
| MPP2_MOUSE | RDLELTPTSGTLCGSLSGKK | [13] Carbamidomethyl |
| MPP2_MOUSE | VCVLDVNPQAVK | [2] Carbamidomethyl |
| MPP6_MOUSE | DWDNSGPFCGTISNK | [9] Carbamidomethyl |
| MPP6_MOUSE | TCILDVNPQALK | [2] Carbamidomethyl |
| NDUS1_MOUSE | DCFIVYQGHHGDVGAPMADVILPGAAYTEK | [2] Carbamidomethyl |
| NDUS1_MOUSE | DLLNKVDSDNLCTEEIFPTEGAGTDLR | [12] Carbamidomethyl |
| NDUS1_MOUSE | MCLVEIEK | [2] Carbamidomethyl |
| NDUS1_MOUSE | MLFLLGADGGCITR | [11] Carbamidomethyl |
| NDUS1_MOUSE | VDSDNLCTEEIFPTEGAGTDLR | [7] Carbamidomethyl |

| | | |
|---|---|---|
| NFASC_MOUSE | DNILIECEAK | [7] Carbamidomethyl |
| NFASC_MOUSE | DQGSYTCMASTELDQDLAK | [7] Carbamidomethyl |
| NFASC_MOUSE | ITNVSEEDSGEYFCLASNK | [14] Carbamidomethyl |
| NFASC_MOUSE | LDCPFFGSPIPTLR | [3] Carbamidomethyl |
| NFASC_MOUSE | SGGRPEEYEGEYQCFAR | [14] Carbamidomethyl |
| NFASC_MOUSE | TRLDCPFFGSPIPTLR | [5] Carbamidomethyl |
| NRCAM_MOUSE | AETYEGVYQCTAR | [10] Carbamidomethyl |
| NRCAM_MOUSE | DSTGTYTCVAR | [8] Carbamidomethyl |
| NRCAM_MOUSE | TLQITHVSEADSGNYQCIAK | [17] Carbamidomethyl |
| NSF_MOUSE | CPTDELSLSNCAVVNEK | [1] Carbamidomethyl \|[11] Carbamidomethyl |
| NSF_MOUSE | GILLYGPPGCGK | [10] Carbamidomethyl |
| NSF_MOUSE | SQLSCVVVDDIER | [5] Carbamidomethyl |
| NSF_MOUSE | THPSVVPGCIAFSLPQR | [9] Carbamidomethyl |
| NSF_MOUSE | VFPPEIVEQMGCK | [12] Carbamidomethyl |
| NTRI_MOUSE | EQSGEYECSASNDVAAPVVR | [8] Carbamidomethyl |
| NTRI_MOUSE | GTLQCEASAVPSAEFQWFK | [5] Carbamidomethyl |
| ODO1_MOUSE | AEQFYCGDTEGK | [6] Carbamidomethyl |
| ODO1_MOUSE | DVVVDLVCYR | [8] Carbamidomethyl |
| ODO1_MOUSE | ELEQIFCQFDSKLEAADEGSGDMK | [7] Carbamidomethyl |
| ODO1_MOUSE | FGLEGCEVLIPALK | [6] Carbamidomethyl |
| ODO1_MOUSE | ICEEAFTR | [2] Carbamidomethyl |
| ODO1_MOUSE | RFGLEGCEVLIPALK | [7] Carbamidomethyl |
| ODO1_MOUSE | SMTCPSTGLEEDVLFHIGK | [4] Carbamidomethyl |
| ODO1_MOUSE | SMTCPSTGLEEDVLFHIGK | [2] Oxidation (M)\|[4] Carbamidomethyl |
| ODO1_MOUSE | VVNAPIFHVNSDDPEAVMYVCK | [21] Carbamidomethyl |
| ODO1_MOUSE | YPNAELAWCQEEHK | [9] Carbamidomethyl |
| ODPX_MOUSE | DVSAPPPVSKPPAPTQPSPQPQIPCPAR | [25] Carbamidomethyl |
| ODPX_MOUSE | STVPHAYATADCDLGAVLK | [12] Carbamidomethyl |

| | | |
|---|---|---|
| OPA1_MOUSE | EGCTVSPETISLNVK | [3] Carbamidomethyl |
| PCCA_MOUSE | MADEAVCVGPAPTSK | [7] Carbamidomethyl |
| PDE2A_MOUSE | ATDQVVALACAFNK | [10] Carbamidomethyl |
| PDE2A_MOUSE | LVCEDPPHELPQEGK | [3] Carbamidomethyl |
| PI4KA_MOUSE | ICWQAAIFK | [2] Carbamidomethyl |
| PI4KA_MOUSE | QNTTLGATQLTERPACVK | [16] Carbamidomethyl |
| PLCB1_MOUSE | EVIEAIAECAFK | [9] Carbamidomethyl |
| PLCB1_MOUSE | LTDVAEECQNNQLK | [8] Carbamidomethyl |
| PLCB1_MOUSE | RVETALEACSLPSSR | [9] Carbamidomethyl |
| PLCB1_MOUSE | VVLPSLACLR | [8] Carbamidomethyl |
| PLPR4_MOUSE | NAEGSTVTCTGSIR | [9] Carbamidomethyl |
| PP2BA_MOUSE | FKEPPAYGPMCDILWSDPLEDFGNEK | [11] Carbamidomethyl |
| PRDX6_MOUSE | DFTPVCTTELGR | [6] Carbamidomethyl |
| PTPRD_MOUSE | TATMLCAASGNPDPEITWFK | [6] Carbamidomethyl |
| PTPRD_MOUSE | TPVDQTGVSGGVASFICQATGDPRPK | [17] Carbamidomethyl |
| PTPRD_MOUSE | VCLQPIR | [2] Carbamidomethyl |
| PTPRD_MOUSE | YECVATNSAGTR | [3] Carbamidomethyl |
| PTPRS_MOUSE | TATMLCAASGNPDPEITWFK | [6] Carbamidomethyl |
| PTPRS_MOUSE | VCLQPIR | [2] Carbamidomethyl |
| PYGB_MOUSE | TCAYTNHTVLPEALER | [2] Carbamidomethyl |
| PYGB_MOUSE | TCFETFPDK | [2] Carbamidomethyl |
| RAP2B_MOUSE | ALAEEWSCPFMETSAK | [8] Carbamidomethyl |
| RASH_MOUSE | LNPPDESGPGCMSCK | [11] Carbamidomethyl \|[14] Carbamidomethyl |
| RHOG_MOUSE | YLECSALQQDGVK | [4] Carbamidomethyl |
| S39AC_MOUSE | YFGTSSSQCMETK | [9] Carbamidomethyl |
| SAHH2_MOUSE | LCVPAMNVNDSVTK | [2] Carbamidomethyl |
| SC6A1_MOUSE | QCDNPWNTDR | [2] Carbamidomethyl |
| SDHB_MOUSE | CHTIMNCTQTCPK | [1] Carbamidomethyl \|[7] Carbamidomethyl \|[11] Carbamidomethyl |

| | | |
|---|---|---|
| SEPT8_MOUSE | ELEEETNAFNCR | [11] Carbamidomethyl |
| SEPT8_MOUSE | QYPWGVVQVENENHCDFVK | [15] Carbamidomethyl |
| SEPT8_MOUSE | RRELEEETNAFNCR | [13] Carbamidomethyl |
| SEPT8_MOUSE | STLMNTLFNTTFETEEASHHEECVR | [23] Carbamidomethyl |
| SRBS2_MOUSE | DLMNSEVICSVK | [9] Carbamidomethyl |
| SSDH_MOUSE | EVGEVLCTDPLVSK | [7] Carbamidomethyl |
| SSDH_MOUSE | LGTVADCGVPEAR | [7] Carbamidomethyl |
| STXB1_MOUSE | AAHVFFTDSCPDALFNELVK | [10] Carbamidomethyl |
| STXB1_MOUSE | LAEQIATLCATLK | [9] Carbamidomethyl |
| STXB1_MOUSE | YSTHLHLAEDCMK | [11] Carbamidomethyl |
| SYIM_MOUSE | VHFVPGWDCHGLPIETK | [9] Carbamidomethyl |
| SYNJ1_MOUSE | TSPCQSPTVPEYSAPSLPIRPSR | [4] Carbamidomethyl |
| SYNPR_MOUSE | EVLLLMSACK | [9] Carbamidomethyl |
| SYNPR_MOUSE | LQQVTFEVPTCEGK | [11] Carbamidomethyl |
| SYT2_MOUSE | LGDICTSLR | [5] Carbamidomethyl |
| SYT2_MOUSE | LTVCILEAK | [4] Carbamidomethyl |
| TERA_MOUSE | AIANECQANFISIK | [6] Carbamidomethyl |
| TERA_MOUSE | LGDVISIQPCPDVK | [10] Carbamidomethyl |
| TERA_MOUSE | QAAPCVLFFDELDSIAK | [5] Carbamidomethyl |
| THY1_MOUSE | VTSLTACLVNQNLR | [7] Carbamidomethyl |
| UBA1_MOUSE | DNPGVVTCLDEAR | [8] Carbamidomethyl |
| VA0D1_MOUSE | NIVWIAECIAQR | [8] Carbamidomethyl |
| VDAC2_MOUSE | SCSGVEFSTSGSSNTDTGK | [2] Carbamidomethyl |
| VDAC2_MOUSE | WCEYGLTFTEK | [2] Carbamidomethyl |
| VGLU2_MOUSE | ILQGLVEGVTYPACHGIWSK | [14] Carbamidomethyl |

**Table S4. Residual PPT1 substrates**. Proteins identified in the secondary validation screen that were not identified as putative substrates in the primary screen.

| Uniprot ID | Gene Name |
|---|---|
| 1433E_MOUSE | *Ywhae* |
| 1433F_MOUSE | *Ywhah* |
| 1433G_MOUSE | *Ywhag* |
| 1433S_MOUSE | *Sfn* |
| 1433T_MOUSE | *Ywhaq* |
| 1433Z_MOUSE | *Ywhaz* |
| 2A5E_MOUSE | *Ppp2r5e* |
| 2AAA_MOUSE | *Ppp2r1a* |
| 2AAB_MOUSE | *Ppp2r1b* |
| 2ABA_MOUSE | *Ppp2r2a* |
| 2ABB_MOUSE | *Ppp2r2b* |
| 2ABD_MOUSE | *Ppp2r2d* |
| 2ABG_MOUSE | *Ppp2r2c* |
| 3HIDH_MOUSE | *Hibadh* |
| 4F2_MOUSE | *Slc3a2* |
| A2MG_MOUSE | *A2m* |
| A4_MOUSE | *App* |
| AAK1_MOUSE | *Aak1* |
| AATC_MOUSE | *Got1* |
| AATM_MOUSE | *Got2* |
| ACAD9_MOUSE | *Acad9* |
| ACADL_MOUSE | *Acadl* |
| ACADM_MOUSE | *Acadm* |
| ACADV_MOUSE | *Acadvl* |
| ACLY_MOUSE | *Acly* |
| ACO10_MOUSE | *Acot10* |
| ACO13_MOUSE | *Acot13* |
| ACOT9_MOUSE | *Acot9* |
| ACPM_MOUSE | *Ndufab1* |
| ACSL6_MOUSE | *Acsl6* |
| ACTA_MOUSE | *Acta2* |
| ACTB_MOUSE | *Actb* |
| ACTC_MOUSE | *Actc1* |
| ACTN2_MOUSE | *Actn2* |
| ACTN4_MOUSE | *Actn4* |
| ACYP1_MOUSE | *Acyp1* |

| | |
|---|---|
| ADA22_MOUSE | *Adam22* |
| ADA23_MOUSE | *Adam23* |
| ADDA_MOUSE | *Add1* |
| ADDB_MOUSE | *Add2* |
| ADT1_MOUSE | *Slc25a4* |
| ADT2_MOUSE | *Slc25a5* |
| AFG2H_MOUSE | *Spata5* |
| AFG32_MOUSE | *Afg3l2* |
| AGFG1_MOUSE | *Agfg1* |
| AGK_MOUSE | *Agk* |
| AHSA1_MOUSE | *Ahsa1* |
| AINX_MOUSE | *Ina* |
| AK1A1_MOUSE | *Akr1a1* |
| AKAP5_MOUSE | *Akap5* |
| AL1B1_MOUSE | *Aldh1b1* |
| AL7A1_MOUSE | *Aldh7a1* |
| ALBU_MOUSE | *Alb* |
| ALDH2_MOUSE | *Aldh2* |
| ALDOA_MOUSE | *Aldoa* |
| ALDOC_MOUSE | *Aldoc* |
| AMPH_MOUSE | *Amph* |
| AMPL_MOUSE | *Lap3* |
| AMRP_MOUSE | *Lrpap1* |
| ANK2_MOUSE | *Ank2* |
| ANS1B_MOUSE | *Anks1b* |
| ANXA5_MOUSE | *Anxa5* |
| ANXA6_MOUSE | *Anxa6* |
| ANXA7_MOUSE | *Anxa7* |
| AOFB_MOUSE | *Maob* |
| AP180_MOUSE | *Snap91* |
| AP1B1_MOUSE | *Ap1b1* |
| AP2A2_MOUSE | *Ap2a2* |
| AP2S1_MOUSE | *Ap2s1* |
| AP3B2_MOUSE | *Ap3b2* |
| AP3D1_MOUSE | *Ap3d1* |
| AP3M2_MOUSE | *Ap3m2* |
| APOE_MOUSE | *Apoe* |
| ARC1A_MOUSE | *Arpc1a* |
| ARF1_MOUSE | *Arf1* |

| | |
|---|---|
| ARF2_MOUSE | *Arf2* |
| ARF5_MOUSE | *Arf5* |
| ARFG1_MOUSE | *Arfgap1* |
| ARM10_MOUSE | *Armc10* |
| ARMC1_MOUSE | *Armc1* |
| ARP2_MOUSE | *Actr2* |
| ARP3_MOUSE | *Actr3* |
| ARP3B_MOUSE | *Actr3b* |
| ARP5L_MOUSE | *Arpc5l* |
| ARPC2_MOUSE | *Arpc2* |
| ARPC3_MOUSE | *Arpc3* |
| ARPC4_MOUSE | *Arpc4* |
| ARPC5_MOUSE | *Arpc5* |
| ARSB_MOUSE | *Arsb* |
| ASGL1_MOUSE | *Asrgl1* |
| ASNA_MOUSE | *Asna1* |
| ASSY_MOUSE | *Ass1* |
| AT2A1_MOUSE | *Atp2a1* |
| AT2A2_MOUSE | *Atp2a2* |
| AT2B1_MOUSE | *Atp2b1* |
| AT2B2_MOUSE | *Atp2b2* |
| AT5F1_MOUSE | *Atp5pb* |
| AT8A1_MOUSE | *Atp8a1* |
| ATIF1_MOUSE | *ATP5IF1* |
| ATLA1_MOUSE | *Atl1* |
| ATP5H_MOUSE | *Atp5pd* |
| ATP5L_MOUSE | *Atp5mg* |
| ATPA_MOUSE | *Atp5f1a* |
| ATPB_MOUSE | *Atp5f1b* |
| ATPD_MOUSE | *Atp5f1d* |
| ATPK_MOUSE | *Atp5mf* |
| ATPMD_MOUSE | *Atp5md* |
| AUHM_MOUSE | *Auh* |
| AUXI_MOUSE | *Dnajc6* |
| B2L13_MOUSE | *Bcl2l13* |
| BACH_MOUSE | *Acot7* |
| BASP1_MOUSE | *Basp1* |
| BCAS1_MOUSE | *Bcas1* |
| BCS1_MOUSE | *Bcs1l* |

| | |
|---|---|
| BDH_MOUSE | *Bdh1* |
| BIN1_MOUSE | *Bin1* |
| BIN2_MOUSE | *Bin2* |
| BIP_MOUSE | *Hspa5* |
| BORG4_MOUSE | *Cdc42ep4* |
| BPHL_MOUSE | *Bphl* |
| BSN_MOUSE | *Bsn* |
| C1QBP_MOUSE | *C1qbp* |
| C1TM_MOUSE | *Mthfd1l* |
| C2C2L_MOUSE | *C2cd2l* |
| CAD13_MOUSE | *Cdh13* |
| CADH2_MOUSE | *Cdh2* |
| CADM1_MOUSE | *Cadm1* |
| CADM3_MOUSE | *Cadm3* |
| CADM4_MOUSE | *Cadm4* |
| CAH2_MOUSE | *Ca2* |
| CALB1_MOUSE | *Calb1* |
| CALB2_MOUSE | *Calb2* |
| CALL3_MOUSE | *Calml3* |
| CALM1_MOUSE | *Calm1* |
| CALR_MOUSE | *Calr* |
| CALU_MOUSE | *Calu* |
| CALX_MOUSE | *Canx* |
| CAMKV_MOUSE | *Camkv* |
| CANB1_MOUSE | *Ppp3r1* |
| CAND1_MOUSE | *Cand1* |
| CAP1_MOUSE | *Cap1* |
| CAP2_MOUSE | *Cap2* |
| CAPZB_MOUSE | *Capzb* |
| CATB_MOUSE | *Ctsb* |
| CAZA1_MOUSE | *Capza1* |
| CAZA2_MOUSE | *Capza2* |
| CBR1_MOUSE | *Cbr1* |
| CC136_MOUSE | *Ccdc136* |
| CC50A_MOUSE | *Tmem30a* |
| CC50B_MOUSE | *Tmem30b* |
| CD166_MOUSE | *Alcam* |
| CDC37_MOUSE | *Cdc37* |
| CDC42_MOUSE | *Cdc42* |

| | |
|---|---|
| CEND_MOUSE | *Cend1* |
| CH10_MOUSE | *Hspe1* |
| CH60_MOUSE | *Hspd1* |
| CHP1_MOUSE | *Chp1* |
| CLAP2_MOUSE | *Clasp2* |
| CLCA_MOUSE | *Clta* |
| CLCB_MOUSE | *Cltb* |
| CLD11_MOUSE | *Cldn11* |
| CLIC4_MOUSE | *Clic4* |
| CMC1_MOUSE | *Slc25a12* |
| CMTD1_MOUSE | *Comtd1* |
| CN37_MOUSE | *Cnp* |
| CNRP1_MOUSE | *Cnrip1* |
| CNTN1_MOUSE | *Cntn1* |
| CNTN2_MOUSE | *Cntn2* |
| CNTP2_MOUSE | *Cntnap2* |
| CO1A1_MOUSE | *Col1a1* |
| CO3_MOUSE | *C3* |
| CO4B_MOUSE | *C4b* |
| COF1_MOUSE | *Cfl1* |
| COQ9_MOUSE | *Coq9* |
| COR1A_MOUSE | *Coro1a* |
| COR1B_MOUSE | *Coro1b* |
| COR1C_MOUSE | *Coro1c* |
| COX5A_MOUSE | *Cox5a* |
| COX5B_MOUSE | *Cox5b* |
| CPLX1_MOUSE | *Cplx1* |
| CPLX2_MOUSE | *Cplx2* |
| CPLX3_MOUSE | *Cplx3* |
| CPNE6_MOUSE | *Cpne6* |
| CRK_MOUSE | *Crk* |
| CRKL_MOUSE | *Crkl* |
| CRYAB_MOUSE | *Cryab* |
| CRYM_MOUSE | *Crym* |
| CSK21_MOUSE | *Csnk2a1* |
| CSK2B_MOUSE | *Csnk2b* |
| CSKI1_MOUSE | *Caskin1* |
| CSN1_MOUSE | *Gps1* |
| CSN2_MOUSE | *Cops2* |

| | |
|---|---|
| CSN4_MOUSE | *Cops4* |
| CSPG5_MOUSE | *Cspg5* |
| CSRP1_MOUSE | *Csrp1* |
| CTBP1_MOUSE | *Ctbp1* |
| CTNB1_MOUSE | *Ctnnb1* |
| CTND2_MOUSE | *Ctnnd2* |
| CTTB2_MOUSE | *Cttnbp2* |
| CX6B1_MOUSE | *Cox6b1* |
| CY1_MOUSE | *Cyc1* |
| CYC_MOUSE | *Cycs* |
| CYC2_MOUSE | *Cyct* |
| CYFP2_MOUSE | *Cyfip2* |
| DBNL_MOUSE | *Dbnl* |
| DC1I1_MOUSE | *Dync1i1* |
| DC1I2_MOUSE | *Dync1i2* |
| DC1L1_MOUSE | *Dync1li1* |
| DCTN2_MOUSE | *Dctn2* |
| DDAH1_MOUSE | *Ddah1* |
| DECR_MOUSE | *Decr1* |
| DEMA_MOUSE | *Dmtn* |
| DESM_MOUSE | *Des* |
| DHE3_MOUSE | *Glud1* |
| DHPR_MOUSE | *Qdpr* |
| DIRA2_MOUSE | *Diras2* |
| DLDH_MOUSE | *Dld* |
| DLG1_MOUSE | *Dlg1* |
| DLG2_MOUSE | *Dlg2* |
| DLG3_MOUSE | *Dlg3* |
| DLG4_MOUSE | *Dlg4* |
| DLGP4_MOUSE | *Dlgap4* |
| DMXL1_MOUSE | *Dmxl1* |
| DNM1L_MOUSE | *Dnm1l* |
| DOC2B_MOUSE | *Doc2b* |
| DOPD_MOUSE | *Ddt* |
| DPP10_MOUSE | *Dpp10* |
| DPYL1_MOUSE | *Crmp1* |
| DPYL2_MOUSE | *Dpysl2* |
| DPYL3_MOUSE | *Dpysl3* |
| DPYL4_MOUSE | *Dpysl4* |

| | |
|---|---|
| DPYL5_MOUSE | *Dpysl5* |
| DREB_MOUSE | *Dbn1* |
| DUS3_MOUSE | *Dusp3* |
| DYHC1_MOUSE | *Dync1h1* |
| DYN3_MOUSE | *Dnm3* |
| E41L1_MOUSE | *Epb41l1* |
| E41L2_MOUSE | *Epb41l2* |
| E41L3_MOUSE | *Epb41l3* |
| EAA1_MOUSE | *Slc1a3* |
| EAA2_MOUSE | *Slc1a2* |
| ECHA_MOUSE | *Hadha* |
| ECHB_MOUSE | *Hadhb* |
| ECHM_MOUSE | *Echs1* |
| EF1A1_MOUSE | *Eef1a1* |
| EF1A2_MOUSE | *Eef1a2* |
| EF1G_MOUSE | *Eef1g* |
| EF2_MOUSE | *Eef2* |
| EFHD1_MOUSE | *Efhd1* |
| EFHD2_MOUSE | *Efhd2* |
| EHD1_MOUSE | *Ehd1* |
| EHD3_MOUSE | *Ehd3* |
| ELMO2_MOUSE | *Elmo2* |
| ENOA_MOUSE | *Eno1* |
| ENOG_MOUSE | *Eno2* |
| ENPL_MOUSE | *Hsp90b1* |
| EPN1_MOUSE | *Epn1* |
| ERC2_MOUSE | *Erc2* |
| ERMIN_MOUSE | *Ermn* |
| ERP44_MOUSE | *Erp44* |
| ETFA_MOUSE | *Etfa* |
| ETFB_MOUSE | *Etfb* |
| ETFD_MOUSE | *Etfdh* |
| EXOG_MOUSE | *Exog* |
| EZRI_MOUSE | *Ezr* |
| FABP5_MOUSE | *Fabp5* |
| FABPH_MOUSE | *Fabp3* |
| FAHD2_MOUSE | *Fahd2* |
| FKB1A_MOUSE | *Fkbp1a* |
| FKBP2_MOUSE | *Fkbp2* |

| | |
|---|---|
| FLOT1_MOUSE | *Flot1* |
| FLOT2_MOUSE | *Flot2* |
| FRIH_MOUSE | *Fth1* |
| FSCN1_MOUSE | *Fscn1* |
| G3P_MOUSE | *Gapdh* |
| G3PT_MOUSE | *Gapdhs* |
| G6PI_MOUSE | *Gpi* |
| GABT_MOUSE | *Abat* |
| GAK_MOUSE | *Gak* |
| GAL3A_MOUSE | *Gatd3a* |
| GANAB_MOUSE | *Ganab* |
| GARS_MOUSE | *Gars1* |
| GBB1_MOUSE | *Gnb1* |
| GBB3_MOUSE | *Gnb3* |
| GBRA1_MOUSE | *Gabra1* |
| GD1L1_MOUSE | *Gdap1l1* |
| GDAP1_MOUSE | *Gdap1* |
| GDIA_MOUSE | *Gdi1* |
| GDIB_MOUSE | *Gdi2* |
| GDIR1_MOUSE | *Arhgdia* |
| GELS_MOUSE | *Gsn* |
| GEPH_MOUSE | *Gphn* |
| GFAP_MOUSE | *Gfap* |
| GGT7_MOUSE | *Ggt7* |
| GHC1_MOUSE | *Slc25a22* |
| GHC2_MOUSE | *Slc25a18* |
| GIT1_MOUSE | *Git1* |
| GLNA_MOUSE | *Glul* |
| GLO2_MOUSE | *Hagh* |
| GLOD4_MOUSE | *Glod4* |
| GLPK_MOUSE | *Gk* |
| GLPK2_MOUSE | *Gk2* |
| GLRX5_MOUSE | *Glrx5* |
| GLSK_MOUSE | *Gls* |
| GLU2B_MOUSE | *Prkcsh* |
| GMFB_MOUSE | *Gmfb* |
| GMFG_MOUSE | *Gmfg* |
| GNA11_MOUSE | *Gna11* |
| GNA12_MOUSE | *Gna12* |

| | |
|---|---|
| GNAS1_MOUSE | *Gnas* |
| GNB5_MOUSE | *Gnb5* |
| GP158_MOUSE | *Gpr158* |
| GPC1_MOUSE | *Gpc1* |
| GPD1L_MOUSE | *Gpd1l* |
| GPDM_MOUSE | *Gpd2* |
| GPM6B_MOUSE | *Gpm6b* |
| GRB2_MOUSE | *Grb2* |
| GRIN1_MOUSE | *Gprin1* |
| GRM3_MOUSE | *Grm3* |
| GRP75_MOUSE | *Hspa9* |
| GRPE1_MOUSE | *Grpel1* |
| GSTM1_MOUSE | *Gstm1* |
| GSTM3_MOUSE | *Gstm3* |
| GSTM5_MOUSE | *Gstm5* |
| GSTM6_MOUSE | *Gstm6* |
| GSTP1_MOUSE | *Gstp1* |
| GSTP2_MOUSE | *Gstp2* |
| GTR3_MOUSE | *Slc2a3* |
| GUAD_MOUSE | *Gda* |
| H4_MOUSE | *H4c1* |
| HBA_MOUSE | *Hba* |
| HBAZ_MOUSE | *Hbz* |
| HBB1_MOUSE | *Hbb-b1* |
| HCD2_MOUSE | *Hsd17b10* |
| HCDH_MOUSE | *Hadh* |
| HDHD2_MOUSE | *Hdhd2* |
| HECAM_MOUSE | *Hepacam* |
| HEMH_MOUSE | *Fech* |
| HGS_MOUSE | *Hgs* |
| HIBCH_MOUSE | *Hibch* |
| HINT2_MOUSE | *Hint2* |
| HMGCL_MOUSE | *Hmgcl* |
| HMOX2_MOUSE | *Hmox2* |
| HNRPK_MOUSE | *Hnrnpk* |
| HOME1_MOUSE | *Homer1* |
| HOME3_MOUSE | *Homer3* |
| HPCA_MOUSE | *Hpca* |
| HPCL4_MOUSE | *Hpcal4* |

| | |
|---|---|
| HPRT_MOUSE | *Hprt1* |
| HS105_MOUSE | *Hsph1* |
| HS12B_MOUSE | *Hspa12b* |
| HS71A_MOUSE | *Hspa1a* |
| HS74L_MOUSE | *Hspa4l* |
| HS90A_MOUSE | *Hsp90aa1* |
| HS90B_MOUSE | *Hsp90ab1* |
| HSP72_MOUSE | *Hspa2* |
| HSP7C_MOUSE | *Hspa8* |
| ICAM5_MOUSE | *Icam5* |
| IDH3A_MOUSE | *Idh3a* |
| IDHC_MOUSE | *Idh1* |
| IDHG1_MOUSE | *Idh3g* |
| IDHP_MOUSE | *Idh2* |
| IF4B_MOUSE | *Eif4b* |
| IF4H_MOUSE | *Eif4h* |
| IF5A1_MOUSE | *Eif5a* |
| IF5A2_MOUSE | *Eif5a2* |
| IGSF8_MOUSE | *Igsf8* |
| IMB1_MOUSE | *Kpnb1* |
| IPO5_MOUSE | *Ipo5* |
| IPYR_MOUSE | *Ppa1* |
| IPYR2_MOUSE | *Ppa2* |
| IQEC1_MOUSE | *Iqsec1* |
| IQEC2_MOUSE | *Iqsec2* |
| ITIH2_MOUSE | *Itih2* |
| K0513_MOUSE | *Kiaa0513* |
| KAD1_MOUSE | *Ak1* |
| KAD2_MOUSE | *Ak2* |
| KAD3_MOUSE | *Ak3* |
| KAD4_MOUSE | *Ak4* |
| KAP2_MOUSE | *Prkar2a* |
| KAP3_MOUSE | *Prkar2b* |
| KCAB2_MOUSE | *Kcnab2* |
| KCC2A_MOUSE | *Camk2a* |
| KCC2B_MOUSE | *Camk2b* |
| KCC2G_MOUSE | *Camk2g* |
| KCRS_MOUSE | *Ckmt2* |
| KCRU_MOUSE | *Ckmt1* |

| | |
|---|---|
| KCY_MOUSE | *Cmpk1* |
| KIF2A_MOUSE | *Kif2a* |
| KLC1_MOUSE | *Klc1* |
| KLC2_MOUSE | *Klc2* |
| KPCB_MOUSE | *Prkcb* |
| KPCG_MOUSE | *Prkcg* |
| KPYM_MOUSE | *Pkm* |
| KPYR_MOUSE | *Pklr* |
| L1CAM_MOUSE | *L1cam* |
| LACTB_MOUSE | *Lactb* |
| LANC2_MOUSE | *Lancl2* |
| LASP1_MOUSE | *Lasp1* |
| LDHA_MOUSE | *Ldha* |
| LDHC_MOUSE | *Ldhc* |
| LGUL_MOUSE | *Glo1* |
| LIGO1_MOUSE | *Lingo1* |
| LIN7A_MOUSE | *Lin7a* |
| LIPA2_MOUSE | *Ppfia2* |
| LNEBL_MOUSE | *Nebl* |
| LONM_MOUSE | *Lonp1* |
| LPPRC_MOUSE | *Lrpprc* |
| LRC59_MOUSE | *Lrrc59* |
| LSAMP_MOUSE | *Lsamp* |
| LXN_MOUSE | *Lxn* |
| LY6H_MOUSE | *Ly6h* |
| LYAG_MOUSE | *Gaa* |
| LYRIC_MOUSE | *Mtdh* |
| MAG_MOUSE | *Mag* |
| MAOM_MOUSE | *Me2* |
| MAON_MOUSE | *Me3* |
| MAP1A_MOUSE | *Map1a* |
| MAP1B_MOUSE | *Map1b* |
| MAP4_MOUSE | *Map4* |
| MAP6_MOUSE | *Map6* |
| MARC2_MOUSE | *Marc2* |
| MARCS_MOUSE | *Marcks* |
| MARE1_MOUSE | *Mapre1* |
| MARE2_MOUSE | *Mapre2* |
| MARE3_MOUSE | *Mapre3* |

| | |
|---|---|
| MBP_MOUSE | *Mbp* |
| MCU_MOUSE | *Mcu* |
| MDHC_MOUSE | *Mdh1* |
| MDHM_MOUSE | *Mdh2* |
| MFF_MOUSE | *Mff* |
| MFN2_MOUSE | *Mfn2* |
| MFR1L_MOUSE | *Mtfr1l* |
| MGLL_MOUSE | *Mgll* |
| MIC13_MOUSE | *Micos13* |
| MIC19_MOUSE | *Chchd3* |
| MIC25_MOUSE | *Chchd6* |
| MIC26_MOUSE | *Apoo* |
| MIC60_MOUSE | *Immt* |
| MIF_MOUSE | *Mif* |
| MK01_MOUSE | *Mapk1* |
| MK03_MOUSE | *Mapk3* |
| MK04_MOUSE | *Mapk4* |
| ML12B_MOUSE | *Myl12b* |
| MMSA_MOUSE | *Aldh6a1* |
| MOG_MOUSE | *Mog* |
| MP2K1_MOUSE | *Map2k1* |
| MPCP_MOUSE | *Slc25a3* |
| MPPA_MOUSE | *Pmpca* |
| MTAP2_MOUSE | *Map2* |
| MTCH1_MOUSE | *Mtch1* |
| MTCH2_MOUSE | *Mtch2* |
| MTCL1_MOUSE | *Mtcl1* |
| MTX1_MOUSE | *Mtx1* |
| MTX2_MOUSE | *Mtx2* |
| MUTA_MOUSE | *Mmut* |
| MY18A_MOUSE | *Myo18a* |
| MYH10_MOUSE | *Myh10* |
| MYL6_MOUSE | *Myl6* |
| MYL6B_MOUSE | *Myl6b* |
| MYL9_MOUSE | *Myl9* |
| MYO5A_MOUSE | *Myo5a* |
| MYO5B_MOUSE | *Myo5b* |
| MYPR_MOUSE | *Plp1* |
| NAC2_MOUSE | *Slc8a2* |

| | |
|---|---|
| NB5R1_MOUSE | *Cyb5r1* |
| NB5R3_MOUSE | *Cyb5r3* |
| NCAM1_MOUSE | *Ncam1* |
| NCAM2_MOUSE | *Ncam2* |
| NCAN_MOUSE | *Ncan* |
| NCDN_MOUSE | *Ncdn* |
| NCEH1_MOUSE | *Nceh1* |
| NCKPL_MOUSE | *Nckap1l* |
| NCS1_MOUSE | *Ncs1* |
| NDKB_MOUSE | *Nme2* |
| NDRG1_MOUSE | *Ndrg1* |
| NDRG2_MOUSE | *Ndrg2* |
| NDRG3_MOUSE | *Ndrg3* |
| NDUA2_MOUSE | *Ndufa2* |
| NDUA4_MOUSE | *Ndufa4* |
| NDUA5_MOUSE | *Ndufa5* |
| NDUA7_MOUSE | *Ndufa7* |
| NDUA8_MOUSE | *Ndufa8* |
| NDUA9_MOUSE | *Ndufa9* |
| NDUAA_MOUSE | *Ndufa10* |
| NDUAC_MOUSE | *Ndufa12* |
| NDUAD_MOUSE | *Ndufa13* |
| NDUB3_MOUSE | *Ndufb3* |
| NDUB4_MOUSE | *Ndufb4* |
| NDUB5_MOUSE | *Ndufb5* |
| NDUB6_MOUSE | *Ndufb6* |
| NDUB7_MOUSE | *Ndufb7* |
| NDUB8_MOUSE | *Ndufb8* |
| NDUB9_MOUSE | *Ndufb9* |
| NDUBA_MOUSE | *Ndufb10* |
| NDUBB_MOUSE | *Ndufb11* |
| NDUC2_MOUSE | *Ndufc2* |
| NDUF2_MOUSE | *Ndufaf2* |
| NDUS2_MOUSE | *Ndufs2* |
| NDUS3_MOUSE | *Ndufs3* |
| NDUS4_MOUSE | *Ndufs4* |
| NDUS5_MOUSE | *Ndufs5* |
| NDUS6_MOUSE | *Ndufs6* |
| NDUS7_MOUSE | *Ndufs7* |

| | |
|---|---|
| NDUS8_MOUSE | *Ndufs8* |
| NDUV1_MOUSE | *Ndufv1* |
| NDUV2_MOUSE | *Ndufv2* |
| NEB2_MOUSE | *Ppp1r9b* |
| NECP1_MOUSE | *Necap1* |
| NECT1_MOUSE | *Nectin1* |
| NEGR1_MOUSE | *Negr1* |
| NEUM_MOUSE | *Gap43* |
| NFH_MOUSE | *Nefh* |
| NFL_MOUSE | *Nefl* |
| NFM_MOUSE | *Nefm* |
| NFS1_MOUSE | *Nfs1* |
| NFU1_MOUSE | *Nfu1* |
| NHRF1_MOUSE | *Slc9a3r1* |
| NIPS1_MOUSE | *Nipsnap1* |
| NIPS2_MOUSE | *Nipsnap2* |
| NLGN2_MOUSE | *Nlgn2* |
| NLGN3_MOUSE | *Nlgn3* |
| NMDE2_MOUSE | *Grin2b* |
| NMDZ1_MOUSE | *Grin1* |
| NNRD_MOUSE | *Naxd* |
| NNRE_MOUSE | *Naxe* |
| NOE1_MOUSE | *Olfm1* |
| NP1L1_MOUSE | *Nap1l1* |
| NP1L4_MOUSE | *Nap1l4* |
| NPTN_MOUSE | *Nptn* |
| NPTX1_MOUSE | *Nptx1* |
| NPTXR_MOUSE | *Nptxr* |
| NRX1A_MOUSE | *Nrxn1* |
| NRX1B_MOUSE | *Nrxn1* |
| NRX3B_MOUSE | *Nrxn3* |
| NSF1C_MOUSE | *Nsfl1c* |
| NT5C_MOUSE | *Nt5c* |
| NT5D3_MOUSE | *Nt5dc3* |
| NTRK2_MOUSE | *Ntrk2* |
| OCAD1_MOUSE | *Ociad1* |
| OCAD2_MOUSE | *Ociad2* |
| ODO2_MOUSE | *Dlst* |
| ODP2_MOUSE | *Dlat* |

| | |
|---|---|
| ODPA_MOUSE | *Pdha1* |
| ODPB_MOUSE | *Pdhb* |
| OMGP_MOUSE | *Omg* |
| OMP_MOUSE | *Omp* |
| OST48_MOUSE | *Ddost* |
| OTUB1_MOUSE | *Otub1* |
| OX2G_MOUSE | *Cd200* |
| PA1B2_MOUSE | *Pafah1b2* |
| PACN1_MOUSE | *Pacsin1* |
| PACS1_MOUSE | *Pacs1* |
| PAK1_MOUSE | *Pak1* |
| PALM_MOUSE | *Palm* |
| PARK7_MOUSE | *Park7* |
| PCBP1_MOUSE | *Pcbp1* |
| PCBP2_MOUSE | *Pcbp2* |
| PCCB_MOUSE | *Pccb* |
| PCLO_MOUSE | *Pclo* |
| PDCD6_MOUSE | *Pdcd6* |
| PDIA1_MOUSE | *P4hb* |
| PDIA3_MOUSE | *Pdia3* |
| PDIA6_MOUSE | *Pdia6* |
| PDXK_MOUSE | *Pdxk* |
| PEA15_MOUSE | *Pea15* |
| PEBP1_MOUSE | *Pebp1* |
| PFKAL_MOUSE | *Pfkl* |
| PFKAM_MOUSE | *Pfkm* |
| PFKAP_MOUSE | *Pfkp* |
| PGAM1_MOUSE | *Pgam1* |
| PGAM2_MOUSE | *Pgam2* |
| PGCB_MOUSE | *Bcan* |
| PGES2_MOUSE | *Ptges2* |
| PGK1_MOUSE | *Pgk1* |
| PGM2L_MOUSE | *Pgm2l1* |
| PGRC1_MOUSE | *Pgrmc1* |
| PHB_MOUSE | *Phb* |
| PHB2_MOUSE | *Phb2* |
| PHF24_MOUSE | *Phf24* |
| PHIPL_MOUSE | *Phyhipl* |
| PI51C_MOUSE | *Pip5k1c* |

| | |
|---|---|
| PIMT_MOUSE | *Pcmt1* |
| PIPNA_MOUSE | *Pitpna* |
| PKP4_MOUSE | *Pkp4* |
| PLEC_MOUSE | *Plec* |
| PP1A_MOUSE | *Ppp1ca* |
| PP1R7_MOUSE | *Ppp1r7* |
| PP2AB_MOUSE | *Ppp2cb* |
| PP2BB_MOUSE | *Ppp3cb* |
| PP2BC_MOUSE | *Ppp3cc* |
| PPIA_MOUSE | *Ppia* |
| PPIB_MOUSE | *Ppib* |
| PRDX1_MOUSE | *Prdx1* |
| PRDX2_MOUSE | *Prdx2* |
| PRDX3_MOUSE | *Prdx3* |
| PRDX5_MOUSE | *Prdx5* |
| PROF1_MOUSE | *Pfn1* |
| PROF2_MOUSE | *Pfn2* |
| PRRT2_MOUSE | *Prrt2* |
| PSA_MOUSE | *Npepps* |
| PSA1_MOUSE | *Psma1* |
| PSA5_MOUSE | *Psma5* |
| PSA7_MOUSE | *Psma7* |
| PSD3_MOUSE | *Psd3* |
| PTN11_MOUSE | *Ptpn11* |
| PTPR2_MOUSE | *Ptprn2* |
| PTPRZ_MOUSE | *Ptprz1* |
| PURA_MOUSE | *Pura* |
| PURB_MOUSE | *Purb* |
| PYC_MOUSE | *Pc* |
| PYGM_MOUSE | *Pygm* |
| QCR1_MOUSE | *Uqcrc1* |
| QCR2_MOUSE | *Uqcrc2* |
| QCR7_MOUSE | *Uqcrb* |
| QCR8_MOUSE | *Uqcrq* |
| RAB10_MOUSE | *Rab10* |
| RAB14_MOUSE | *Rab14* |
| RAB2A_MOUSE | *Rab2a* |
| RAB3A_MOUSE | *Rab3a* |
| RAB3C_MOUSE | *Rab3c* |

| | |
|---|---|
| RAB5B_MOUSE | *Rab5b* |
| RAB5C_MOUSE | *Rab5c* |
| RAB6A_MOUSE | *Rab6a* |
| RAC1_MOUSE | *Rac1* |
| RAC2_MOUSE | *Rac2* |
| RACK1_MOUSE | *Rack1* |
| RALA_MOUSE | *Rala* |
| RANG_MOUSE | *Ranbp1* |
| RAP2A_MOUSE | *Rap2a* |
| RASL1_MOUSE | *Rasal1* |
| RB11A_MOUSE | *Rab11a* |
| RB11B_MOUSE | *Rab11b* |
| RB27B_MOUSE | *Rab27b* |
| RCN2_MOUSE | *Rcn2* |
| RD23B_MOUSE | *Rad23b* |
| RGS6_MOUSE | *Rgs6* |
| RGS7_MOUSE | *Rgs7* |
| RHG01_MOUSE | *Arhgap1* |
| RHOA_MOUSE | *Rhoa* |
| RHOB_MOUSE | *Rhob* |
| RIDA_MOUSE | *Rida* |
| RILP_MOUSE | *Rilp* |
| RIMS1_MOUSE | *Rims1* |
| RL13_MOUSE | *Rpl13* |
| RLA2_MOUSE | *Rplp2* |
| RP3A_MOUSE | *Rph3a* |
| RS18_MOUSE | *Rps18* |
| RS19_MOUSE | *Rps19* |
| RS28_MOUSE | *Rps28* |
| RT23_MOUSE | *Mrps23* |
| RT36_MOUSE | *Mrps36* |
| RTN1_MOUSE | *Rtn1* |
| RTN3_MOUSE | *Rtn3* |
| RTN4_MOUSE | *Rtn4* |
| RUFY3_MOUSE | *Rufy3* |
| S12A5_MOUSE | *Slc12a5* |
| S4A10_MOUSE | *Slc4a10* |
| S4A4_MOUSE | *Slc4a4* |
| S4A8_MOUSE | *Slc4a8* |

| | |
|---|---|
| S6A11_MOUSE | *Slc6a11* |
| S6A17_MOUSE | *Slc6a17* |
| SAHH3_MOUSE | *Ahcyl2* |
| SAM50_MOUSE | *Samm50* |
| SC22B_MOUSE | *Sec22b* |
| SC6A6_MOUSE | *Slc6a6* |
| SCMC3_MOUSE | *Slc25a23* |
| SCN2A_MOUSE | *Scn2a* |
| SCN2B_MOUSE | *Scn2b* |
| SCOT1_MOUSE | *Oxct1* |
| SCPDL_MOUSE | *Sccpdh* |
| SCRN1_MOUSE | *Scrn1* |
| SDHA_MOUSE | *Sdha* |
| SEP11_MOUSE | *Septin11* |
| SEPT3_MOUSE | *Septin3* |
| SEPT4_MOUSE | *Septin4* |
| SEPT5_MOUSE | *Septin5* |
| SEPT6_MOUSE | *Septin6* |
| SEPT7_MOUSE | *Septin7* |
| SEPT9_MOUSE | *Septin9* |
| SERA_MOUSE | *Phgdh* |
| SERC_MOUSE | *Psat1* |
| SFXN5_MOUSE | *Sfxn5* |
| SGIP1_MOUSE | *Sgip1* |
| SGTA_MOUSE | *Sgta* |
| SH3G1_MOUSE | *Sh3gl1* |
| SH3G2_MOUSE | *Sh3gl2* |
| SHAN1_MOUSE | *Shank1* |
| SHAN2_MOUSE | *Shank2* |
| SHAN3_MOUSE | *Shank3* |
| SHLB2_MOUSE | *Sh3glb2* |
| SHPS1_MOUSE | *Sirpa* |
| SHSA7_MOUSE | *Shisa7* |
| SIR2_MOUSE | *Sirt2* |
| SKP1_MOUSE | *Skp1* |
| SLIRP_MOUSE | *Slirp* |
| SMCE1_MOUSE | *Smarce1* |
| SNAA_MOUSE | *Napa* |
| SNAB_MOUSE | *Napb* |

| | |
|---|---|
| SNAG_MOUSE | *Napg* |
| SNG1_MOUSE | *Syngr1* |
| SNG3_MOUSE | *Syngr3* |
| SNP23_MOUSE | *Snap23* |
| SNP25_MOUSE | *Snap25* |
| SNPH_MOUSE | *Snph* |
| SNX1_MOUSE | *Snx1* |
| SODC_MOUSE | *Sod1* |
| SODM_MOUSE | *Sod2* |
| SOGA3_MOUSE | *Soga3* |
| SPRE_MOUSE | *Spr* |
| SPTB1_MOUSE | *Sptb* |
| SPTB2_MOUSE | *Sptbn1* |
| SPTN1_MOUSE | *Sptan1* |
| SRC8_MOUSE | *Cttn* |
| SRCN1_MOUSE | *Srcin1* |
| SRR_MOUSE | *Srr* |
| SSBP_MOUSE | *Ssbp1* |
| STAM1_MOUSE | *Stam* |
| STIP1_MOUSE | *Stip1* |
| STMN1_MOUSE | *Stmn1* |
| STX12_MOUSE | *Stx12* |
| STX1A_MOUSE | *Stx1a* |
| STX1B_MOUSE | *Stx1b* |
| STX2_MOUSE | *Stx2* |
| SUCA_MOUSE | *Suclg1* |
| SUCB1_MOUSE | *Sucla2* |
| SV2A_MOUSE | *Sv2a* |
| SV2B_MOUSE | *Sv2b* |
| SYAC_MOUSE | *Aars* |
| SYBU_MOUSE | *Sybu* |
| SYGP1_MOUSE | *Syngap1* |
| SYN1_MOUSE | *Syn1* |
| SYN2_MOUSE | *Syn2* |
| SYN3_MOUSE | *Syn3* |
| SYNPO_MOUSE | *Synpo* |
| SYPH_MOUSE | *Syp* |
| SYSC_MOUSE | *Sars* |
| SYT1_MOUSE | *Syt1* |

| | |
|---|---|
| SYUB_MOUSE | *Sncb* |
| TAGL_MOUSE | *Tagln* |
| TAGL2_MOUSE | *Tagln2* |
| TAGL3_MOUSE | *Tagln3* |
| TALDO_MOUSE | *Taldo1* |
| TAU_MOUSE | *Mapt* |
| TBA4A_MOUSE | *Tuba4a* |
| TBB2A_MOUSE | *Tubb2a* |
| TBB3_MOUSE | *Tubb3* |
| TBB4A_MOUSE | *Tubb4a* |
| TBB4B_MOUSE | *Tubb4b* |
| TBB5_MOUSE | *Tubb5* |
| TBB6_MOUSE | *Tubb6* |
| TBC8B_MOUSE | *Tbc1d8b* |
| TCPA_MOUSE | *Tcp1* |
| TCPB_MOUSE | *Cct2* |
| TCPD_MOUSE | *Cct4* |
| TCPE_MOUSE | *Cct5* |
| TCPG_MOUSE | *Cct3* |
| TCPQ_MOUSE | *Cct8* |
| TCPW_MOUSE | *Cct6b* |
| TCPZ_MOUSE | *Cct6a* |
| TCTP_MOUSE | *Tpt1* |
| TENR_MOUSE | *Tnr* |
| THEM4_MOUSE | *Them4* |
| THIL_MOUSE | *Acat1* |
| THIM_MOUSE | *Acaa2* |
| THIO_MOUSE | *Txn* |
| THIOM_MOUSE | *Txn2* |
| THTM_MOUSE | *Mpst* |
| THTR_MOUSE | *Tst* |
| TIM44_MOUSE | *Timm44* |
| TIM50_MOUSE | *Timm50* |
| TKT_MOUSE | *Tkt* |
| TLN2_MOUSE | *Tln2* |
| TM1L2_MOUSE | *Tom1l2* |
| TMM65_MOUSE | *Tmem65* |
| TMOD2_MOUSE | *Tmod2* |
| TMOD3_MOUSE | *Tmod3* |

| | |
|---|---|
| TNNC2_MOUSE | *Tnnc2* |
| TOLIP_MOUSE | *Tollip* |
| TOM1_MOUSE | *Tom1* |
| TOM22_MOUSE | *Tomm22* |
| TOM70_MOUSE | *Tomm70* |
| TPD52_MOUSE | *Tpd52* |
| TPD54_MOUSE | *Tpd52l2* |
| TPIS_MOUSE | *Tpi1* |
| TPM2_MOUSE | *Tpm2* |
| TPM3_MOUSE | *Tpm3* |
| TPM4_MOUSE | *Tpm4* |
| TPPP_MOUSE | *Tppp* |
| TPRGL_MOUSE | *Tprg1l* |
| TRAP1_MOUSE | *Trap1* |
| TRFE_MOUSE | *Tf* |
| TTHY_MOUSE | *Ttr* |
| TTYH1_MOUSE | *Ttyh1* |
| TWF2_MOUSE | *Twf2* |
| TXTP_MOUSE | *Slc25a1* |
| UB2V1_MOUSE | *Ube2v1* |
| UBA1Y_MOUSE | *Uba1y* |
| UBE2N_MOUSE | *Ube2n* |
| UBP5_MOUSE | *Usp5* |
| UCHL1_MOUSE | *Uchl1* |
| UCHL3_MOUSE | *Uchl3* |
| UCHL4_MOUSE | *Uchl4* |
| UGPA_MOUSE | *Ugp2* |
| VAMP2_MOUSE | *Vamp2* |
| VAMP3_MOUSE | *Vamp3* |
| VAPA_MOUSE | *Vapa* |
| VAPB_MOUSE | *Vapb* |
| VAT1L_MOUSE | *Vat1l* |
| VATA_MOUSE | *Atp6v1a* |
| VATB2_MOUSE | *Atp6v1b2* |
| VATC1_MOUSE | *Atp6v1c1* |
| VATD_MOUSE | *Atp6v1d* |
| VATE2_MOUSE | *Atp6v1e2* |
| VATF_MOUSE | *Atp6v1f* |
| VATG2_MOUSE | *Atp6v1g2* |

| | |
|---|---|
| VATH_MOUSE | *Atp6v1h* |
| VDAC1_MOUSE | *Vdac1* |
| VDAC3_MOUSE | *Vdac3* |
| VGLU1_MOUSE | *Slc17a7* |
| VGLU3_MOUSE | *Slc17a8* |
| VINC_MOUSE | *Vcl* |
| VP26B_MOUSE | *Vps26b* |
| VPP1_MOUSE | *Atp6v0a1* |
| VPP2_MOUSE | *Atp6v0a2* |
| WASF1_MOUSE | *Wasf1* |
| WASF3_MOUSE | *Wasf3* |
| WBP2_MOUSE | *Wbp2* |
| WDR1_MOUSE | *Wdr1* |
| WDR7_MOUSE | *Wdr7* |
| YKT6_MOUSE | *Ykt6* |

**Table S5. Carbamidomethyl-modified peptides for high-confidence PPT1 substrates.** Protein localization and disulfide bond data were gathered from the UniProt entry for each protein. Carbamidomethyl (carba.) sites that were *not* validated in the tertiary screen are indicated by an asterisk (*). Disulfide bonds validated by a dNEM moiety in the tertiary screen are indicated (X). Mitochondrial membrane (MM), synaptic plasma membrane (SPM), GPi anchor (GPI).

| UniProt Accession | Peptide | Carba. site in peptide | Protein localization | Peptide site in protein | Peptide localization | Cysteine site | Disulfide Bond | dNEM |
|---|---|---|---|---|---|---|---|---|
| AT1A1_MOUSE | LIIVEG**C**QR | 7 | SPM | 699-707 | Cytoplasmic | 705 | | |
| AT1A2_MOUSE | LIIVEG**C**QR | 7 | SPM | 696-704 | Cytoplasmic | 702 | | |
| AT1A3_MOUSE | LIIVEG**C**QR | 7 | SPM | 689-697 | Cytoplasmic | 705 | | |
| AT1A3_MOUSE | VLGF**C**HYYLPEEQFPK | 5 | SPM | 542-557 | Cytoplasmic | 546 | | |
| AT1A3_MOUSE | YNTD**C**VQGLTHSK | 5 | SPM | 45-56 | Cytoplasmic | 49 | | |
| AT1B1_MOUSE | DSAQKDDMIFED**C**GNVPSEPK | 13 | SPM | 114-134 | Extracellular | 126 | to 149 | X |
| AT1B1_MOUSE | YNPNVLPVQ**C**TGK | 10 | SPM | 205-217 | Extracellular | 214 | to 277 | |
| AT1B2_MOUSE | TQLGD**C**SGIGDPTHYGYSTGQP**C**VFIK | 6<br>23 | SPM | 155-181 | Extracellular | 160<br>177 | to 177<br>to 160 | X |
| AT1B2_MOUSE | FLNVTPNVEVNVE**C**R | 14 | SPM | 248-262 | Extracellular | 261 | to 200 | X |
| ATPO_MOUSE | GEVP**C**TVTTASPLDDAVLSELK | 5 | MM | 137-158 | | 141 | | |
| BASI_MOUSE | SGEYS**C**IFLPEPVGR | 6 | SPM | 198-212 | Extracellular | 203 | to 157 | |
| CADM2_MOUSE | IIPSTPFPQEGQALTLT**C**ESK | 18 | SPM | 231-251 | Extracellular | 248 | to 296 | |
| CADM2_MOUSE | AYLTVLGVPEKPQISGFSSPVMEGDLMQLT**C**K | 31 | SPM | 116-147 | Extracellular | 146 | to 203 | |
| CD81_MOUSE | TFHETLN**CC**GSNALTTLTTTILR | 8<br>9 | SPM | 149-171 | Extracellular | 156<br>157 | to 190<br>to 175 | |
| CISD1_MOUSE | KFPF**C**DGAHIK | 5 | MM | 79-89 | | 83 | | |
| DYL2_MOUSE | NADMSEDMQQDAVD**C**ATQAMEK | 15 | Cytosolic | 10-31 | | 24 | | |
| GBB2_MOUSE | VS**C**LGVTDDGMAVATGSWDSFLK | 3 | Cytosolic | 315-337 | | 317 | | |
| GPM6A_MOUSE | KI**C**TASENFLR | 3 | SPM | 190-200 | Extracellular | 192 | to 174 | X |
| GRIA1_MOUSE | RGNAGD**C**LANPAVPWGQGIDIQR | 7 | SPM | 317-339 | Extracellular | 323 | to 75 | |
| KCRB_MOUSE | F**C**TGLTQIETLFK | 2 | Cytosolic | 253-265 | | 254 | | |
| LDHB_MOUSE | VIGSG**C**NLDSAR | 6 | Cytosolic | 159-170 | | 164 | | |
| LGI1_MOUSE | DFD**C**IITEFAK | 4 | Secreted | 218-228 | | 221 | | |
| NDKA_MOUSE | GDF**C**IQVGR | 4 | Nucleus | 106-114 | | 109 | | |
| NDUS1_MOUSE | M**C**LVEIEK | 2* | MM | 77-84 | | 78 | | |

| | | | | | | | | |
|---|---|---|---|---|---|---|---|---|
| NFASC_MOUSE | SGGRPEEYEGEYQ**C**FAR | 14 | SPM | 105-121 | Extracellular | 118 | to 63 | |
| NRCAM_MOUSE | DSTGTYT**C**VAR | 8* | SPM | 512-521 | Extracellular | 519 | to 470 | |
| NRCAM_MOUSE | TLQITHVSEADSGNYQ**C**IAK | 17 | SPM | 318-337 | Extracellular | 334 | to 286 | |
| NTRI_MOUSE | GTLQ**C**EASAVPSAEFQWFK | 5 | SPM-GPI | 239-257 | Extracellular | 243 | to 295 | |
| NTRI_MOUSE | EQSGEYE**C**SASNDVAAPVVR | 8 | SPM-GPI | 194-213 | | 201 | to 257 | |
| STXB1_MOUSE | AAHVFFTDS**C**PDALFNELVK | 10 | Cytosolic | 101-120 | | 110 | | |
| THY1_MOUSE | VTSLTA**C**LVNQNLR | 7 | SPM-GPI | 22-35 | Extracellular | 28 | to 131 | X |
| VDAC2_MOUSE | W**C**EYGLTFTEK | 2 | SPM | 76-86 | SPM | 77 | | |
| VDAC2_MOUSE | WNTDNTLGTEIAIEDQI**C**QGLK | 18 | SPM | 87-108 | SPM | 104 | | |
| VISL1_MOUSE | EFI**C**ALSITSR | 4* | Cytosolic | 84-94 | | 87 | | |
| VISL1_MOUSE | SDPSIVLLLQ**C**DIQK | 11 | Cytosolic | 177-191 | | 187 | | |
